# Cross-strain transferability of CRISPRi systems and design rules from laboratory to clinical *Escherichia coli* strains

**DOI:** 10.64898/2026.01.28.702340

**Authors:** Hyerim Ban, Stephen N. Rondthaler, Matthew Lebovich, Marcos A. Lora, Brandon Ugbesia, Lauren B. Andrews

**Affiliations:** Department of Chemical and Biomolecular Engineering, University of Massachusetts Amherst, Amherst, MA 01003 USA; Molecular and Cellular Biology Graduate Program, University of Massachusetts Amherst, Amherst, MA 01003 USA; Biotechnology Training Program, University of Massachusetts Amherst, Amherst, MA 01003 USA

**Author notes:** These authors contributed equally to this work and share first authorship (Hyerim Ban, Stephen N. Rondthaler).

## Abstract

CRISPR interference (CRISPRi) has emerged as a versatile approach for targeted gene repression in many organisms, including microbes and bacteria, due to the simple design of sequence-specific transcriptional silencing of gene expression. However, the strain-specific effects on repression efficiency and the host when translating a CRISPRi system from a laboratory strain to non-model strains are not well understood, yet they can present important limitations to its use. Here, we investigated the repression efficiency and toxicity of three CRISPRi systems (one dCas9 and two dCas12a variants) across four different *Escherichia coli* strains, including a laboratory K-12 strain (MG1655) and three non-model strains that are clinical isolates (probiotic Nissle 1917, uropathogenic CFT073, and uropathogenic UMN026). We evaluated the repression in each strain using sets of guide RNAs (gRNAs) targeting along the gene sequence and assayed cytotoxicity of expressing each dCas protein. Growth toxicity from expression of the different dCas proteins notably differed and showed high variation between some host strains. We also observed variable repression among the strains and notably poorer repression in multiple clinical strains. Therefore, we developed a dual gRNA CRISPRi system for enhanced gene silencing among the strains, which achieved up to 824-fold repression in CFT073. The results demonstrate that strain-specific design considerations can arise when a CRISPRi genetic system is transferred to a closely related bacterial strain. These findings provide insight into the relationships between criteria used for CRISPRi genetic design and in vivo activity across non-model *E. coli* strains, providing guidelines for diverse applications of these tools.

## Introduction

Synthetic biologists have established and employed many different genetic tools for bacteria, yet primarily this work is done in laboratory model strains, such as derivatives of *Escherichia coli* K-12.^1–4^ Recently, interest has turned to non-model strains and microorganisms for various applications,^5–7^ such as the study of pathogenic strains or the manipulation of strains with natural advantageous traits over common laboratory strains. However, understanding how the performance of genetic tools may differ when transferred to different strains of the same microbial species is limited. CRISPRi is one example of a popular genetic tool that is commonly used to regulate gene expression of a target gene in bacteria. This technology was first developed using specific class 2 CRISPR systems (e.g. CRISPR-Cas9 then CRISPR-Cas12a) and employ a deactivated CRISPR-associated Cas (dCas) protein that has been engineered to remove its nuclease activity.^8,9^ A guide RNA (gRNA) complexes with the dCas protein and directs binding to a target nucleic acid sequence via complementarity with the short spacer sequence (∼20 nucleotides) located on the gRNA. CRISPRi can repress expression by over 100-fold via steric hindrance of RNA polymerase to prevent transcription initiation or elongation.^8–10^ Due to its simplicity and strong repression, CRISPRi has been adopted for many different applications in diverse bacteria, including studying gene function,^11–16^ genome-wide mapping of genotypes to phenotypes,^3,16–23^ genetic circuits,^24–30^ and metabolic engineering.^30–36^

Many natural CRISPR-Cas systems spanning different classes, types, subtypes, and variants^37^ have been engineered for CRISPRi gene regulation. Of these, the CRISPRi technologies most commonly used in bacteria are derived from class 2 CRISPR-Cas and utilize a single dCas effector protein with mutations to deactivate the natural DNA nuclease activity. These include the widely used type II dCas9 from *Streptococcus pyogenes*^8^ and type V dCas12a (originally called dCpf1) from *Francisella novicida*,^10,38,39^ *Acidaminococcus* sp.,^9,40^ and *Lachnospiraceae bacterium.*^10,41^ CRISPRi using engineered Cas9 orthologs (e.g. *Staphylococcus aureus,*^43,44^ Streptococci^42,43^ and *Geobacillus thermodenitrificans*^44^) and Cas12a orthologs (e.g. *Eubacterium eligens*^45^) have also been established. Class 1 CRISPR-Cas systems, marked by their multi-subunit effector complexes composed of multiple Cas proteins, have also been modified for gene repression.^46–51^ However, they have typically been used in the endogenous bacterial species as transferability is challenging due to the large number of genes required. Natural CRISPR systems that target DNA often require at least one protospacer adjacent motif (PAM) sequence,^52,53^ and this requirement is specified as a design rule for a given CRISPRi system to repress a gene target.^8–10^ Although CRISPRi approaches that bind DNA for transcriptional repression have been more common, CRISPR systems that target RNA (e.g., type VI Cas13a,^54,55^ Cas13b,^56^ Cas13d,^57^ and type III Cas10^22^) have also been manipulated for gene repression and silencing in bacteria.

Given the breadth of CRISPRi systems available, a challenge that emerges is choosing and designing an optimal CRISPRi system for gene regulation in a specific bacterial host strain. Knowledge of how CRISPRi systems may function differently across strains of the same species (and different species) remains limited compared to the diversity of CRISPRi systems and bacterial strains available. Even for the most well-studied CRISPRi systems, the host-dependent effects on their gene regulation and the relative gene silencing for different CRISPRi systems are often unknown when transferring to a new strain. Few studies have quantitatively compared gene repression for different CRISPRi systems in the same bacterial strain,^42,45,58^ yet these works have shown that the common dCas9 from *Streptococcus pyogenes* is not efficacious in all bacteria. These studies have generally found differences in gene silencing efficiency and other characteristics (e.g., cytotoxicity of dCas expression) between different CRISPRi systems, including within the same subtype. While several studies have measured phenotypic changes with repression using CRISPRi in different bacterial species or strains,^44,59–65^ few have quantified and compared the repression in multiple strains.^51,59^ Despite the number of CRISPRi studies in *E. coli*, a comparison of CRISPRi systems in non-model strains has not been reported. Notably, previous work has identified significant strain-specific differences for gene expression, metabolic physiology, and functionality of genetic designs in different strains of *E. coli*. ^7,60,61^ A better understanding is needed to determine how the activity of a CRISPRi system may change in diverse strains and genetic backgrounds.

Here, we sought to investigate the strain-dependent performance of CRISPRi systems using laboratory and clinical *E. coli* strains, and we focus our study on a few of the most widely used class 2 CRISPRi systems (**Figure 1A**). Using inducible CRISPRi tools and a common genetic design, we systematically characterized gene repression for libraries of gRNAs and assayed the corresponding growth toxicity for each CRISPRi system in each strain (**Figure 1B**). We demonstrate that the level of repression and growth toxicity differed depending on the combination of CRISPRi system and host strain, although gRNA design rules were generally shared between the strains. We aimed to establish robust CRISPRi gene silencing in all three clinical strains, and to achieve this, we developed a multiplexed dual gRNA CRISPRi system. This work shows that strain-specific design (or re-design) of CRISPRi may be necessary for efficient gene silencing in a non-model bacterial strain to realize the potential transferability of CRISPRi gene regulation between bacterial strains.

**Figure 1.**
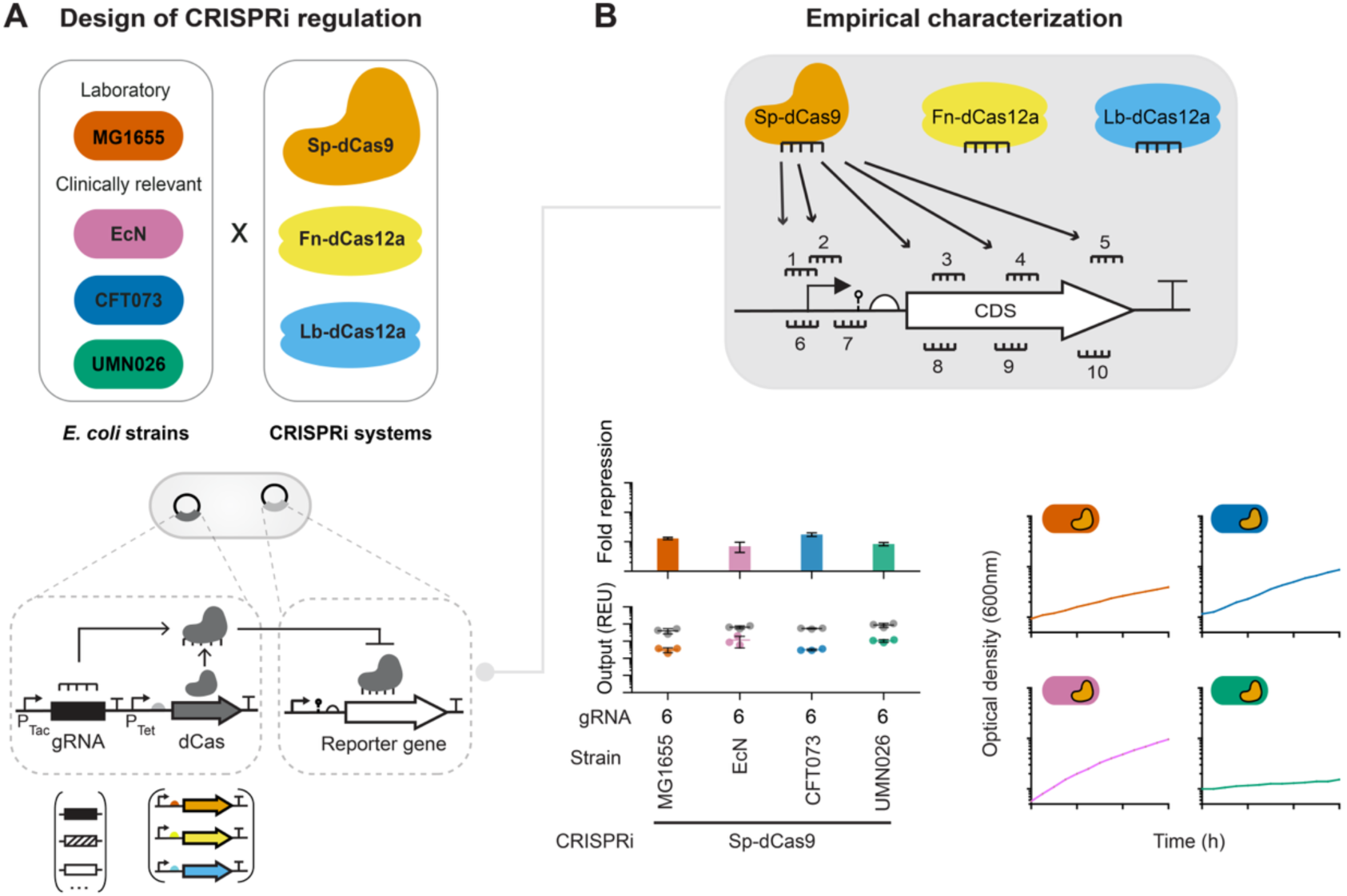
Overview of the study of class 2 CRISPRi systems within diverse *E. coli* strains. **(A)** The four *E. coli* strains include laboratory *E. coli* K-12 MG1655 (MG1655) and three clinical strains: commensal *E. coli* Nissle 1917 (EcN), uropathogenic *E. coli* CFT073, and uropathogenic *E. coli* UMN026. The three CRISPRi systems include dCas9 from *Streptococcus pyogenes* (Sp-dCas9), dCas12a from *Francisella novicida* U112 (Fn-dCas12a), and dCas12a from *Lachnospiraceae bacterium* ND2006 (Lb-dCas12a). Genetic schematic shows the two-plasmid system used to characterize the CRISPRi systems in *E. coli* strains. One plasmid expresses the deactivated Cas (dCas) protein and guide RNA (gRNA), each separately controlled by an inducible promoter. The second plasmid harbors the target gene, which is a fluorescent reporter. **(B)** To characterize each CRISPRi system, 10 gRNA were designed to bind along the upstream 5’ untranslated region and coding sequence (CDS) of the reporter gene, with 5 targeting each strand of DNA. Flow cytometry was used to measure fluorescence with and without expression of the gRNA to determine CRISPRi repression. Growth toxicity was determined using growth curve assays.

## Results and Discussion

### Characterizing CRISPRi systems in four strains of *E. coli*

We selected three popular class 2 CRISPRi systems in bacteria to quantify and compare their gene repression. CRISPRi using type II dCas9 from *Streptococcus pyogenes* (Sp-dCas9) is the first reported and one of the most commonly used in bacteria.^3,8,24,30,60^ To guide binding to the target DNA, Sp-dCas9 uses an engineered single guide RNA (sgRNA) comprised of a chimeric fusion of the natural tracrRNA and crRNA (**Figure 2A**).^61^ Sp-dCas9 requires a short NGG PAM sequence.^8,62^ However, Sp-dCas9 has been reported to be toxic in many species of bacteria ^17,24,43^ and cause changes in cell morphology,^63^ slower growth rate,^64–66^ and inability to transform plasmids containing Sp-dCas9.^67^ We also selected two type V dCas12a orthologs from *Francisella novicida* U112 (Fn-dCas12a) and *Lachnospiraceae bacterium* ND2006 (Lb-dCas12a),^10,38^ which have RNA processing activity of precursor crRNA that facilitates multiplex targeting.^9,10^ Unlike with Sp-dCas9, CRISPRi with dCas12a typically uses the natural crRNA structure as the gRNA to guide the CRISPRi complex, which involves a unique repeat sequence for each ortholog.^38^ Both Fn-dCas12a (**Figure 2B**) and Lb-dCas12a (**Figure 2C**) share the canonical AT-rich TTTV (V = A, G, C) PAM sequence, although they have different non-canonical PAM sequences.^10,68^ Differences in binding affinity have also been observed for Fn-dCas12a and Lb-dCas12a in biochemical assays.^69^ Additionally, dCas12a has been reported to be less toxic than Sp-dCas9 in several bacteria.^43,70–73^

**Figure 2.**
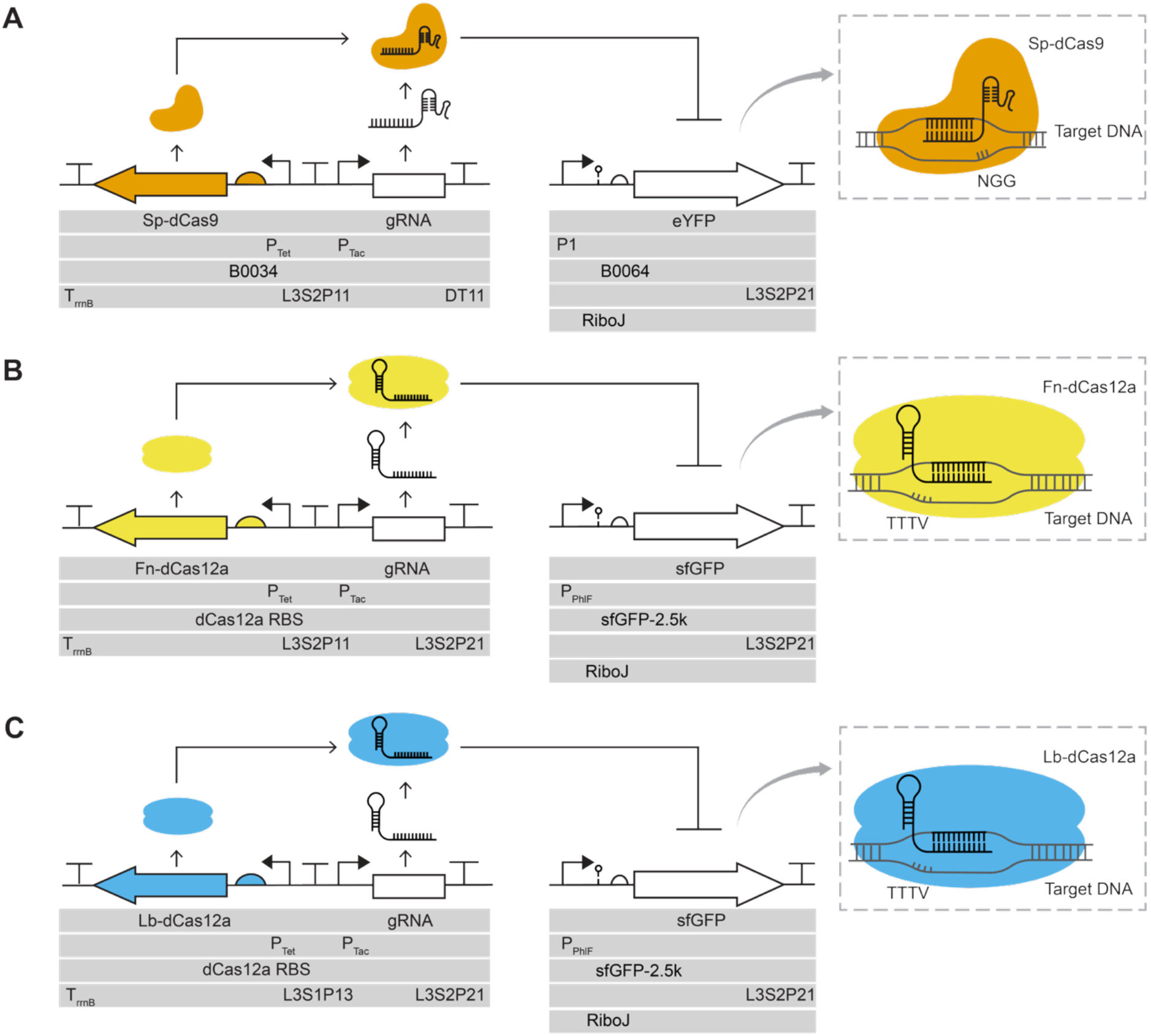
Genetic design for repression with each CRISPRi system. Inducible promoters (P_Tet_ and P_Tac_) regulate the expression of the dCas protein and gRNA, respectively, from a plasmid constructed for each dCas-gRNA combination. **(A)** The gRNA for the Sp-dCas9 system contains a 20 nt spacer sequence and a synthetic chimeric repeat sequence composed of the natural repeat sequences of the crRNA and tracrRNA. The dCas9-gRNA complex recognizes the NGG protospacer adjacent motif (PAM) to bind and repress the *eyfp* target gene expression from a constitutive promoter on the output plasmid. **(B)** Fn-dCas12a and **(C)** Lb-dCas12a gRNA contain a 20 nt spacer sequence but unique direct repeat sequences. Both dCas12a variants recognize the TTTV PAM to bind and repress the expression of the *sfgfp* target gene on the output plasmid. Genetic part sequences are in **Table S2**.

To explore CRISPRi efficiency when transferred to non-laboratory strains, four strains of *E. coli* were selected for this study. The common laboratory strain *E. coli* K-12 substr. MG1655 (MG1655) was included as the baseline, along with three strains that are clinical isolates: commensal *E. coli* Nissle 1917 (EcN), uropathogenic *E. coli* CFT073 (CFT073), and uropathogenic *E. coli* UMN026 (UMN026). These strains represent phylogenetic diversity of *E. coli* and are members of three phylogroups (**Figure S1** and **Table S1**). MG1655 from phylogroup A has a single chromosome of 4.64 Mb^72,73^ with no native plasmids. The non-model strains EcN^72,73^ and CFT073^72^ were selected from phylogroup B2 due to their high phylogenetic similarity yet contrasting relationships with the human host. EcN is a commensal strain that has been administered as a probiotic^73^ and has demonstrated potential as engineered living therapeutics,^74–77^ including approval for use in clinical trials.^78,79^ In contrast, CFT073 is a commonly studied uropathogenic *E. coli* (UPEC) strain that was isolated from the blood and urine of a woman with acute pyelonephritis.^80^ Both strains share a similar genome size (5.07 Mb for EcN^81,82^ and 5.23 Mb for CFT073^81^), although EcN contains two cryptic plasmids (pMUT1 and pMUT2) while CFT073 contains no native plasmids. Notably, *E. coli* is the predominant cause of urinary tract infections, which are among the most common infectious disease in the US.^82,83^ Therefore, a second UPEC strain UMN026^84^ was selected from phylogroup D. This strain is a urine isolate from a woman with acute cystitis. UMN026 has the largest genome of this set at 5.35 Mb ^85,86^ and contains two large (>33 kb) plasmids (p1ESCUM and p2ESCUM).^85^

Prior studies have not demonstrated and compared all three CRISPRi systems (Sp-dCas9, Fn-dCas12a, and Lb-dCas12a) in any of the four selected *E. coli* strains. CRISPRi using Sp-dCas9 has been well-established in MG1655 for many applications, including genetic circuits,^24,28,86^ metabolic engineering,^28,31–34^ investigating gene function,^87^ and genome-wide functional mapping.^3,17,21^ While CRISPRi using Fn-dCas12a has been reported in MG1655 for genetic circuits^86^ and RNA sensing,^29^ studies using Lb-dCas12a for CRISPRi in MG1655 have not been reported. In EcN, CRISPRi using Sp-dCas9 has been established (e.g., to study gene function^88,89^ and for metabolic engineering^90^) but has not been reported for either dCas12a variant. A couple studies have utilized CRISPRi in CFT073,^5,91^ yet only for Sp-dCas9, while CRISPRi has not previously been reported for UMN026 despite it being a reference strain for extraintestinal pathogenic *E. coli*.

We utilized a similar genetic design to characterize all CRISPRi systems in the *E. coli* strains. Inducible promoters were used to regulate expression of the dCas protein (P_Tet_) and gRNA (P_Tac_) on a low-copy plasmid (p15A replicon) (**Figure 2** and **Table S2**). Incorporating two inducible promoters can reduce leaky expression that causes unregulated repression, which has been observed in previous CRISPRi designs using a single inducible promoter.^19,92–94^ To limit transcription readthrough, the two transcriptional units were arranged in a divergent orientation with a strong terminator^95^ between them. The ribosomal binding site (RBS) for each dCas protein was designed to have similar predicted strength. To characterize the induction of each promoter, the response functions for P_Tet_ and P_Tac_ (associated with aTc- and IPTG-induced sensors, respectively) were assayed in each strain of *E. coli* using an insulated eYFP fluorescent reporter output from pAN1717^7,96^ (**Figures S2** and **S3**). The output is reported in relative promoter units (RPU) using the previously established RPU standard plasmid (pAN1717^7,96^), which generated similar fluorescence in the EcN, CFT073, and UMN026 *E. coli* strains and slightly greater fluorescence in MG1655 (**Figure S4**). Both P_Tet_ and P_Tac_ exhibited high induction (>800-fold for P_Tet_ and >300-fold for P_Tac_) and low basal activity in all strains (**Table S3**). For CRISPRi characterization, a low copy output plasmid (pSC101 replicon) containing the target reporter gene was used.

### Sp-dCas9 CRISPRi exhibited transferability between *E. coli* strains but differing repression and toxicity

First, the gene repression of Sp-dCas9 CRISPRi was assayed in each of the four *E. coli* strains. A library of ten gRNAs (sgRNAs) was designed to target sites along the *eyfp* output transcriptional unit with an adjacent PAM (NGG) (**Figure 3A**). gRNAs targeted the promoter region and coding sequence (CDS) on both the template and non-template DNA strands, as these design criteria have been shown to affect repression.^8–10,62,97^ Each *E. coli* strain was transformed with the output plasmid and CRISPRi plasmid for each gRNA, and single-cell fluorescence output was measured by flow cytometry with Sp-dCas9 and gRNA expression uninduced or induced at appropriate levels for each strain. The gRNA was fully induced in all strains while the induction of the dCas protein was chosen from preliminary titration experiments so that repression was observed and a sufficient number of cells was collected by flow (**Figure S5**). To compare the output between strains, fluorescence was converted to relative expression units (REU) by subtracting the strain’s autofluorescence of wildtype cells from measurements and dividing by the fluorescence of cells harboring the RPU standard plasmid (**Methods**). Fold repression was calculated as the observed dynamic range for regulated CRISPRi repression (i.e. ratio of output for the cells with dCas and gRNA uninduced to induced).

**Figure 3.**
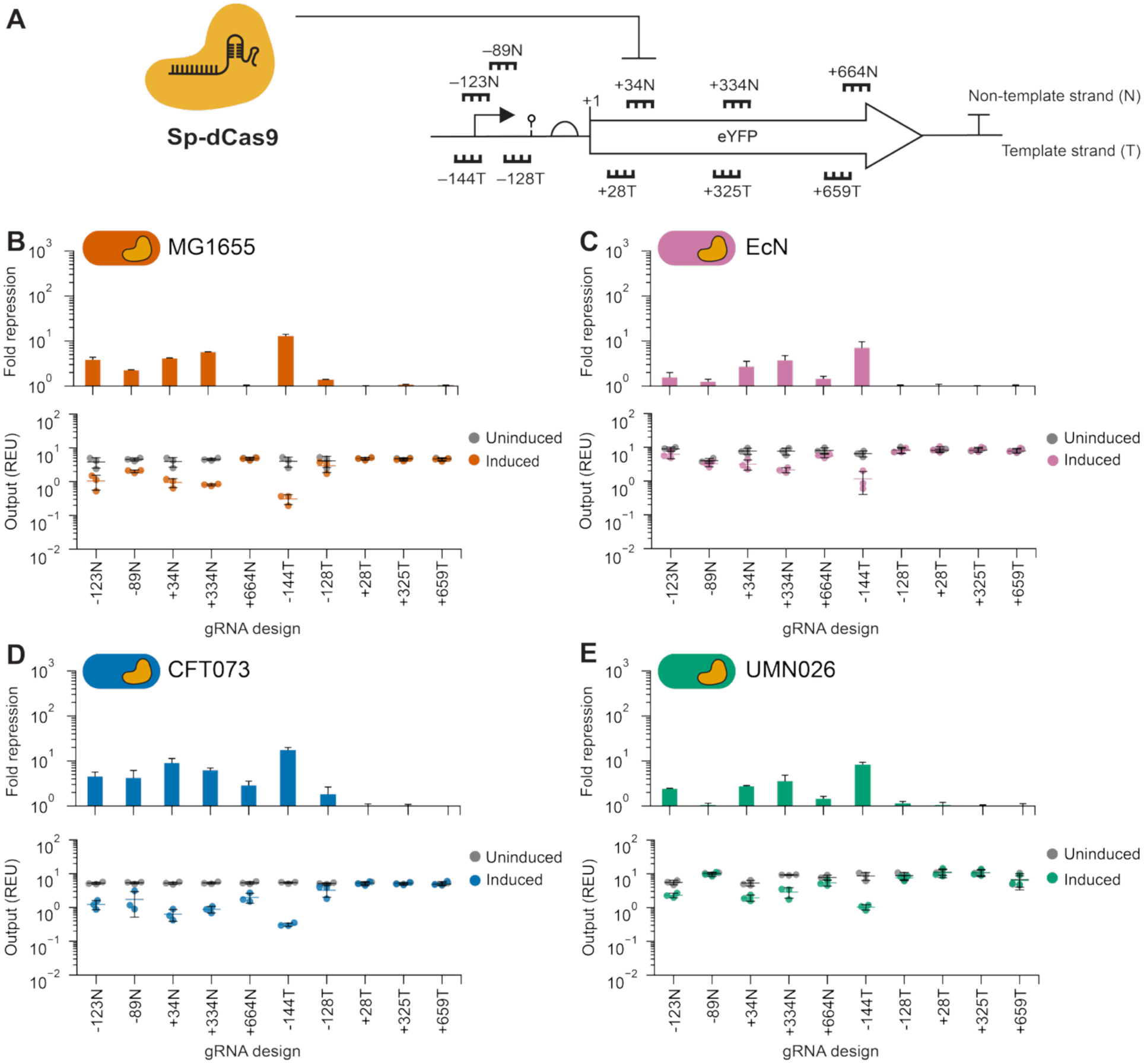
Sp-dCas9 CRISPRi repression in four diverse strains of *E. coli*. **(A)** Genetic schematic depicts the target DNA site of each gRNA design, named according to its binding location and target strand. gRNA names indicate the location relative to the start of the *eyfp* coding sequence (+1) and whether it targets the non-template (N) or template (T) strand of DNA. For each gRNA design, the output of the eYFP reporter was measured in *E. coli* **(B)** MG1655, **(C)** EcN, **(D)** CFT073, and **(E)** UMN026 by flow cytometry with uninduced (gray) or induced (colored) expression of dCas9 and gRNA, using aTc concentrations of 0.25 ng/mL for MG1655, CFT073, and UMN026 and 0.125 ng/mL for EcN. Fluorescent output was converted to relative expression units (REU) in each strain (**Methods**). Fold repression was calculated as the ratio of the output without and with induction. Error bars indicate the standard deviation of three identical experiments performed on separate days. Fluorescence histograms are in **Figures S35-S38**.

Strong correlations between Sp-dCas9 CRISPRi repression for the gRNAs across the strains were observed despite small differences in output REU values (**Figures 3** and **S6**). In MG1655, the output was a narrow range without CRISPRi induction (3.8 – 4.8 REU), while induction generated repression up to 13-fold (**Figure 3B**). EcN yielded greater output without induction (4.0 – 8.9 REU) and less repression, up to only 7.0-fold (**Figure 3C**). Interestingly, the results in CFT073 were more similar to those of MG1655 than EcN, exhibiting moderate output without induction (4.9 – 5.4 REU) and repression up to 17.6-fold (**Figure 3D**). Sp-dCas9 CRISPRi yielded the greatest output when uninduced in UMN026 (5.3 – 11.1 REU) and produced repression up to 8.3-fold (**Figure 3E**). The fold repression for 16.7% (10/60) of all combinations of gRNA and strain pairs was significantly different, involving five gRNA (–144T, –123N, –89N, +34N, and +664N) (one-way ANOVA with Tukey post-hoc analysis, *p* = 0.0010 – 0.036, **Table S4**). Of these differing gRNA-strain pairs, all but one of these combinations included CFT073 (9/10), and half (5/10) were between CFT073 and EcN, due to greater repression of active gRNA for Sp-dCas9 in CFT073 compared to other strains. Despite these differences, correlations in fold repression for all gRNA between each pair of strains were strong (Pearson’s *r* = 0.889 – 0.975, **Figure S6**), suggesting common gRNA design rules. Overall, modest repression was achieved, and these results demonstrate transferability of gRNA designs and Sp-dCas9 CRISPRi repression between the *E. coli* strains, although with notable differences in the repressed output between strains.

Toxicity of Sp-dCas9 expression was assayed in each *E. coli* strain by measuring the growth kinetics for a range of induction via aTc inducer concentration (**Figures 4** and **S7**). With Sp-dCas9 expression, growth toxicity was observed in all strains, yet to varying extents. At high induction of Sp-dCas9 (2 or 4 ng/mL aTc), all four strains showed significant reduction in OD_600_ at 4 hours compared to the respective wildtype strain (Welch’s one-sided, unpaired *t*-test, *p* < 0.05), ranging from 30% (EcN at 2 ng/mL aTc) to 94% (UMN026 at 4 ng/mL aTc) reduction (**Figure 4A**). All strains except EcN exhibited significantly reduced specific growth rate with high induction of Sp-dCas9 expression, ranging from 32% (CFT073 at 2 ng/mL) to 72% (UMN026 at 4 ng/mL) reduced growth rate (**Figure 4B**). Low induction of Sp-dCas9 expression (0.25 ng/mL aTc) resulted in significant toxicity for UMN026 but not the other *E. coli* strains. Without induction, UMN026 also displayed a significant decrease in cell growth (46% lower OD_600_ at 4 h, *p* = 0.0010) and growth rate (12%, *p* = 0.021). Neither the strains’ differences in P_Tet_ promoter output (**Figure S8**) nor their wildtype growth correlate with the observed strain-specific toxicity of Sp-dCas9, suggesting other factors contribute to the uniquely high toxicity in UMN026. Taken together, these growth data illustrate strain-specific differences in growth toxicity that can arise, highlighting the importance of assessing Sp-dCas9 toxicity when introducing the CRISPRi system to a new bacterial strain.

**Figure 4.**
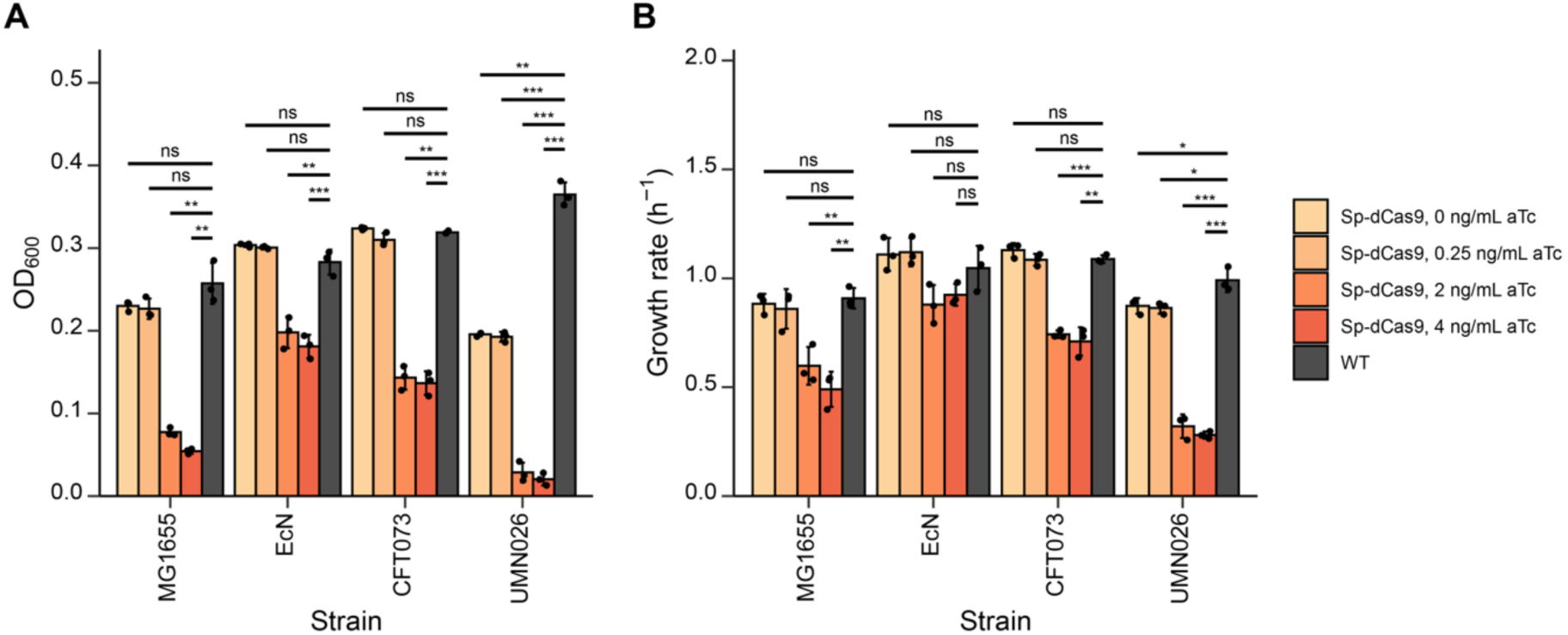
Assaying growth toxicity of Sp-dCas9 expression in *E. coli* strains. Sp-dCas9 expression from the CRISPRi plasmid in four *E. coli* strains was uninduced or induced with 0.25, 2, or 4 ng/mL aTc (orange). Wildtype (WT) cells for each strain (gray), which did not contain the CRISPRi plasmid, were assayed for comparison. The OD_600_ was measured every 15 minutes over an 8-hour incubation using a microplate reader (**Figure S8**). Growth between each induction level of Sp-dCas9 and wildtype for each strain is compared for **(A)** the OD_600_ at four hours of incubation and **(B)** the specific growth rate fitted to the growth curve. Experiments were performed in triplicate on three different days. Mean ± s.d. (*n* = 3) is plotted. Results of Welch’s one-sided, unpaired *t*-tests are indicated as not significant (ns) or significant (\**p* < 0.05, \*\**p* < 0.01, and \*\*\**p* < 0.001).

### Fn-dCas12a CRISPRi demonstrated stronger repression and lower toxicity than Sp-dCas9

Next, we investigated Fn-dCas12a CRISPRi activity in the four *E. coli* strains. Given the different canonical PAM sequence of dCas12a (TTTV) compared to dCas9 (NGG) and limited potential target sites in the *eyfp* output gene, an *sfgfp* output was used to assay the dCas12a CRISPRi systems. Ten gRNA were designed to target either DNA strand of the promoter region and along the CDS (**Figure 5A**). Repression and growth toxicity were then assayed for Fn-dCas12a CRISPRi in the strains as previously described for Sp-dCas9.

**Figure 5.**
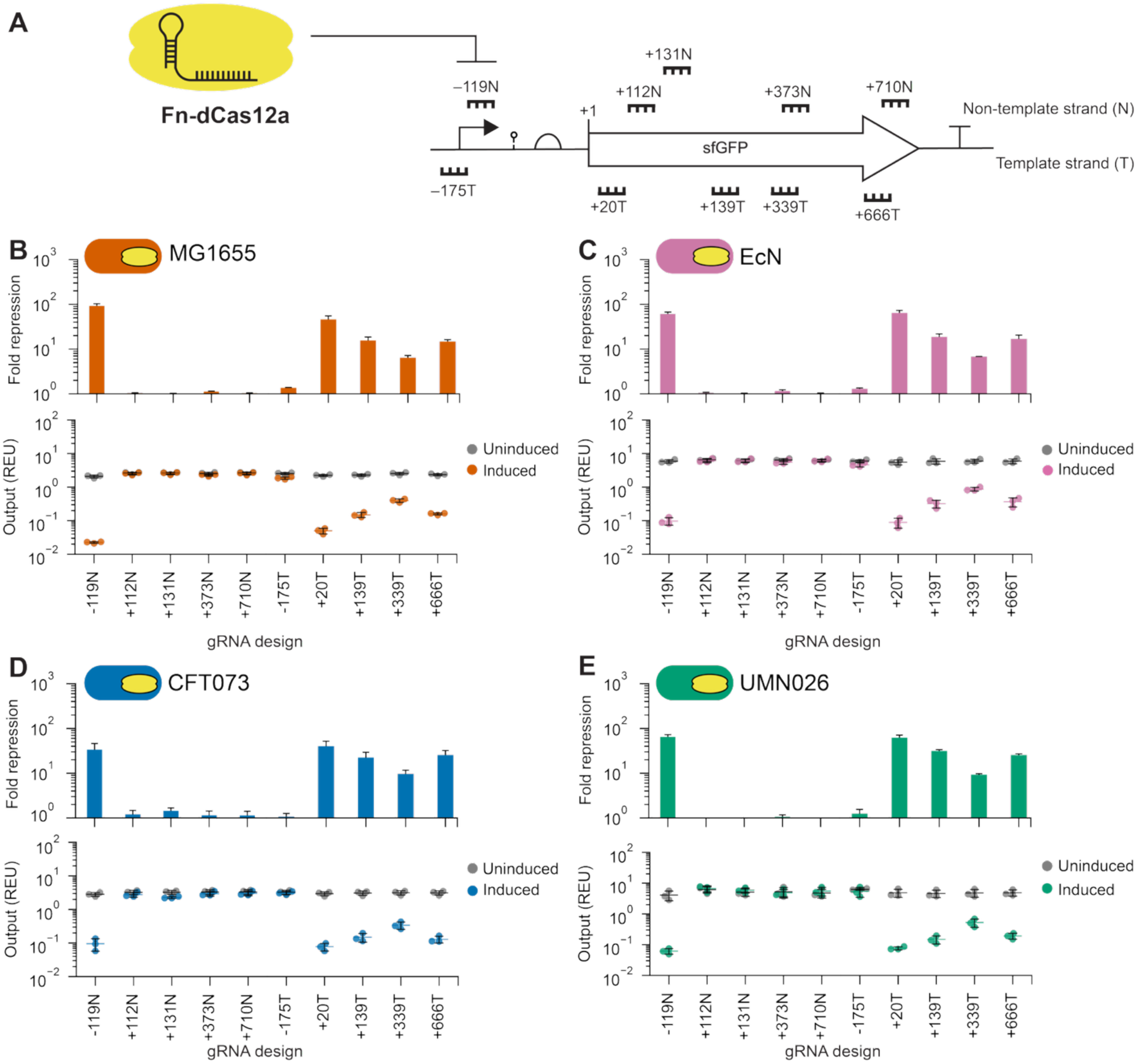
Characterization of Fn-dCas12a CRISPRi repression in *E. coli* strains. **(A)** DNA target sites of gRNA designed for the dCas12a systems are shown in the genetic schematic. gRNA names indicate the targeted location relative to the start of the *sfgfp* coding sequence (+1) and whether the non-template (N) or template (T) DNA strand is targeted. For each gRNA, the fluorescence output was measured in **(B)** MG1655, **(C)** EcN, **(D)** CFT073, and **(E)** UMN026 by flow cytometry. Expression of Fn-dCas12a and the gRNA was uninduced (gray) or induced with aTc (0.25 ng/mL for MG1655, 0.125 ng/mL for EcN, and 0.5 ng/mL for CFT073 and UMN026) and IPTG (1 mM for all strains) (colored). Fluorescence was converted to relative expression units (REU) for each strain (Methods). Fold repression was calculated as the ratio of the output without and with induction. Error bars are the standard deviation of three identical experiments performed on separate days. Fluorescence histograms are in **Figures S39-S42**.

Strong repression was observed in all strains using Fn-dCas12a CRISPRi (**Figure 5**). MG1655 demonstrated the lowest uninduced output (2.1 – 2.6 REU) but achieved the greatest repression (up to 93-fold) (**Figure 5B**). In EcN, greater output was observed without induction (5.5 – 6.4 REU) and repression reached up to 65-fold (**Figure 5C**). CFT073 presented relatively lower uninduced output (2.8 – 3.3 REU) and the least repression among the strains (up to 40-fold) (**Figure 5D**). UMN026 showed similar results to EcN in both uninduced output (4.1 – 7.0 REU) and magnitude of repression (up to 65-fold) (**Figure 5E**). While 58% (35/60) of gRNA and strain pair combinations showed significantly different uninduced output (*p* = 9.7 × 10^-6^ – 0.045), only 8.3% (5/60) exhibited different repression between strains (*p* = 0.0059 – 0.048) (**Table S4**). All five of these combinations contained CFT073, further illustrating strain-specific differences in CRISPRi activity. The maximum CRISPRi repression observed using Fn-dCas12a was significantly greater than Sp-dCas9 in all strains (one-way ANOVA with Tukey post-hoc, *p* = 4.6 × 10^-6^ – 0.038, **Figure S9**). Additionally, the repression for gRNA targeting the preferred DNA strand within the CDS was greater for Fn-dCas12a (template^9,10^) than that for Sp-dCas9 (non-template^8^) (*p* = 3.3 × 10^-9^ – 4.4 × 10^-5^) (**Figure S10**). Correlations in repression between strains were also slightly stronger for CRISPRi with Fn-dCas12a (*r* = 0.967 – 0.994, **Figure S11**) than Sp-dCas9. Altogether, these results demonstrate strong repression achieved for Fn-dCas12a CRISPRi across all *E. coli* strains with fewer differences between strains compared to Sp-dCas9.

Growth assays revealed significant toxicity for Fn-dCas12a expression in all strains (**Figures 6** and **S12**). With high induction of Fn-dCas12a expression (2 and 4 ng/mL aTc), all four *E. coli* strains showed reduced growth relative to their respective wildtype strain (*p* = 2.2 × 10^-5^ – 0.030) (**Figure 6A**). Growth reductions ranged from 33% for EcN (2 ng/mL aTc) up to 84% for MG1655 (4 ng/mL aTc), which exhibited the greatest apparent toxicity. Similarly, all four strains exhibited decreased specific growth rates (*p* < 0.05) with high induction, ranging from 17% (EcN at 2 ng/mL) to 57% (MG1655 at 4 ng/mL) (**Figure 6B**). Low induction of Fn-dCas12a did not significantly reduce growth in any strain. For UMN026, growth was statistically greater with Fn-dCas12a than with Sp-dCas9 expression at all aTc concentrations (one-way ANOVA with Tukey post-hoc analysis, *p* = 2.1 × 10^-6^ – 0.015), but not for other strains (**Figure S13**). Prior studies have reported less toxicity for Fn-dCas12a relative to Sp-dCas9 in MG1655 and some other bacteria.^71,86,98^ However, we observed that CRISPRi-related growth effects can vary markedly across *E. coli* strains.

**Figure 6.**
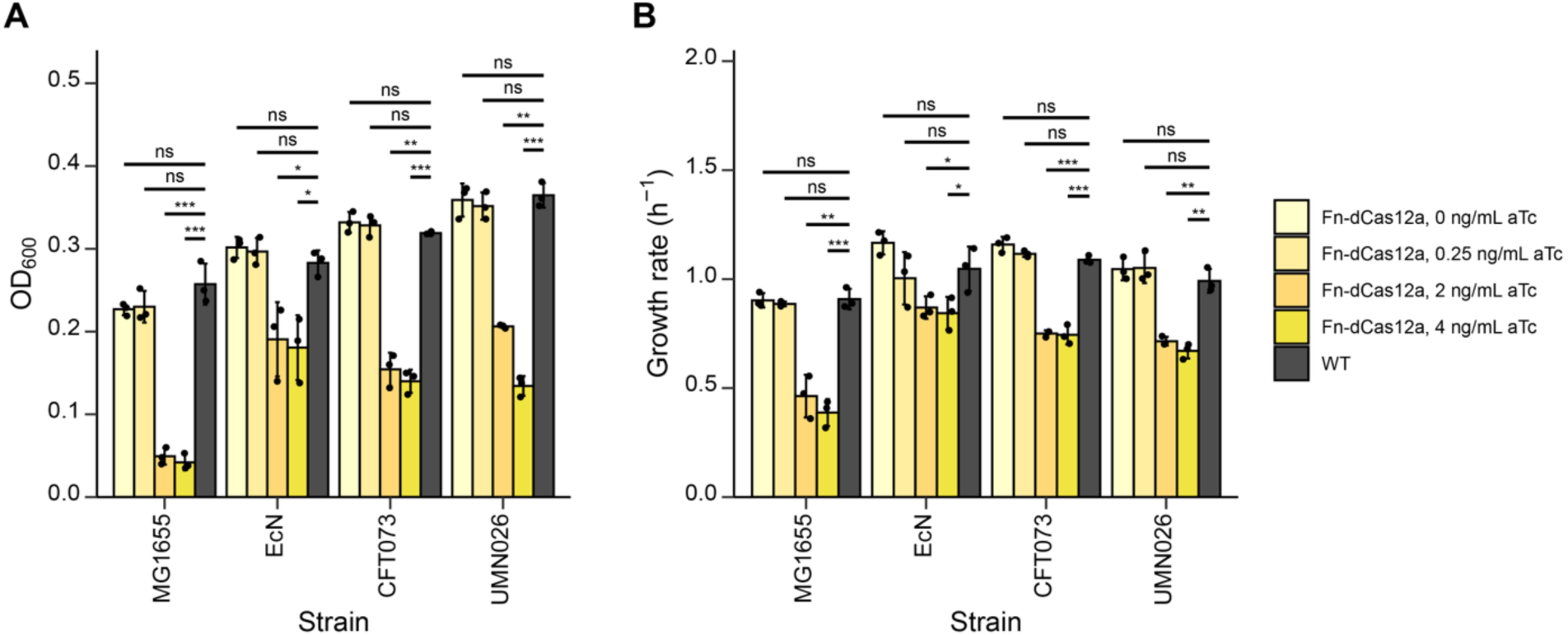
Toxicity of Fn-dCas12a expression assayed in *E. coli* strains. The expression of Fn-dCas12a was induced in four strains of *E. coli* using 0, 0.25, 2, or 4 ng/mL aTc (yellow). The wildtype cells (WT, gray) of each strain were assayed without aTc for comparison. Cell growth was monitored by measuring OD_600_ every 15 minutes over an 8-hour incubation (**Figure S12**). **(A)** Optical density at 600 nm (OD_600_) at four hours of incubation and **(B)** the fitted specific growth rate are plotted. All experiments were performed in triplicate on three different days (mean ± s.d.). Results of statistical analysis are shown as not significant (ns) or significant (\**p* < 0.05, \*\**p* < 0.01, and \*\*\**p* < 0.001).

### Lb-dCas12a CRISPRi displayed repression in all *E. coli* strains and lowest growth toxicity

As Lb-dCas12a shares the conical TTTV PAM sequence of Cas12a orthologs,^10,68^ the same set of gRNA spacer designs and output plasmid were used to characterize Lb-dCas12a CRISPRi in the four strains of *E. coli* as used with Fn-dCas12a (**Figure 7A**). The direct repeat sequence for Lb-dCas12a was incorporated into the gRNA designs. A new set of CRISPRi plasmids was constructed to express Lb-dCas12a and each gRNA. Each CRISPRi plasmid was transformed with the output plasmid into each *E. coli* strain for assaying its repression.

**Figure 7.**
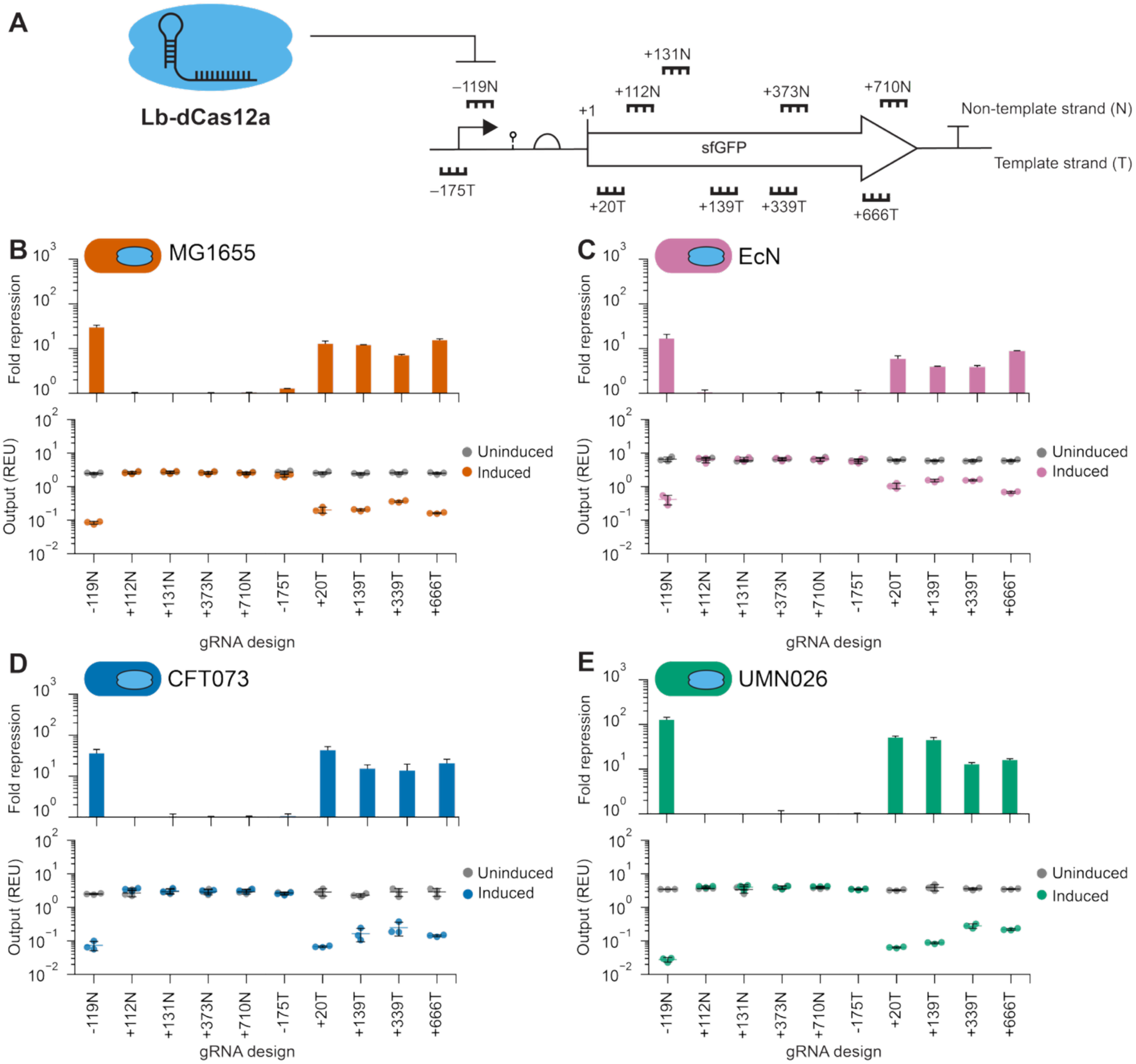
Characterization of Lb-dCas12a CRISPRi repression in *E. coli* strains. **(A)** Genetic map depicting the DNA target sites of gRNAs used for assaying Lb-dCas12a CRISPRi repression. The *sfgfp* reporter gene output was measured in **(B)** MG1655, **(C)** EcN, **(D)** CFT073, and **(E)** UMN026 for each of the 10 gRNA using flow cytometry with Lb-dCas12a and the gRNA expression uninduced (gray) or induced with aTc (0.25 ng/mL for MG1655, 0.125 ng/mL for EcN, and 2 ng/mL for CFT073 and UMN026) and IPTG (1 mM for all strains) (colored). Output in relative expression units (REU) was calculated for each strain and gRNA (**Methods**). Fold repression is the ratio of the output uninduced to induced. Mean and standard deviation of three identical experiments performed on separate days are plotted. Fluorescence histograms are in **Figures S43-S47**.

CRISPRi repression was observed in all *E. coli* strains using Lb-dCas12a (**Figure 7**). High output expression was observed for all strains when Lb-dCas12a CRISPRi was not induced. However, the magnitude of repression among the strains notably differed. For MG1655 and EcN, Lb-dCas12a CRISPRi achieved only moderate repression (up to 30-fold in MG1655 and 17-fold in EcN), which is substantially less than Fn-dCas12a repression for the maximal activity gRNA (*p* < 0.001 for MG1655; *p* < 0.05 for EcN) (**Figure S9**). For the UPEC strains, silencing with Lb-Cas12a was strong in CFT073 (up to 43-fold) and very strong in UMN026 (up to 130-fold) (**Figures 7D** and **7E**). Compared to Fn-dCas12a, CRISPRi with Lb-dCas12a achieved similar maximal repression in CFT073 (*p* = 0.94) but stronger repression in UMN026 (*p* < 0.01). These results illustrate important differences in repression efficiency between even highly related CRISPRi systems transferred to different bacterial strains. Despite greater differences in the magnitude of repression observed between strains for the Lb-dCas12a system, repression among the library of gRNAs were strongly correlated between all strain pairs (Pearson’s *r* = 0.947 – 0.983) (**Figure S14**).

Growth assays were performed for Lb-Cas12a expression in the four *E. coli* strains (**Figure S15**). Reduced cell growth was only observed with high induction of Lb-dCas12a expression (2 or 4 ng/mL aTc) for all strains (*p* < 0.05) (**Figure 8A**). Significant reduction in growth rate (*p* < 0.05) was observed for only MG1655, CFT073, and UMN026 with high induction (**Figure 8B**), ranging from 14% (UMN026 at 4 ng/mL) to 47% (MG1655 at 4 ng/mL). Specific growth was significantly greater in UMN026 with high induction of Lb-dCas12a compared to Fn-dCas12a (2 and 4 ng/mL aTc with *p* = 0.0010 and 2.1 × 10^-4^, respectively) but comparable for the other strains (**Figure S13**). These results show notable differences in toxicity observed for dCas expression of different orthologs and between *E. coli* strains. Overall, Lb-dCas12a exhibited the least growth toxicity of the dCas proteins tested in UMN026 and comparable toxicity in the other three strains of *E. coli*.

**Figure 8.**
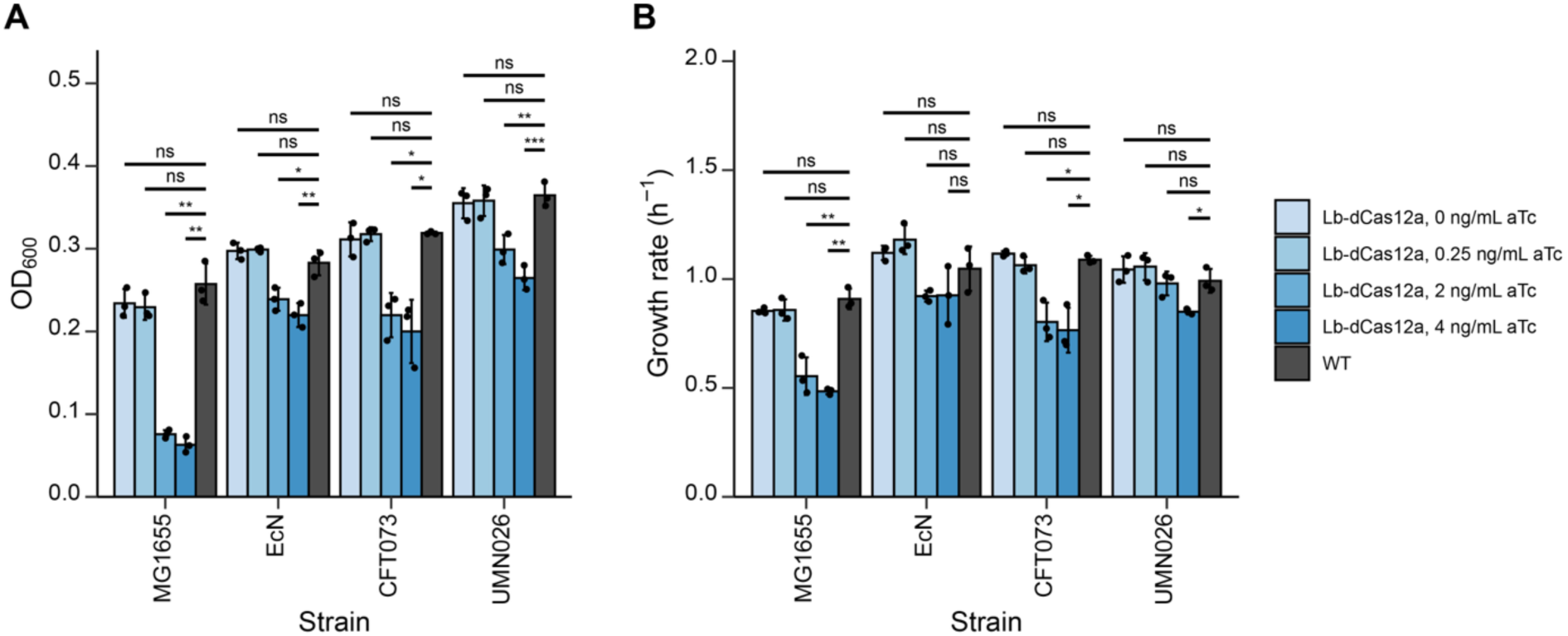
Toxicity of Lb-dCas12a expression assayed in *E. coli* strains. Expression of Lb-dCas12a in each *E. coli* strain was induced using 0, 0.25, 2, or 4 ng/mL aTc (blue). The wildtype cells (WT, gray) of each strain were assayed without aTc for comparison. Cell growth was monitored by measuring optical density (OD_600_) every 15 minutes over an 8-hour incubation (**Figure S15**). **(A)** The OD_600_ at four hours of growth and **(B)** specific growth rate were determined to compare growth in each condition. All experiments were performed in triplicate on three separate days. Statistical analysis determined whether results are not significant (ns) or significant (\**p* < 0.05, \*\**p* < 0.01, and \*\*\**p* < 0.001).

### Development of a new CRISPRi system with Lb-dCas12a and dual gRNAs for enhanced repression and strong gene silencing across all *E. coli* strains

Next, we sought to develop a CRISPRi system that could achieve strong gene silencing across all strains with a simple strategy for gRNA design and relatively little toxicity. The results thus far revealed a dependency on both the CRISPRi system and *E. coli* strain for the repression efficiency observed for a target gene. Additionally, the observed repression did not achieve very strong gene silencing (> 100-fold repression) in all strains. Previous studies have demonstrated that multiple gRNA targeting a single gene can increase CRISPRi repression in bacteria for both Sp-dCas9^8^ and Fn-dCas12a.^99,100^ Thus, we hypothesized that targeting the output gene with two gRNA using Lb-dCas12a CRISPRi could similarly increase repression. For this approach, we chose Lb-dCas12a because it (i) showed the least growth toxicity of the dCas proteins tested, (ii) exhibited moderate to strong repression in all strains, and (iii) is a Cas12a ortholog that can process pre-crRNA to facilitate multiplex gRNA arrays.^9,99,101^ Additionally, we aimed to target only the gene’s CDS to make this approach easier to employ for non-model strains with less thoroughly annotated genomes. Therefore, we selected four gRNA designs targeting the preferred DNA strand (template strand) within the CDS to create dual gRNA CRISPR arrays to test this approach. We constructed a library of 12 CRISPRi plasmids with all combinations of the gRNAs in CRISPR arrays to characterize CRISPRi repression. Both gRNAs were contained in one transcriptional unit regulated by the P_Tac_ promoter with an identical direct repeat sequence in each gRNA that targeted a different site in *sfgfp* via two separate dCas12a-gRNA complexes (**Figure 9A**).

**Figure 9.**
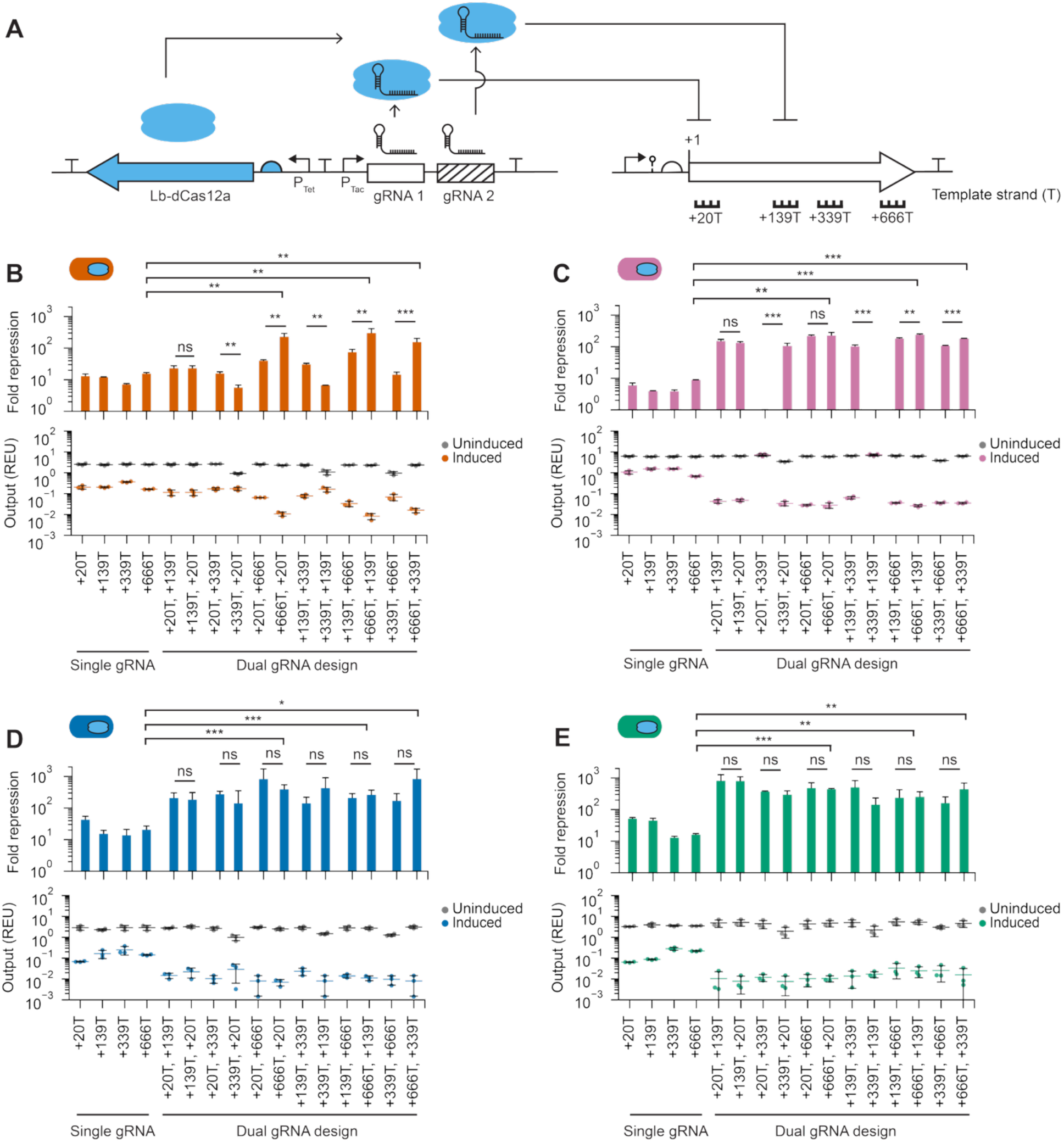
CRISPRi repression using Lb-dCas12a with dual gRNA arrays in *E. coli* strains. **(A)** The genetic design of the dual gRNA system for Lb-dCas12a uses two gRNAs in a CRISPR array to bind two different sites in the *sfgfp* target gene. Binding locations of the four selected gRNA are depicted. Dual gRNA CRISPR arrays were created for all 12 combinations of gRNAs with the 5′ to 3′ position of each gRNA denoted by the ordering in the name. The output of the *sfgfp* reporter gene was measured in **(B)** MG1655 (orange), **(C)** EcN (pink), **(D)** CFT073 (blue), and **(E)** UMN026 (green) for each CRISPRi plasmid with a dual gRNA or single gRNA using flow cytometry with Lb-dCas12a and the gRNA array uninduced (gray) or induced (colored). Error bars indicate the standard deviation of three identical experiments performed on separate days. Statistical analysis of repression for dual gRNA versus single gRNA (+666T shown) was determined using Welch’s one-sided, unpaired *t*-tests, with the results for all combinations given in **Figure S16**. Statistical analysis of repression comparing both orderings of the gRNA in the dual gRNA arrays used a Welch’s two-sided, unpaired *t*-test. Results are shown as not significant (ns) or significant (\**p* < 0.05, \*\**p* < 0.01, and \*\*\**p* < 0.001). Corresponding fluorescence histograms are in **Figures S48-S50**.

In all strains, dual gRNA designs for Lb-dCas12a were able to surpass repression of the single gRNAs, with some designs for each strain achieving very strong (>100-fold) silencing (**Figure 9**). Repression for most dual gRNA designs was significantly greater than the single gRNA constituents, with 66% of dual gRNA arrays (8/12) for MG1655, 83% (10/12) for EcN, 91% (11/12) for CFT073, and 100% (12/12) for UMN026 of the dual gRNA designs showing improved repression (*p* = 1.5 × 10^-8^ – 0.047) (**Figures 9** and **S16**). In the UPEC strains, CRISPRi repression was exceptionally strong for the dual gRNA designs (average of 340-fold in CFT073 and 410-fold in UMN026) and exceeded gene silencing observed in MG1655 (77-fold average) and EcN (140-fold average). Additionally, strong gene silencing in the UPEC strains was not significantly affected by the order of the gRNA in the CRISPR array (**Figures 9D** and **9E**). However, notable differences in repression were observed when the positions in the array were swapped for many gRNA pairs in MG1655 or EcN (**Figures 9B** and **9C**). Unexpectedly, swapping the arrangement for two gRNA combinations in EcN (+20T with +339T and +339T with +139T) resulted in a loss of detectable silencing, which was reproducible after retransforming and assaying sequence-verified plasmids. When comparing the silencing between strains, the repression for each dual gRNA design was not strongly correlated between the strains, except for MG1655 and EcN (Spearman’s *ρ* = 0.86) (**Figure S17**). Lb-dCas12a CRISPRi using dual gRNA arrays demonstrated efficient repression for most designs yet also drastic strain-specific differences.

Dual gRNA designs having the greatest gene silencing showed high output without induction, demonstrating tight gene regulation using this CRISPRi system (**Figure 9**). However, interestingly, three dual gRNA arrays (+339T with +20T, +339T with +666T, and +339T with +139T) displayed decreased output without induction, which was a consistent pattern observed across all four strains. All three arrays share the same gRNA in the first position (+339T), yet leaky repression was not observed for this gRNA individually.

Combined, these results provide evidence that dual gRNA arrays with Lb-dCas12a CRISPRi can substantially increase repression and achieve strong gene silencing in all strains. However, the relative effect on repression can be dependent on the strain and, for some strains, the arrangement of gRNAs within the CRISPR array, emphasizing the need to characterize such designs in new strains. Notably, dual gRNAs with Lb-dCas12a achieved the strongest CRISPRi silencing in all three clinical strains in this study, providing a powerful tool for biological study in these bacteria using gRNA that could be designed from a basic genome annotation.

## Conclusions

In this work, we revealed the general transferability but key strain-specific differences in the behavior of three CRISPRi systems in four diverse strains of *E. coli*. Correlations in fold repression of different gRNA designs between strains were high across all CRISPRi systems, although levels of repression and cellular growth toxicity could vary significantly between strains. Interestingly, these strain-specific differences did not appear to correlate with the evolutionary relationship between these strains. We demonstrated that CRISPRi using dual gRNA arrays and Lb-dCas12a typically increased repression of a target gene, markedly for the non-model UPEC strains CFT073 and UMN026, presenting an improved approach for regulating expression in these hosts.

We found that the dynamic range of repression often differed among the four *E. coli* strains, but similar patterns of gRNAs having significant repression for the CRISPRi systems were shared. Sp-dCas9 CRISPRi showed the greatest variation in repression for gRNA designs across the strains. However, gRNA targeting the non-template strand of the CDS or either strand of the –35 promoter region generally demonstrated significant repression in at least 3 strains, showing agreement with previous reports in MG1655 and other strains.^8,62^ Repression did not necessarily decrease when targeting sites further from the start of the CDS as has been observed in some studies^3,8,62^ but not others.^17,102^ Fn-dCas12a and Lb-dCas12a CRISPRi showed much better agreement in significant repression between strains. In both dCas12a systems, significant repression was observed for gRNA targeting the template strand of the CDS or the non-template strand of the promoter (*p* = 7.9 × 10^-6^ – 0.012), corroborating findings from the literature.^9,10^ The promoter we used here did not contain a PAM site to test targeting the template strand closely within the promoter, which is expected to effectively repression based on these studies. Correlations in repression between strains for the gRNA that were expected to show activity based on these literature reports remained moderate to strong (Pearson’s *r* = 0.665 – 0.990) compared to the correlations of all gRNA (**Figures S18-S20)**. Overall, our findings corroborate reported gRNA design rules for active repression from the literature for each CRISPRi system across each strain, with repression for these gRNA still correlating relatively well across the strains for each CRISPRi system.

However, our results revealed important strain-specific differences in both the magnitude of repression and cellular growth toxicity for each CRISPRi system across the *E. coli* strains. The variation in CRISPRi repression for the same gRNA designs between strains of the same bacterial species follow the few reports of such quantitative data in the literature.^51,59^ For each pair of strains, varying strengths of correlations in fold repression were found across the CRISPRi systems with a single gRNA (**Figures S6, S11, and S14**), even between the most phylogenetically closely-related EcN and CFT073 strains. Additionally, statistically significant differences in repression for single gRNA designs were found between all combinations of strains across the CRISPRi systems. CFT073 and EcN notably demonstrated the greatest number of differences of all strain pairs (12/35 across all systems), while MG1655 and UMN026 exhibited the fewest (3/35) (**Table S4**). This suggests that CRISPRi systems may not necessarily be more transferable between closer phylogenetic relatives of a species.

Due to the large set of CRISPRi designs and constructed strains (168+ strains) in this study and strong silencing we aimed to engineer, we chose to use a fluorescent protein reporter and high-throughput flow cytometry to quantify repression. As a single-cell based assay, flow cytometry allowed us to assess the cell-to-cell variability of expression within the population. This method also allowed us to detect subtle differences in strong silencing, as compared to other methods (e.g. RT-qPCR) for which the gene expression level can drop below the limit of detection for very strong silencing, preventing quantitative comparison. To provide sufficient PAM sites for testing, we switched from the eYFP output (of the RPU standard plasmid) to the sfGFP output for dCas12a. However, we tested Sp-dCas9 targeting *sfgfp* for a couple gRNA designs in MG1655, which showed statistically similar repression as gRNA targeting the same strand of *eyfp* in similar positions (one-way ANOVA with Tukey post-hoc analysis for repression at 0.5 ng/mL aTc, *p* > 0.05) (**Figure S21**).

Our study also revealed differing growth toxicity from expression of dCas proteins across the *E. coli* strains, which has not been previously studied or reported. CFT073 and MG1655 exhibited similar patterns in growth reduction from dCas expression across all CRISPRi systems yet with greater toxicity observed in MG1655 (**Figures 4, 6,** and **8**). EcN and UMN026 also exhibited similar growth toxicity for expression of Fn-dCas12a, but each demonstrated differing toxicity for the other dCas proteins. Overall, EcN showed the least growth reduction among all strains, and Lb-dCas12a showed the least toxicity among the dCas proteins. Specific growth was comparable between the different dCas proteins expressed at all inducer concentrations for all strains, except UMN026 that exhibited much slower growth for Sp-dCas9 expression than the dCas12a variants (**Figure S13**). For the UMN026 strain, our findings agree with reports of decreased toxicity for dCas12a systems relative to dCas9 systems,^70,71^ yet this relationship was not observed for the other *E. coli* strains.

Repression of a target gene using CRISPRi can be fine-tuned for different applications by varying the expression of the gRNA and to some extent dCas.^94,103^ We confirmed this for all gRNA designs in each CRISPRi system in the model strain MG1655 by varying the concentration of each inducer (**SI Note 1, Figures S22-S25**). A kinetics model for CRISPRi repression using our Sp-dCas9 system, which was based on a published model,^104^ generated predictions generally in agreement with observed repression in MG1655 at relatively low levels of expression of Sp-dCas9 (**SI Note 2, Figure S26**). This suggests that a relatively simple kinetics model may aid in the tuning of CRISPRi repression via inducible systems when dCas expression is not toxic. We investigated the interaction in the expression of CRISPRi components on measured repression using a design of experiments model for repression at varying expression levels of the dCas protein and gRNA for each CRISPRi system (**SI Note 3**). The reduced models from the results revealed that, at minimum, expression of both the dCas12a protein and gRNA were statistically significant for repression using both Fn-dCas12a and Lb-dCas12a CRISPRi, while only gRNA expression was a significant factor using Sp-dCas9 CRISPRi (**Figure S27, Tables S5** and **S6**). Altogether, the results presented here demonstrate various methods to fine-tune repression of a target gene.

Here, we developed the dual gRNA Lb-dCas12a CRISPRi system that achieved strong gene silencing in all four *E. coli* strains tested and provides a strategy to circumvent many of the observed strain-specific effects on repression. We demonstrated that repression could be enhanced using two gRNA targeting within the CDS as compared to one. When investigating the epistatic effects of the dual gRNA arrays, we assessed the agreement of observed repression to both additive and multiplicative models of the repression of each single gRNA, yet neither showed strong agreement for all strains (**Figures S28** and **S29**). Like the single gRNA designs, the repression using a dual gRNA design could be tuned by altering the level of expression of Lb-dCas12a and the gRNA array, as we observed in MG1655 (**SI Note 1**, **Figure S30**). The Lb-dCas12a dual gRNA approach provides a method to control gene silencing and often surpasses single gRNAs, yet even for this approach, not every dual gRNA design exhibited optimal repression in all strains.

This study provides evidence of the unexpected host-specific effects on repression efficiency and toxicity that can arise when using CRISPRi in different strains. Overall, our results emphasize the need to characterize CRISPRi systems in a strain before usage. The results demonstrate that CRISPRi repression and toxicity can be highly variable for different strains, and this work provides guidance to researchers on designing appropriate CRISPRi systems for various applications.

## Methods

### Bacterial strains, media, and growth conditions

The *Escherichia coli* strains MG1655, Nissle 1917 (EcN), CFT073, and UMN026 were utilized to characterize each CRISPRi system. Strains were assayed in M9 minimal media [6.78 g/L Na_2_HPO_4_ (Fisher Chemical), 3 g/L KH_2_PO_4_ (Sigma-Aldrich), 1 g/L NH_4_Cl (Sigma), 0.5 g/L NaCl (Sigma-Aldrich), 2 µM MgSO_4_ (Sigma-Aldrich), 0.1 mM CaCl_2_ (anhydrous; Sigma-Aldrich)], supplemented with 0.2% casamino acids (Acros Organics), 0.34 g/L thiamine hydrochloride (Fisher BioReagents), and 0.4% D-glucose (Fisher Bioreagents). Notably, plates for the non-model strains EcN, CFT073, and UMN026 could not be stored a 4 °C prior to starting the repression and growth toxicity assays. The strains NEB 5-alpha (New England Biolabs, NEB) or NEB 10-beta (NEB) were used for the construction and cloning of plasmids. Strains were grown in Luria-Bertani (LB; Fisher BioReagents) broth or on LB agar plates for cloning. Antibiotics were added to the media to maintain plasmids at the following concentrations: 50 µg/mL kanamycin (Goldbio) for all strains and spectinomycin (Goldbio) at 50 µg/mL (MG1655, NEB 5-alpha, and NEB 10-beta), 100 µg/mL (EcN and CFT073), or 300 µg/mL (UMN026). The gRNAs and dCas proteins were expressed using the chemical inducers isopropyl β-D-thiogalactopyranoside (IPTG; GoldBio) and anhydrotetracycline hydrochloride (aTc; Sigma-Aldrich), respectively.

### Design of gRNA

Python scripts were created to design gRNA for Sp-dCas9, Fn-dCas12a, and Lb-dCas12a based on a previous set of published scripts^3^. Briefly, the input parameters include the CRISPRi system, desired number of gRNA designs for the target sequence, strand of DNA to target, GC content range for the spacer sequence, and DNA sequences to avoid in the spacer sequence. For Sp-dCas9 gRNA design, the script can remove designs associated with the “bad seed” effect^17^ for *E. coli*. Off-target effects are predicted using an empirical penalty system based on the number and location of mismatches between the gRNA spacer sequence and potential off-targets in the host organism’s genome. Scripts were designed to incorporate the characteristics of each CRISPRi system (i.e., canonical and non-canonical PAM sequences, regions of gRNA spacer sequences, and penalty values for predicting off-target effects). A detailed description of the scripts is given in **SI Note 4** and **Figure S31**. The scripts and associated files are available at GitHub [https://github.com/AndrewsLabSynBio/CRISPRi_gRNA_library_design/tree/main].

The fluorescent output for Sp-dCas9 characterization used *eyfp* from a standard RPU plasmid (pAN1717^96^) with the promoter P1. However, this *eyfp* output gene contains few TTTV PAM sequences, so an *sfgfp* output gene with a sufficient number of PAM sequences was used to design gRNA for the dCas12a systems. gRNA designs targeting either strand of DNA and within the promoter region or along the length of the coding sequence (beginning, middle, and end) were chosen. Candidate designs were discarded if GC content was not between 30% – 80% (Sp-dCas9) or 25% – 80% (Fn-dCas12a and Lb-dCas12a). Off-target effects were predicted using Cas OFFinder^105^ with the following settings: SpCas9 from *Streptococcus pyogenes* (5′-NGG-3′) or LbCpf1 from *Lachnospiraceae* (5′-TTTV-3′), *Escherichia coli* host genome, up to 5 mismatches, and DNA and RNA bulge size of 0. Candidate designs were discarded if an off-target site contained only three mismatched base pairs with the target site or contained no mismatches in the seed region with only four mismatched base pairs. gRNA designs are named based on the target strand of the DNA (“N” for non-template, “T” for template) and targeting location from the first nucleotide of the target gene’s coding sequence, based on the first nucleotide of the seed region in the spacer sequence. Sequences and additional details on the design of each gRNA used in this work are provided in **Table S7**.

### DNA assembly

Plasmids used in this work are detailed in **Table S8**, with plasmid maps provided in **Figures S32** and **S33**. Destination vectors and the *sfgfp* output plasmid used in this work are available from Addgene. Genetic parts used in the construction of these plasmids are given in **Table S2** and provided in SBOL format in **SI File 2**. Plasmids were constructed using Type IIS DNA assembly (**SI Extended Methods**). CRISPRi destination vectors were designed for efficient insertion of arbitrary gRNA sequences using BsaI (Sp-dCas9) or BbsI (Fn-dCas12a and Lb-dCas12a) to replace a LacZα fragment, allowing for blue-white screening of colonies. gRNA sequences were inserted into destination vectors using either DNA fragments flanked with Type IIS recognition sequences or with appropriate 5′ overhangs.

### Electrocompetent cell preparation for *E. coli* strains

Electroporation was used to introduce constructed plasmids into MG1655, EcN, CFT073, and UMN026. To prepare electrocompetent cells for MG1655 and EcN, a single colony from a freshly streaked agar plate was incubated in 2 mL of LB and grown overnight at 37 °C and 250 rpm. The culture was then diluted down to an OD_600_ of 0.01 A in 50 or 100 mL of fresh LB media and grown under the same conditions to an OD_600_ of 0.55 – 0.65 A (mid-exponential phase). The culture was then chilled in an ice-water bath for 10 minutes and transferred to pre-chilled sterile 50 mL conical centrifuge tubes. All subsequent steps were performed on ice and used sterilized wash solutions. The tubes were centrifuged at 8,000 g at 4 °C for 10 minutes (Eppendorf 5810 R) to pellet the cells, and the supernatant was carefully decanted. The cell pellet was gently resuspended by pipetting and swirling in 50 mL of ice-cold Milli-Q water, starting with resuspension in 1 mL. This washing step was repeated twice more, first resuspending the cells in 50 mL ice-cold Milli-Q water and then 50 mL ice-cold 10% v/v glycerol. After the final centrifugation, the cells were gently resuspended in 700 µL of ice-cold 10% glycerol and portioned into 50 µL aliquots. Aliquots were either used immediately for electroporation or stored at –70 °C for up to 6 months.

For CFT073 and UMN026, modifications to this protocol were made as follows. Strains were grown in no-salt LB media (10 g/L tryptone, 5 g/L yeast extract) overnight, then cultures were diluted by adding 1 mL of the overnight into 50 mL of fresh no-salt LB media and grown to an OD_600_ between only 0.3 and 0.4 A. Volumes of wash solutions used to resuspend the cells decreased every step, starting with 10 mL Milli-Q water, then 7.5 mL Milli-Q water, and finally 5 mL 10% glycerol. Cells were resuspended in a final volume of 500 µL of 10% glycerol and divided into 100 µL aliquots for electroporation. Notably, this protocol could also be used to make EcN electrocompetent.

### Transformation of *E. coli* strains

Electroporations for the non-model strains were slightly modified from that for MG1655. For MG1655, 100 ng of purified output plasmid and 200 ng of CRISPRi plasmid were added to a 50 µL aliquot of electrocompetent cells (DNA:cell ratio ≤ 1:5) and incubated on ice for 30 minutes before transfer to a pre-chilled 1 mm electroporation cuvette. Cells were electroporated (Bio-Rad MicroPulser) using the Ec1 settings (1.8 kV, 1 pulse, no time constant). Immediately after electroporation, 500 µL of pre-warmed SOC recovery media was pipetted into the sample and transferred to a 1.5 mL microfuge tube with an additional 500 µL of SOC. Cells were recovered for an hour at 37 °C and 250 rpm and then plated on LB agar plates with the appropriate antibiotics and grown overnight at 37 °C.

For CFT073 and UMN026, the same protocol for electroporation was performed as in MG1655, except 500 ng of each plasmid was added to 100 µL aliquots of cells due to lower observed transformation efficiencies compared to MG1655. For EcN, however, co-transformation of plasmids was not successful. Therefore, each plasmid was transformed into EcN via electroporation separately, starting with the common reporter plasmid. The resulting strains were then made electrocompetent again prior to electroporating the appropriate CRISPRi plasmid using the same procedure as previously described, except with spectinomycin added to the media to maintain the plasmid. Electroporation in EcN was performed otherwise as described for the CFT073 and UMN026 strains.

### Flow cytometry and data analysis

All experiments to characterize CRISPRi repression were conducted by measuring single-cell fluorescence using flow cytometry with the BD Accuri C6 or DUAL LSRFortessa. Each strain of *E. coli* was transformed with a CRISPRi plasmid with a targeting gRNA and the appropriate output plasmid containing the reporter gene *eyfp* (Sp-dCas9) or *sfgfp* (Fn-dCas12a and Lb-dCas12a). Strains were streaked on LB agar plates with appropriate antibiotics and incubated overnight at 37 °C. Additionally, both the negative control (wildtype) and positive control (containing the pAN1717^7,96^ RPU standard plasmid) strains were streaked for the conversion of cell fluorescence measured in arbitrary units to relative promoter units (RPU) or relative expression units (REU). An individual colony was inoculated into 200 µL of M9 media with appropriate antibiotics in a sterile U-bottom 96-well plate and sealed with a breathable Aeraseal (Excel Scientific). Then, cells were incubated at 37 °C and 1000 rpm in a plate shaking incubator (Elmi DTS-4) for 16 hours.

After incubation, cells were serially diluted via two sequential 15 µL dilutions of the culture into 185 µL of M9 media containing antibiotics as needed (total 177.8-fold dilution), followed by a 3-hour incubation under the same conditions. Next, 15 µL of the incubated cell culture was diluted into 185 µL of fresh M9 media with appropriate antibiotics, then 3 µL of this diluted cell suspension was added to 145 µL of M9 media with antibiotics and inducers as needed. After dilution into this fresh media, cultures were incubated for an additional 5 hours under the same growth conditions. For measuring fluorescence, cells were diluted into PBS containing 2 mg/mL kanamycin and incubated at room temperature for 30 minutes prior to flow cytometry. Fluorescence was measured on the FL1-A and FITC channels on the C6 and LSRFortessa cytometers, respectively. To decrease noise, fluorescence data were collected using thresholding values of 25,000-27,000 (FSC-H) and 1,250 (SSC-H) for the C6 cytometer and default thresholds for the LSRFortessa cytometer (voltages set to FSC = 700 V; SSC = 340 V; and FITC = 399 V). For each sample, 4,500 – 10,000 cells were analyzed (except gRNA-175T for Fn-dCas12a in UMN026 with 2,200 cells analyzed for one sample).

To induce dCas protein expression, aTc was added at 0.25 ng/mL for MG1655 (all systems) and 0.125 ng/mL for EcN (all systems). For CFT073 and UMN026, aTc was added at 0.25 ng/mL for Sp-dCas9, 0.5 ng/mL for Fn-dCas12a, and 2 ng/mL for Lb-dCas12a. These aTc concentrations were chosen as they demonstrated repression for all CRISPRi systems from preliminary titration assays (**Figure S5**) without inhibiting growth such that 5,000 cell events could typically be collected. The corresponding P_Tet_ promoter activity has significant differences across the strains,, except for Sp-dCas9 (**Figure S34A**). gRNA expression was induced with 1 mM IPTG for all strains.

Analysis of all flow cytometry data for the four *E. coli* strains was performed using FlowJo (v10.8.0), incorporating a custom FSC-A vs SSC-A gate for each strain and calculating the median of cell fluorescence in arbitrary units as the fluorescence for the sample (AU_sample_). Arbitrary units of fluorescence were then transformed into relative expression units (REU) using the following equation:

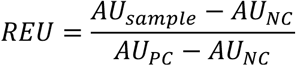

where AU_sample,_, AU_NC_, and AU_PC_ represent the median fluorescence of the sample, the wildtype cells for the strain (AU_NC_), and the cells containing the pAN1717^7,96^ RPU reference plasmid for the strain (AU_PC_), respectively. The REU for the standard reference plasmid is set to 1. Samples with negative REU values were set to a limit of detection of 0.001 REU. Representative histograms of the positive and negative controls for each strain are given in **Figure S4** and of an induced and uninduced sample of each combination of gRNA design, CRISPRi system, and strain in **Figures S35-S50**.

### Toxicity assays for cell growth inhibition in E. coli

To quantify cellular growth toxicity due to dCas expression, cell growth kinetics were measured using a plate reader with different induction levels of the dCas protein. For each strain of *E. coli*, a single colony of the wildtype and strains containing the CRISPRi plasmid were picked from an LB agar plate and inoculated in 5 mL M9 media with antibiotics as needed. The CRISPRi plasmids were those containing the gRNA that demonstrated the greatest repression among the strains (pSR2017 with –144T for Sp-dCas9; pSR2022 with –119N for Fn-dCas12a; pSR3032 with – 119N for Lb-dCas12a). Cultures were incubated overnight at 37 °C in a shaking incubator. The overnight cultures were diluted to an OD_600_ of 0.05 A in 2 mL of M9 media with antibiotics and the specified aTc concentration (0, 0.25, 2.0, or 4.0 ng/mL aTc for strains containing a CRISPRi plasmid). IPTG was not added to the media. 100 µL of each culture was transferred to a well in the middle of a flat-bottom 96-well plate. To reduce evaporation, the outer two rows were kept blank and filled with 200 µL of water, along with any empty wells. The plate was closed with a fitted lid and sealed with parafilm. Cell growth was measured using a microplate reader (Agilent Biotek Synergy HTX), recording the absorbance at 600, 900, and 977 nm every 15 minutes for 8 hours using the BioTek Gen5 software. The plate was incubated at 37 °C, with continuous shaking (810 rpm, orbital path) when absorbance was not being measured. Reported OD measurements are the pathlength corrected values (0.1 cm) for OD_600_.

The pathlength corrected OD_600_ measurements in the exponential growth phase were fit to the following exponential growth equation:

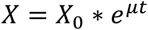

where *X* and *X_0_* are the OD_600_ at times *t* and 0 hours, respectively, and *µ* is the exponential specific growth rate. The data for each replicate were fit to at least 12 points of the log-linear portion of the growth data using the growthrates package^106,107^ in R and visually inspected to ensure appropriate fit. Specifically, the fits were performed using the all_easylinear() function in this package to fit each replicate of all samples at once, with h = 12 to fit at least 12 data points. The resulting fit parameters for each replicate of each combination of strain, CRISPRi system, and aTc concentration are provided in **SI File 1**.

### Statistical analysis

Statistical analyses were performed using custom scripts written in R 4.3.0.^108^ Throughout analysis, results were determined to be significant if *p* < 0.05, with greater significance given for *p* < 0.01 and *p* < 0.001. The results for these analyses are available in **SI File 1**.

All *t*-tests were performed using the t.test() function with options adjusted depending on the type of *t*-test and dataset. Welch’s one-sided, one-sample *t*-tests were performed on the log_10_-transformed fold repression data collected for each combination of gRNA, CRISPRi system, and strain to determine statistically significant repression greater than 1 (µ = 0 on a log scale). Welch’s one-sided, two-sample, unpaired *t*-tests were performed on the specific growth rates and OD_600_ values at 4 hours to determine if these variables measured for the assayed strain were lower than those in the corresponding wildtype strain. Similarly, this type of *t*-test was also performed between the log_10_-transformed measured fold repression data for the dual gRNA designs and each corresponding single gRNA, testing if the repression using the dual gRNA array was greater than that of each single design. Welch’s two-sided, two-sample, unpaired *t*-tests were performed between the log_10_-transformed fold repression data of the Lb-dCas12a dual gRNA designs comprised of the same single gRNAs in reverse order to determine if order of the gRNA affected measured repression.

One-way ANOVA was performed using the aov() function, and Tukey post-hoc analysis was performed using the TukeyHSD() function if the *p*-value of the *F*-statistic was less than 0.05. Default options were used for both functions. These analyses were executed on the following sets of data to determine if significant differences existed between samples in each set. Datasets include: (i) output from P_Tet_ and P_Tac_ (in log_10_-transformed RPU) across the strains at the inducer concentrations used; (ii) output values (in log_10_-transformed REU) across gRNA designs for each combination of strain, CRISPRi system, and CRISPRi induction level (induced and uninduced); (iii) log_10_-transformed fold repression between strains for each combination of gRNA and CRISPRi system, (iv) log_10_-transformed fold repression across CRISPRi systems for gRNA showing the greatest average repression and for gRNA targeting the preferred strand of DNA; (v) specific growth rates between dCas proteins for each combination of aTc concentration and strain; and (vi) log_10_-transformed fold repression between Sp-dCas9 CRISPRi gRNA targeting *eyfp* or *sfgfp* at each aTc concentration in MG1655. Data with nonsignificant results from the ANOVA analysis (*F* > 0.05) were considered to have nonsignificant *p*-values (≥ 0.05).

For each CRISPRi system, two-sided Spearman and Pearson correlations were calculated using the cor.test() function for the log_10_-transformed data for average fold repression between all combinations of strains. These correlations were also calculated for the untransformed fold repression data for all gRNA that were expected to be active for each CRISPRi system (–123N, –89N, +34N, +334N, +664N, –144T, and –128T for Sp-dCas9; –119N, –175T, +20T, +139T, +339T, and +666T for both Fn-dCas12a and Lb-dCas12a). Correlations were classified as strong if *r* (or *ρ*) ≥ 0.80, moderate if 0.80 > r ≥ 0.50, and weak if r < 0.50.

## Author Information

### Corresponding Author

Lauren B. Andrews – Department of Chemical and Biomolecular Engineering, University of Massachusetts Amherst, Amherst, MA 01003, United States; Molecular and Cellular Biology Graduate Program, University of Massachusetts Amherst, Amherst, MA 01003, United States; Biotechnology Training Program, University of Massachusetts Amherst, Amherst, MA 01003, United States

### Authors

Hyerim Ban – Molecular and Cellular Biology Graduate Program, University of Massachusetts Amherst, Amherst, MA 01003, United States; Biotechnology Training Program, University of Massachusetts Amherst, Amherst, MA 01003, United States

Stephen N. Rondthaler – Department of Chemical and Biomolecular Engineering, University of Massachusetts Amherst, Amherst, MA 01003, United States

Matthew Lebovich – Department of Chemical and Biomolecular Engineering, University of Massachusetts Amherst, Amherst, MA 01003, United States; Biotechnology Training Program, University of Massachusetts Amherst, Amherst, MA 01003, United States

Marcos Lora – Department of Chemical and Biomolecular Engineering, University of Massachusetts Amherst, Amherst, MA 01003, United States

Brandon Ugbesia – Department of Chemical and Biomolecular Engineering, University of Massachusetts Amherst, Amherst, MA 01003, United States

## Author Contributions

H.B., S.N.R., and L.B.A. conceived the project and designed experiments. H.B., S.N.R., M.L., M.A.L., and B.U. performed experiments. H.B., S.N.R., and L.B.A. analyzed data. S.N.R. created software and scripts. H.B., S.N.R., and L.B.A. wrote the original draft, and all authors reviewed and edited the manuscript.

## Notes

The authors declare no competing financial interest.

## Acknowledgements

Funding that supported this work include the National Science Foundation (NSF) Award No. DMR-1904901, the Marvin and Eva Schlanger faculty fellowship, a seed grant from the UMass ADVANCE program (funded by NSF Awards No. EES-1824090), NSF Award No. CBET-1943695, and startup funds from the University of Massachusetts Amherst to LBA. Additionally, we thank the NSF Graduate Research Fellowship (Award No. DGE-1451512) for supporting SNR and the UMass Biotechnology Training Program (NIH Award T32 GM135096) for fellowship support to ML and HB. We thank Christopher Voigt for the pAN-PTet-dCas9 plasmid (Addgene #62244) and the Flow Cytometry Core Facility at UMass.

