## Supporting Information for "Cross-strain transferability of CRISPRi systems and design rules from laboratory to clinical *Escherichia coli* strains"

### Supplementary Figures

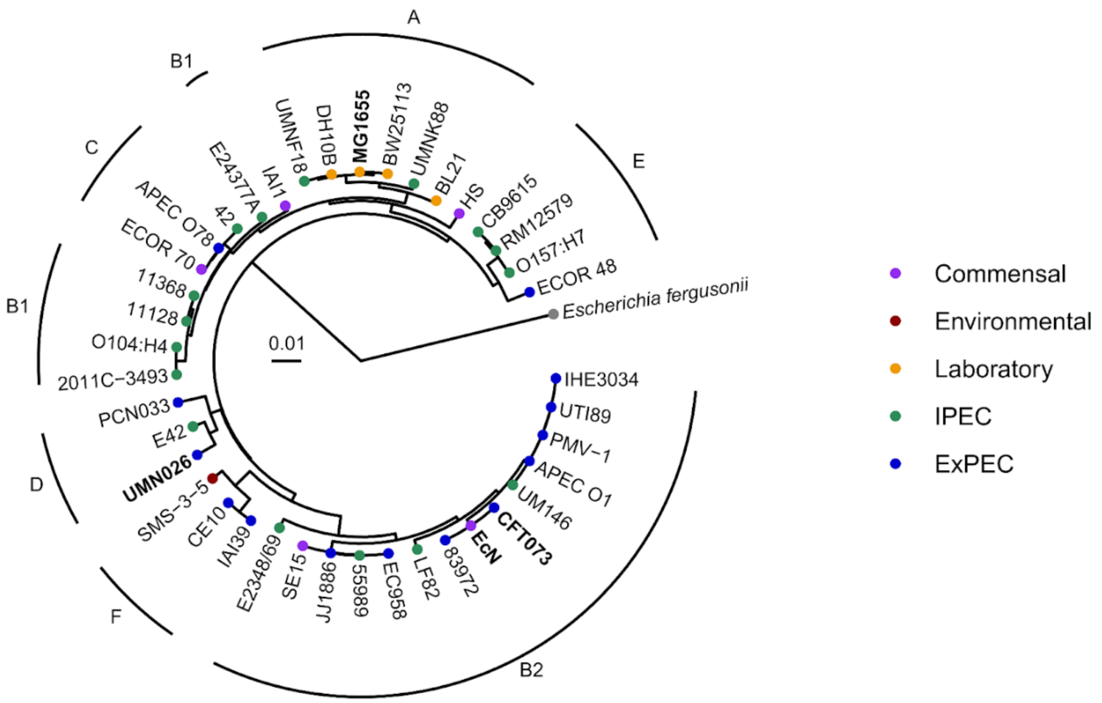

**Figure S1. Phylogenetic tree of diverse strains of *Escherichia coli***

A phylogenetic tree of 40 diverse strains of *E. coli* was generated using multi-locus sequence analysis of seven housekeeping genes and the maximum likelihood method, with *Escherichia fergusonii* included to root the tree (**Extended Methods**). The typical ecological niche of each strain is denoted by the color of the point for each strain with pathogenic strains characterized as intestinal pathogenic *E. coli* (IPEC) or extraintestinal pathogenic *E. coli* (ExPEC). Phylogenetic analysis The phylogroup for each strain, as determined using ClermonTyping,[1] is denoted in the outer chords. Strains in bold are those chosen to study in this work. The scale bar of 0.01 signifies a genetic distance with one nucleotide (nt) change per 100 nt.

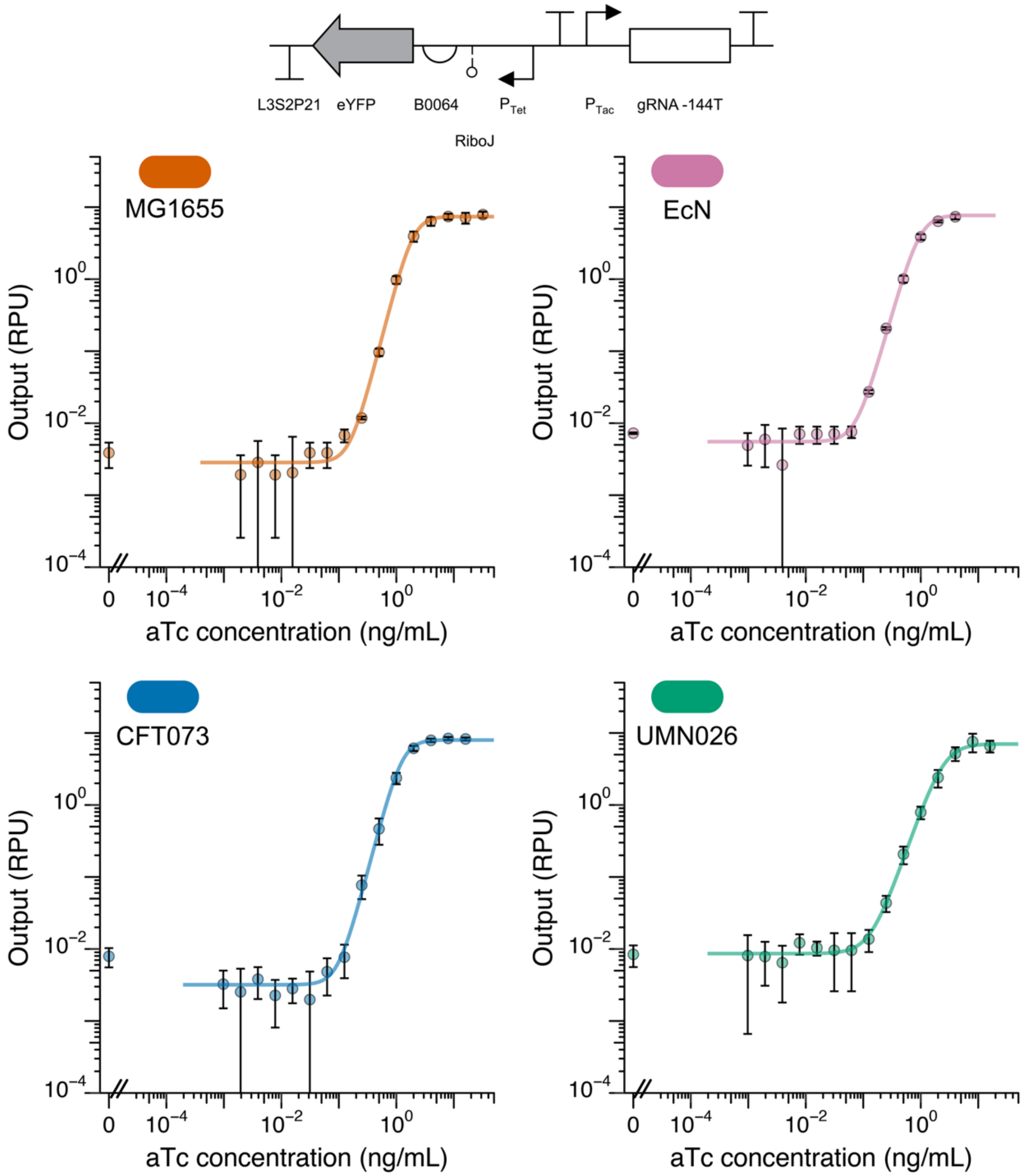

**Figure S2. Response curves for $P_{Tet}$ characterization in four *E. coli* strains**

To determine the sensor response function, the inducible promoter $P_{Tet}$ was placed upstream of a standard *eyfp* cassette (pSR2010) and in the same location as the dCas protein in the CRISPRi plasmids. Different concentrations of the inducer anhydrotetracycline (aTc) were added to the media in separate samples, and the resulting cell fluorescence was measured using single-cell flow cytometry after 5 hours of induction. Fluorescence was converted to relative promoter units (RPU), and the response curve (line) was determined by fitting the data to the Hill equation (**Extended Methods**). The response function parameters for each strain are given in **Table S3**. Markers are the mean of three identical experiments performed on three different days with error bars depicting the standard deviation.

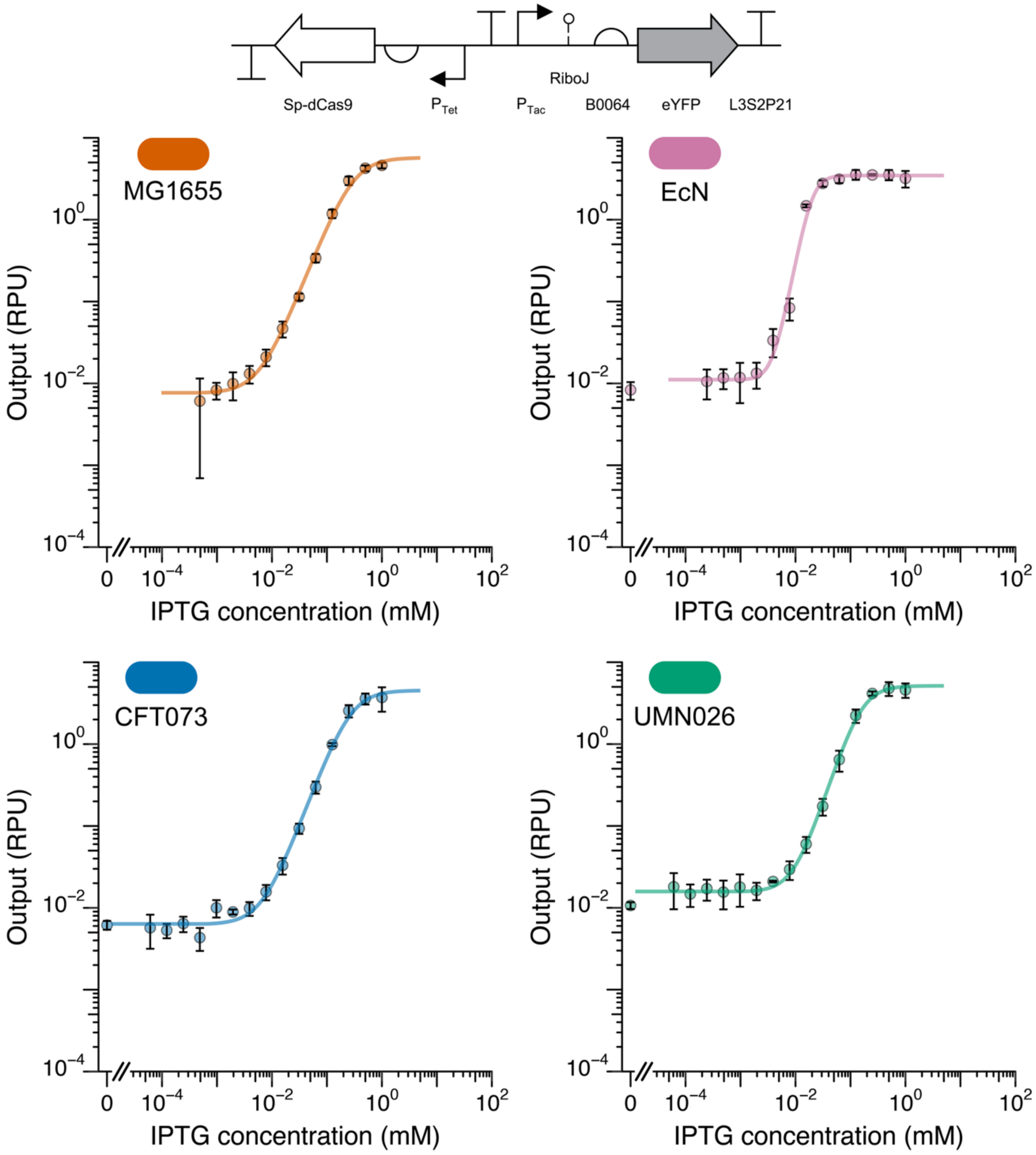

**Figure S3. Response curves for $P_{Tac}$ characterization in four strains of *E. coli***

To determine the sensor response function, he inducible promoter $P_{Tac}$ was placed upstream of a standard *eyfp* cassette (pSR2011) and in the same location as the dCas protein in the CRISPRi plasmids. Different concentrations of the inducer isopropyl β-D-thiogalactopyranoside (IPTG) were added to the media in separate samples, and the resulting cell fluorescence was measured using single-cell flow cytometry after 5 hours of induction. Fluorescence was converted to relative promoter units (RPU), and the response curve (line) was determined by fitting the data to the Hill equation (**Extended Methods**). The response function parameters for each strain are given in **Table S3**. Markers are the mean of three identical experiments performed on three different days with error bars depicting the standard deviation.

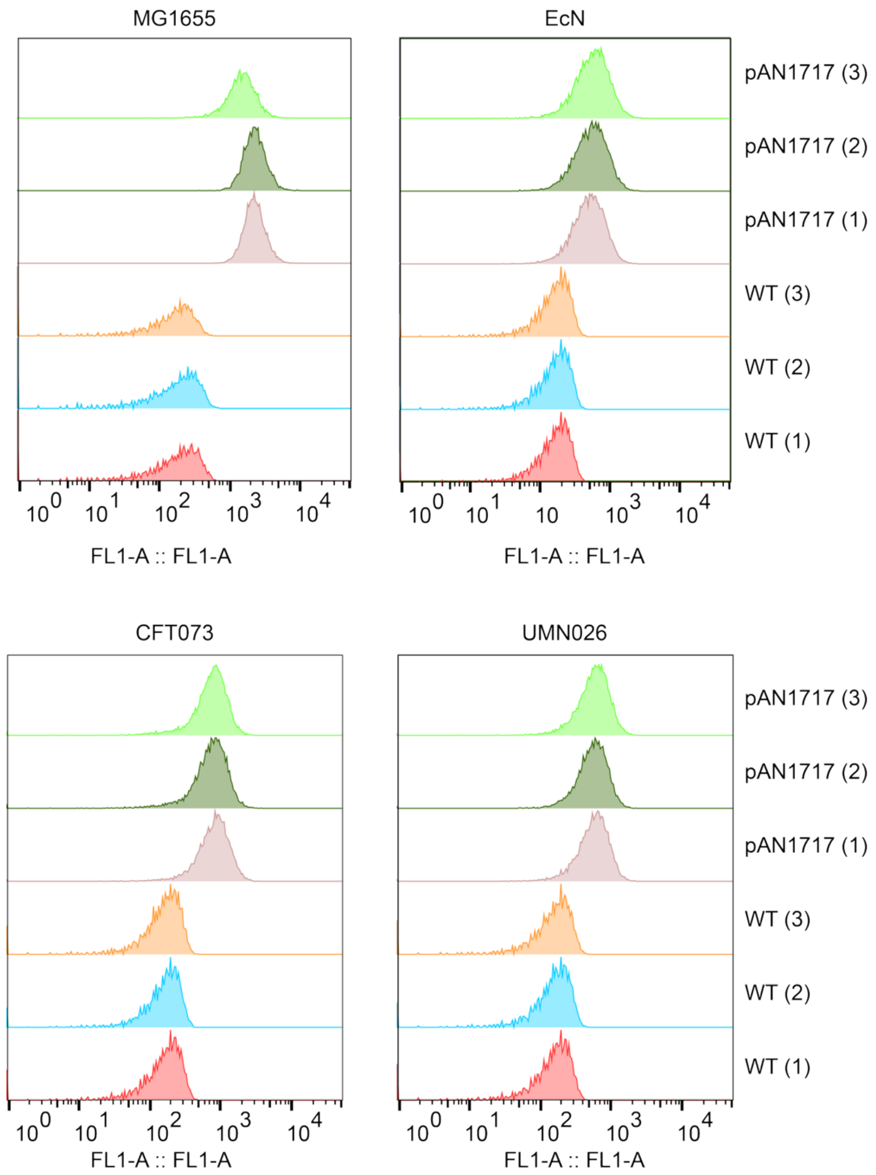

**Figure S4. Representative histograms of fluorescence for wildtype and RPU standard**

Three representative histograms of the normalized cell counts for fluorescence of the wildtype cells (WT) and positive control (containing the pAN1717[2,3] RPU standard reference plasmid) as measured using flow cytometry in MG1655, EcN, CFT073, and UMN026. Histograms are provided for each control in each strain measured on three different days. Each histogram includes > 4,500 cells.

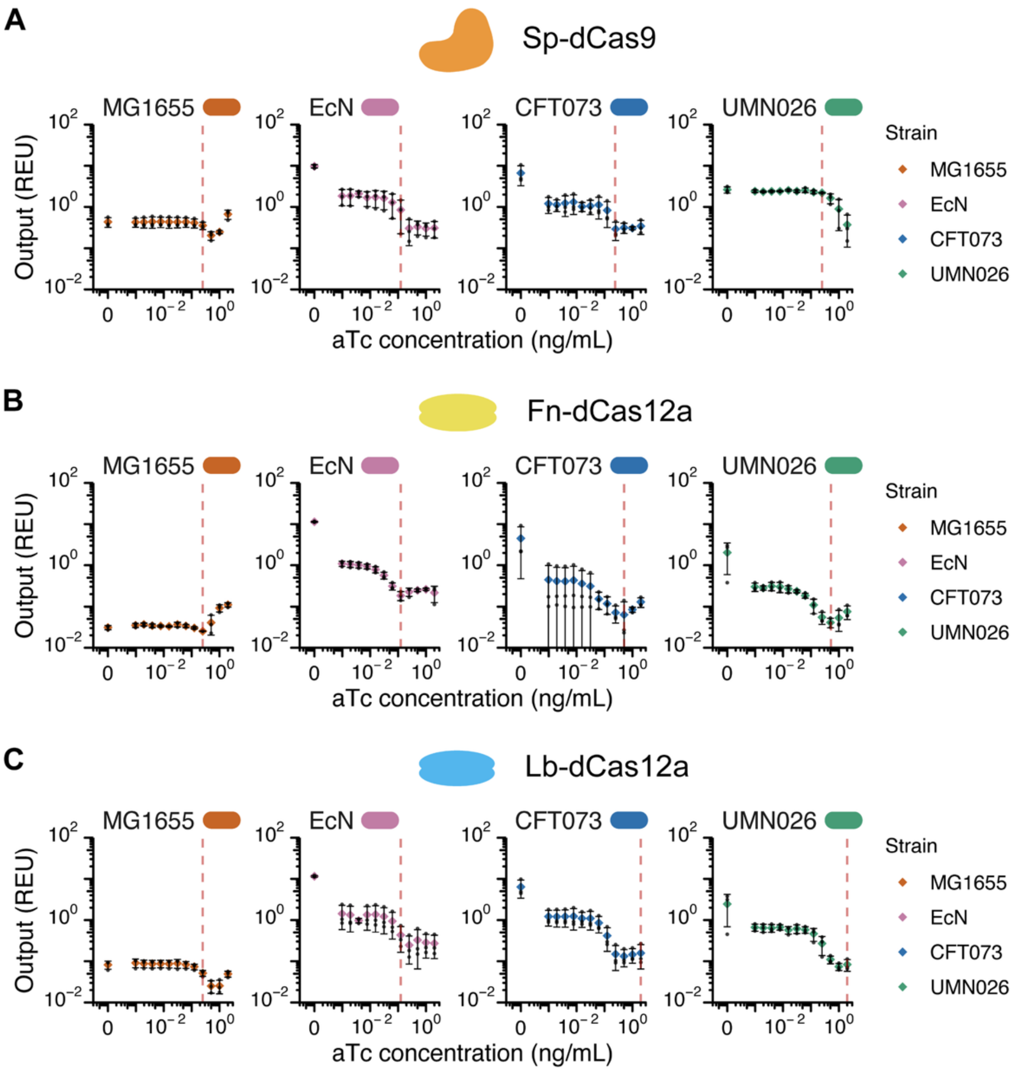

**Figure S5. Preliminary titration experiments for repression assays**

The expression of **(A)** Sp-dCas9, **(B)** Fn-dCas12a, and **(C)** Lb-dCas12a was titrated in MG1655 EcN, CFT073, and UMN026 by varying the amount of aTc in the media. The expression of the gRNA was fully induced with the addition of 1 mM IPTG to the media. The gRNA was selected as those expected to be active and targeting the promoter region or near the start of the coding sequence for CRISPRi using Sp-dCas9 (–144T for all strains), Fn-dCas12a (–119N for MG1655 and +20T for EcN, CFT073, and UMN026), and Lb-dCas12a (+20T for UMN026 and –119N for MG1655, EcN, and CFT073). Fluorescence was assessed using flow cytometry and converted to relative expression units (REU) (**Methods**). The red dashed line is the aTc concentration chosen for the repression assays in each strain for each CRISPRi system. Mean (diamond markers) of three identical experiments (black circles) performed on different days (*n* > 4500 cells per sample) with error bars depicting standard deviation.

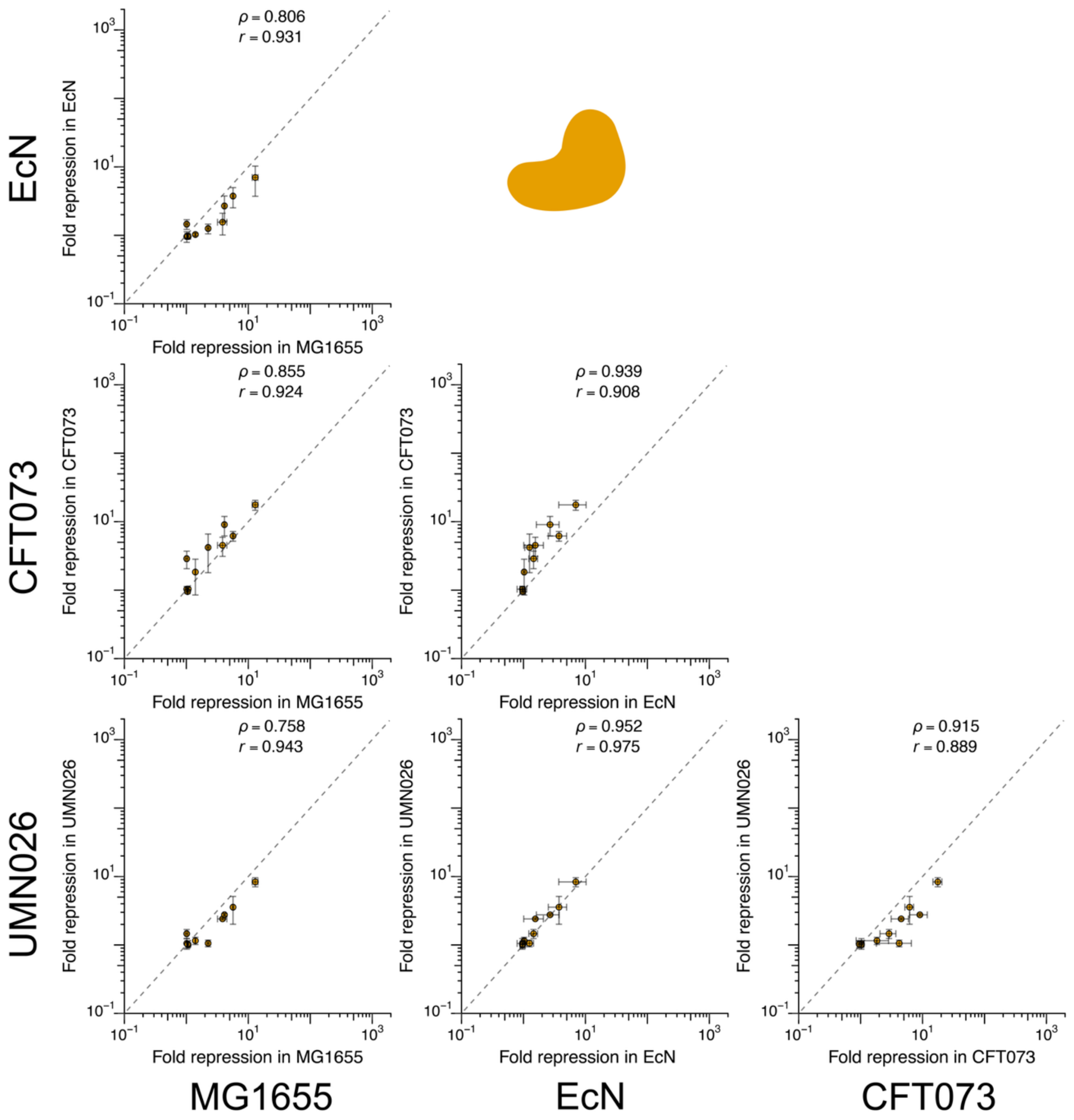

**Figure S6. Correlations of repression for Sp-dCas9 CRISPRi between strains**

The average measured fold repression for all gRNA designs of the Sp-dCas9 system for all pairwise combinations of MG1655, EcN, CFT073, and UMN026 are shown. The diagonal y = x (dashed line) is included for comparison. Pearson (*r*) and Spearman (*ρ*) correlations were performed on the $\log_{10}$-transformed fold repression values of each dataset and are indicated. The *p*-value for each correlation is provided in **SI File 1**.

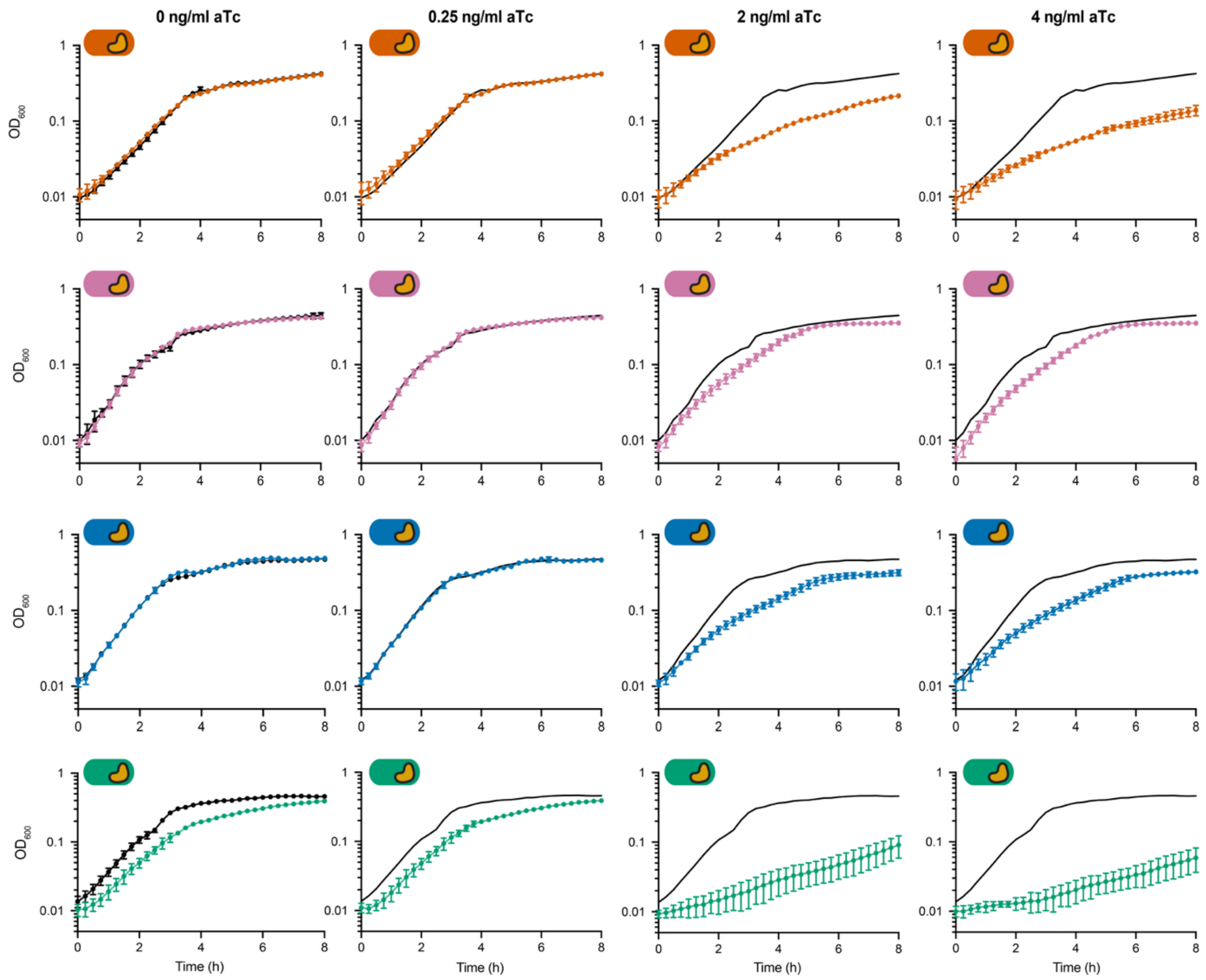

**Figure S7. Growth curves for Sp-dCas9 expression in *E. coli* strains**

Growth curves were performed for MG1655 (orange), EcN (pink), CFT073 (blue), and UMN026 (green) at different induction levels for expression of Sp-dCas9 in plasmid pSR2017 via the indicated concentration of aTc in the media (0, 0.25, 2, and 4 ng/mL aTc shown). Absorbance at 600 nm ($OD_{600}$) was measured at 15-minute intervals during an 8-hour incubation in a microplate reader (**Methods**). Points indicate the mean of three identical experiments performed on three different days, and error bars are the standard deviation. The growth curves for the wildtype cells for each strain (WT, black) without any inducer are shown for comparison in each plot with the error bars for the wildtype only drawn in the first column (0 ng/mL aTc).

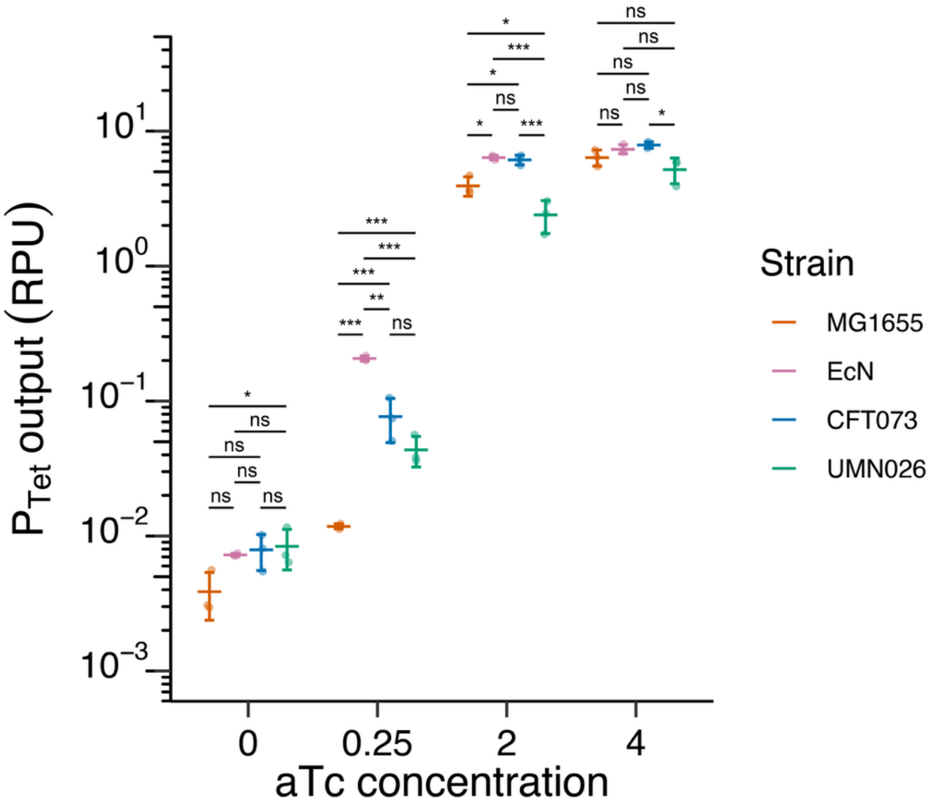

**Figure S8. Comparison of $P_{Tet}$ output used for growth toxicity assays**

The promoter output from $P_{Tet}$ with 0, 0.25, 2, and 4 ng/mL aTc is given in relative promoter units (RPU) for MG1655 (orange), EcN (pink), CFT073 (blue), and UMN026 (green), as determined from sensor response characterization in each strain (**Figure S2**). The results of a one-way ANOVA with Tukey post-hoc analysis are provided for the $\log_{10}$-transformed output values at each aTc concentration with results indicated as not significant (ns) or significant (*$p < 0.05$, **$p < 0.01$, and ***$p < 0.001$). Values are reported in **SI File 1**. The arithmetic mean (dash) of three identical experiments (points) performed on three separate days are plotted with error bars showing SD.

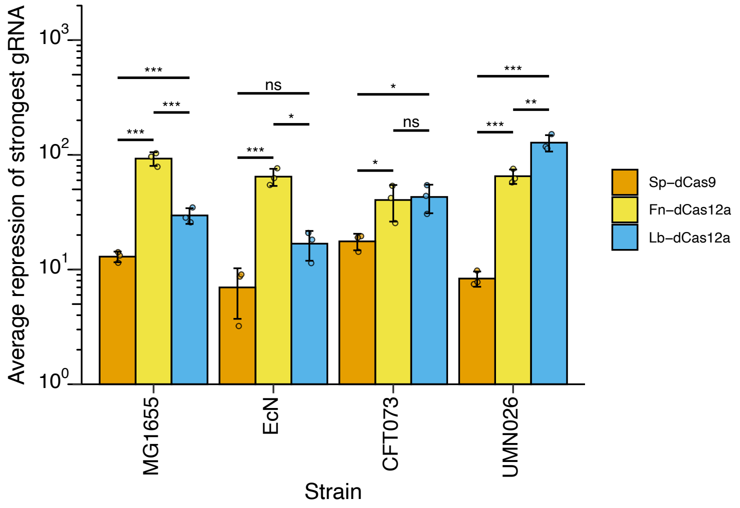

**Figure S9. Maximal observed repression for the CRISPRi systems**

Fold repression of the gRNA showing the maximum average repression in Sp-dCas9, Fn-dCas12a, and Lb-dCas12a in MG1655, EcN, CFT073, and UMN026 is plotted. The gRNA displayed are as follows: –144T for Sp-dCas9 in all strains, –119N (MG1655, UMN026) or +20T (EcN, CFT073) for Fn-dCas12a, and –119N (MG1655, EcN, UMN026) or +20T (CFT073) for Lb-dCas12a. One-way ANOVA with Tukey post-hoc analysis was performed between the $\log_{10}$-transformed repression values for each strain and results are indicated as not significant (ns) or significant (*$p < 0.05$, **$p < 0.01$, and ***$p < 0.001$). Results are provided in **SI File 1**. Bars are the arithmetic mean of three identical experiments (points) performed on three separate days with error bars showing the standard deviation.

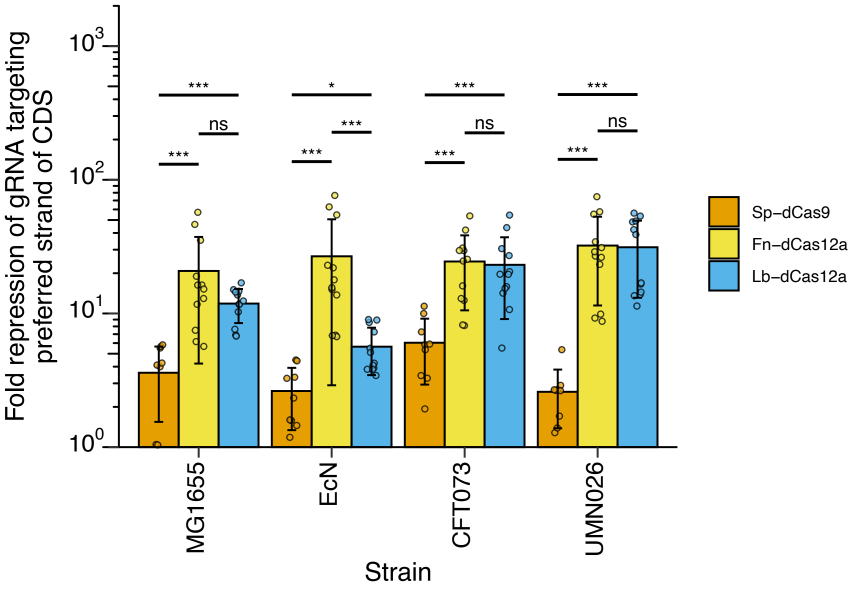

**Figure S10. Observed CRISPRi repression for gRNA targeting preferred DNA strand**

The fold repression of the gRNA targeting the non-template strand (Sp-dCas9) or template strand (Fn-dCas12a and Lb-dCas12a) in MG1655, EcN, CFT073, and UMN026 is provided. The gRNA included in this analysis are as follows: +34N, +334N, and +664N for Sp-dCas9 and +20T, +139T, +339T, and +666T for both dCas12a systems. One-way ANOVA with Tukey post-hoc analysis was performed between the $log_{10}$-transformed repression values for each strain, and results are indicated as not significant (ns) or significant (*$p < 0.05$, **$p < 0.01$, and ***$p < 0.001$). Values are given in **SI File 1**. Bars are the arithmetic mean of all replicates (points) for each gRNA of the CRISPRi system with error bars showing the standard deviation. The data for each gRNA were collected by performing three identical experiments on three separate days.

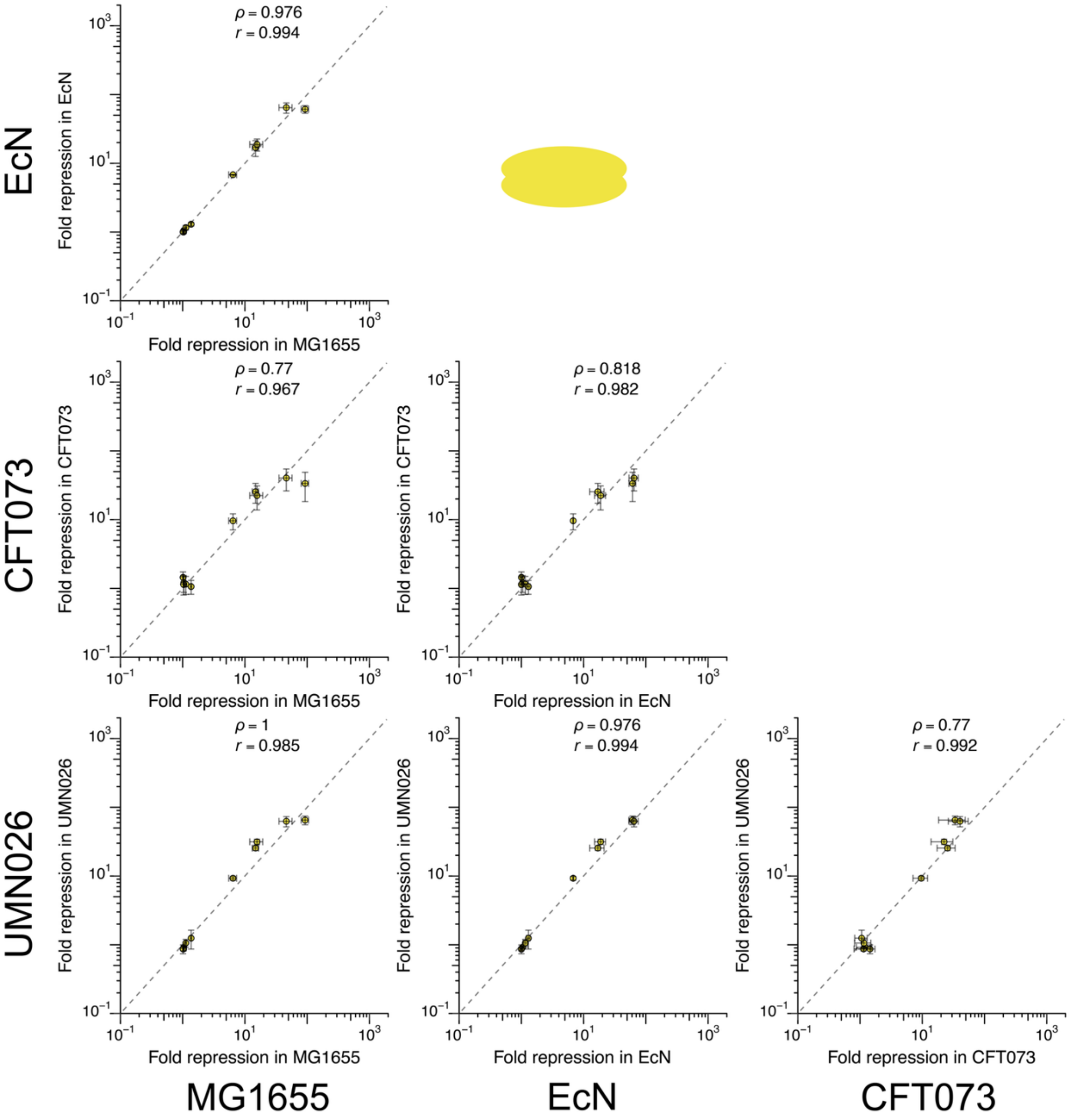

**Figure S11. Correlations of repression for Fn-dCas12a CRISPRi between strains**

The average fold repression values for all gRNA designs of the Fn-dCas12a system for all pairwise combinations of MG1655, EcN, CFT073, and UMN026 strains are shown. The diagonal y = x (dashed line) is included for comparison. Pearson ($r$) and Spearman ($\rho$) correlations were calculated on the $\log_{10}$-transformed fold repression values of each dataset, and values are reported. The $p$-value for each correlation is provided in **SI File 1**.

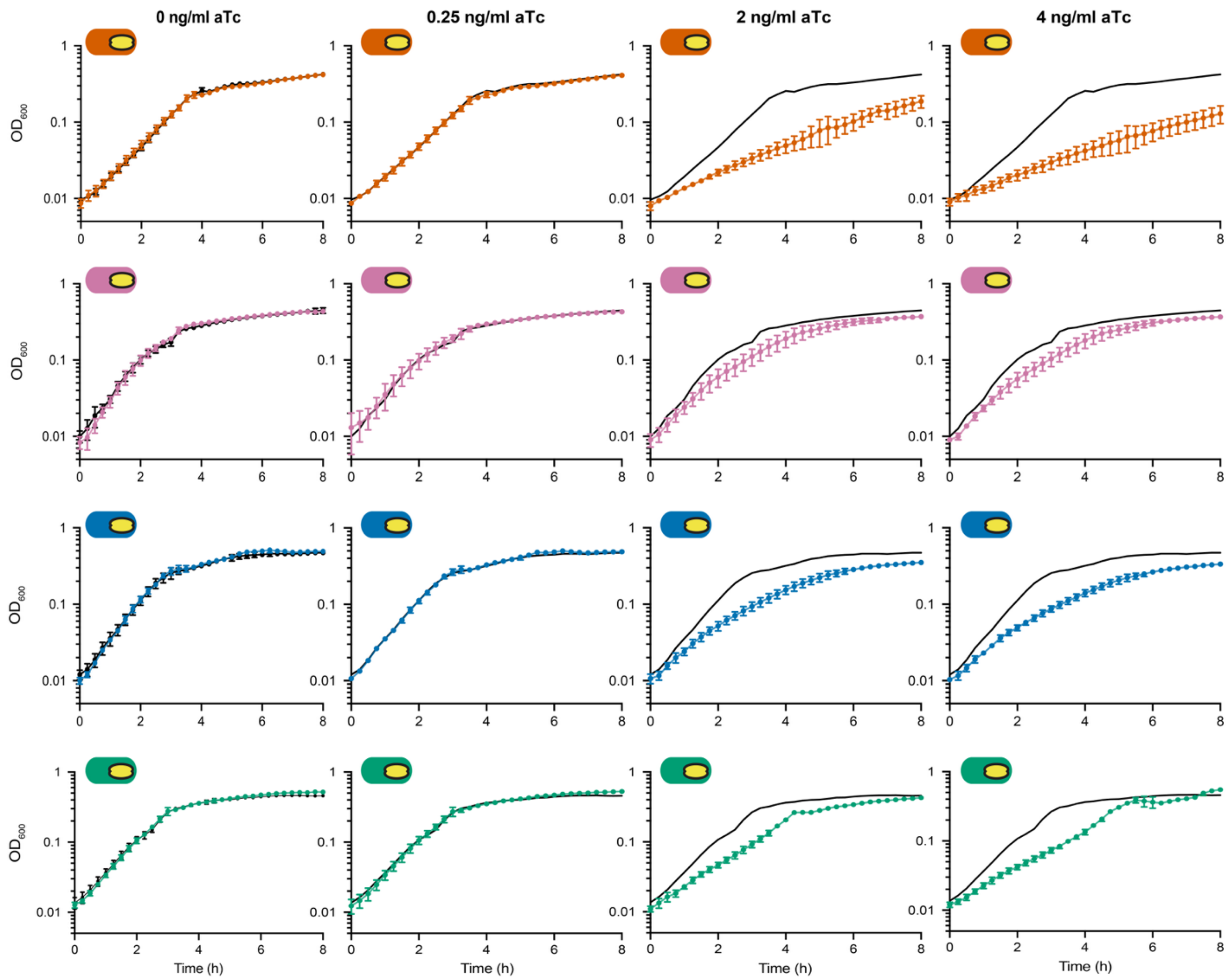

**Figure S12. Growth curves for Fn-dCas12a expression in *E. coli* strains**

Growth curves were performed for MG1655 (orange), EcN (pink), CFT073 (blue), and UMN026 (green) at different induction levels for expression of Fn-dCas12a in plasmid pSR2022 via the indicated concentration of aTc in the media (0, 0.25, 2, and 4 ng/mL aTc shown). Absorbance at 600 nm ($OD_{600}$) was measured at 15-minute intervals during an 8-hour incubation in a microplate reader (**Methods**). Points indicate the mean of three identical experiments performed on three different days, and error bars are the standard deviation. The growth curves for the wildtype cells for each strain (WT, black) without any inducer are shown for comparison in each plot with the error bars for the wildtype only drawn in the first column (0 ng/mL aTc).

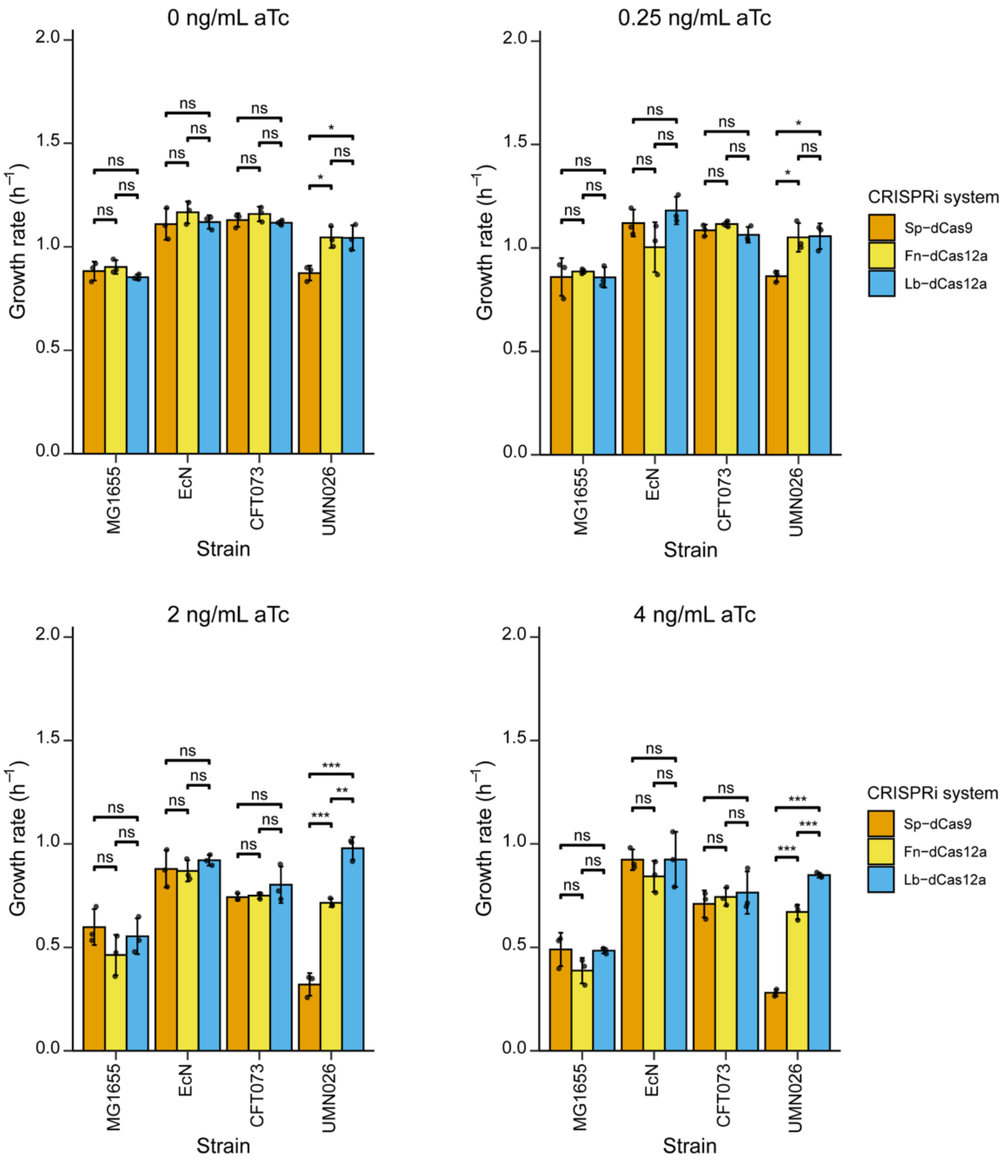

**Figure S13. Comparison of specific growth rates with dCas expression across strains**

Data from the growth curve experiments (**Figures S7, S12 and S15**) were fit to an exponential growth equation to determine the specific growth rate for each condition and sample (**Methods**). Specific growth rates are compared for each dCas induction level (aTc concentration) with the results of a one-way ANOVA with Tukey post-hoc analysis reported as not significant (ns) or significant (*$p < 0.05$, **$p < 0.01$, and ***$p < 0.001$). Calculated values are given in **SI File 1**.

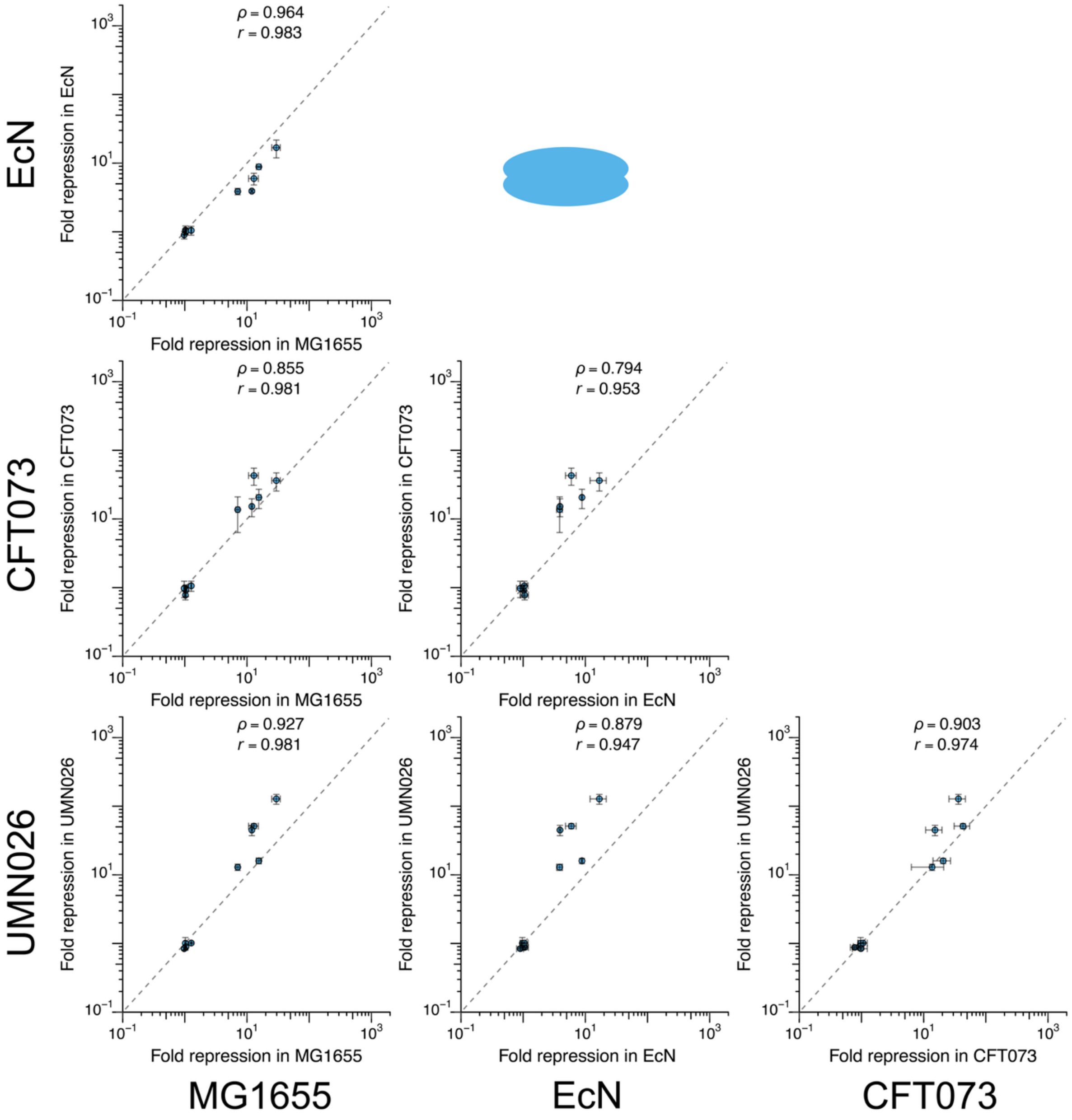

**Figure S14. Correlations of repression for Lb-dCas12a CRISPRi between strains**

The average fold repression values for all gRNA designs of the Lb-dCas12a system for all pairwise combinations of MG1655, EcN, CFT073, and UMN026 strains are shown. The diagonal y = x (dashed line) is included for comparison. Pearson (*r*) and Spearman (*ρ*) correlations were performed on the $\log_{10}$-transformed fold repression values of each dataset and are indicated. The *p*-value for each correlation is provided in **SI File 1**.

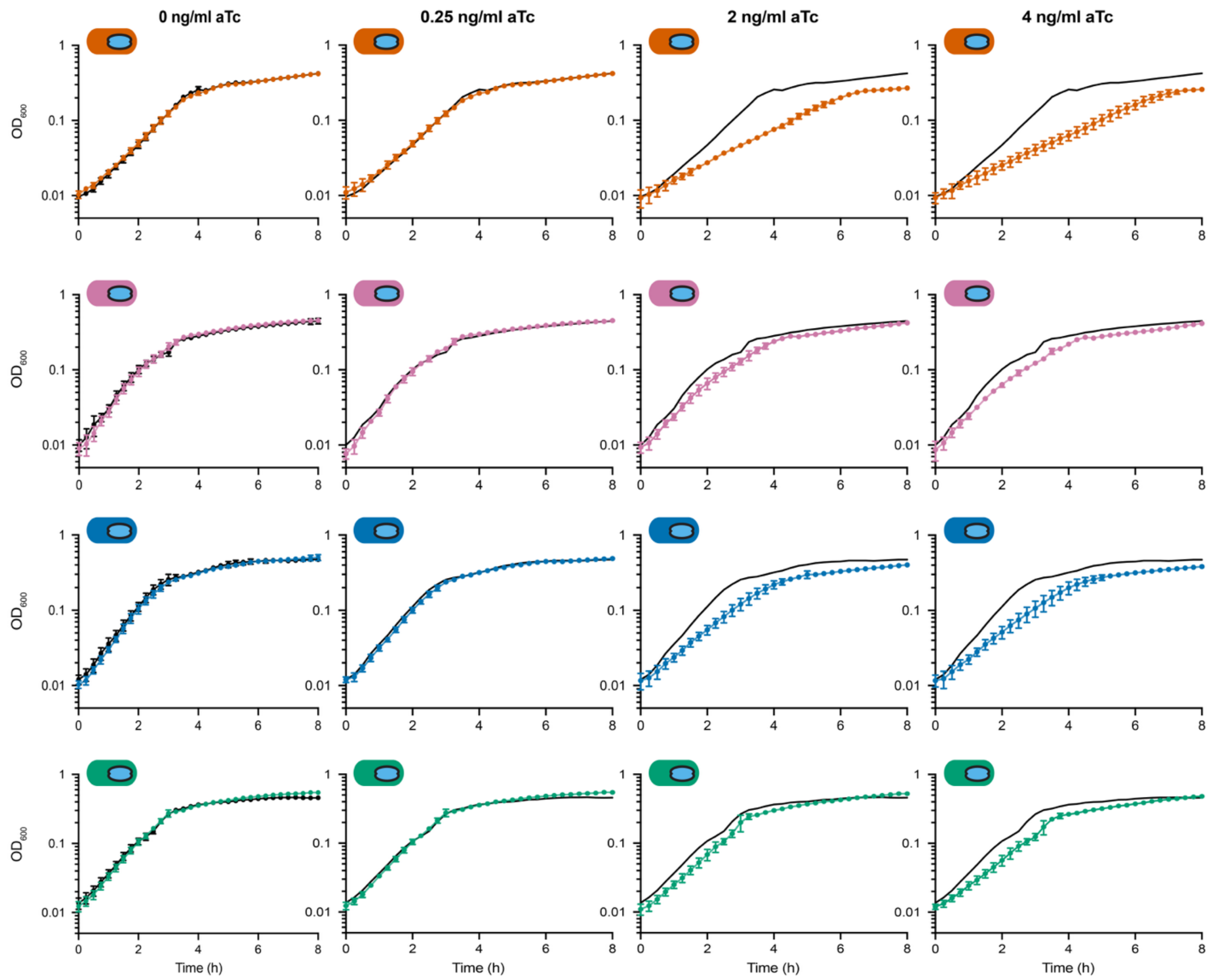

**Figure S15. Growth curves for Lb-dCas12a expression in *E. coli* strains**

Growth curves were performed for MG1655 (orange), EcN (pink), CFT073 (blue), and UMN026 (green) at different induction levels for expression of Lb-dCas12a in plasmid pSR2032 via the indicated concentration of aTc in the media (0, 0.25, 2, and 4 ng/mL aTc shown). Absorbance at 600 nm ($OD_{600}$) was measured at 15-minute intervals during an 8-hour incubation in a microplate reader (**Methods**). Points indicate the mean of three identical experiments performed on three different days, and error bars are the standard deviation. The growth curves for the wildtype cells for each strain (WT, black) without any inducer are shown for comparison in each plot with the error bars for the wildtype only drawn in the first column (0 ng/mL aTc).

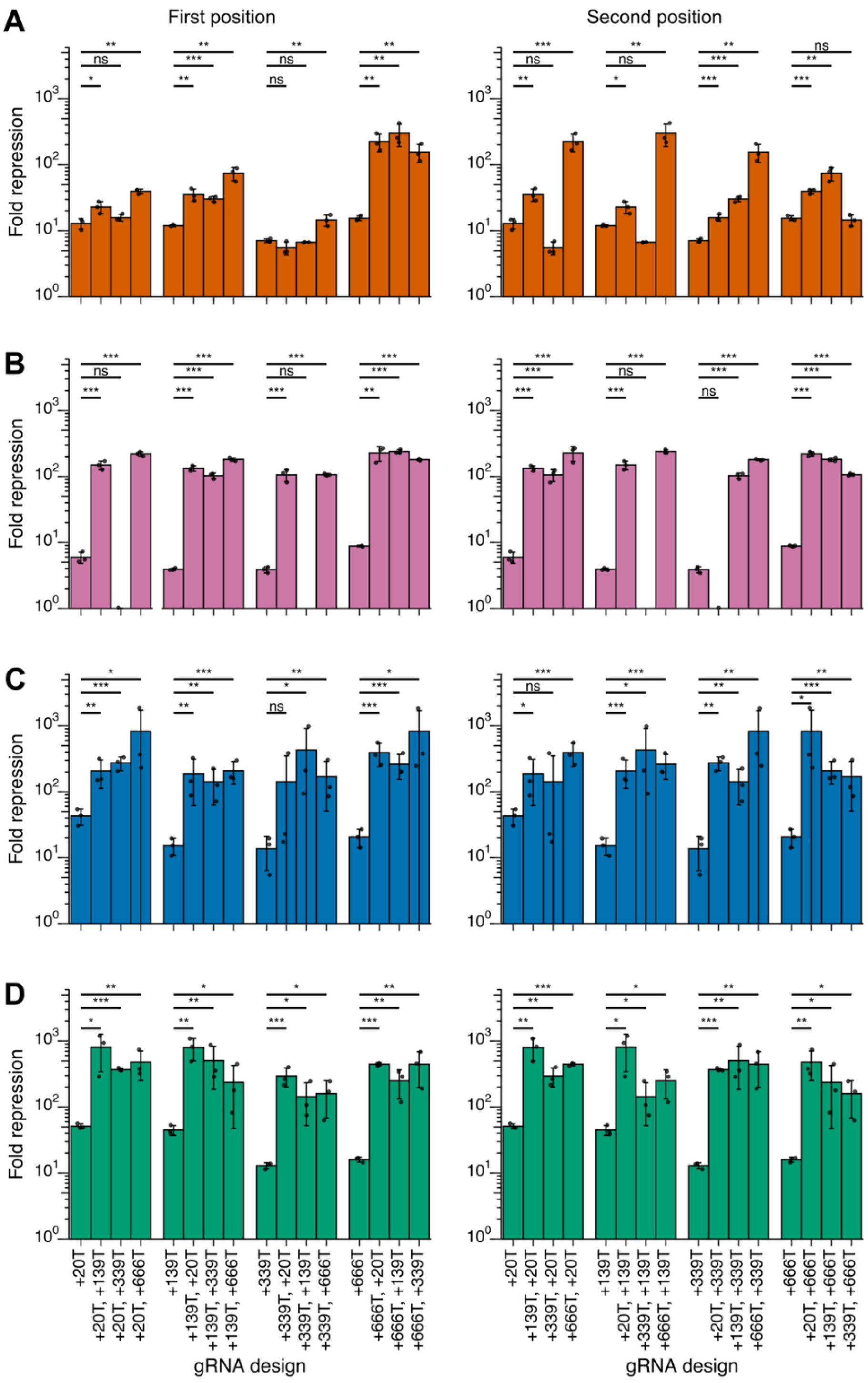

**Figure S16. Comparison of single and dual gRNA for Lb-dCas12a CRISPRi**

Measured repression of each dual gRNA is plotted next to the corresponding single gRNA in the (left) first position and (right) second position in **(A)** MG1655, **(B)** EcN, **(C)** CFT073, and **(D)** UMN026. Results of Welch's one-sided, unpaired *t*-tests are shown as not significant (ns) or significant (*$p < 0.05$, **$p < 0.01$, and ***$p < 0.001$), and values reported in **SI File 1**. The average ± SD of three identical experiments (points) are plotted.

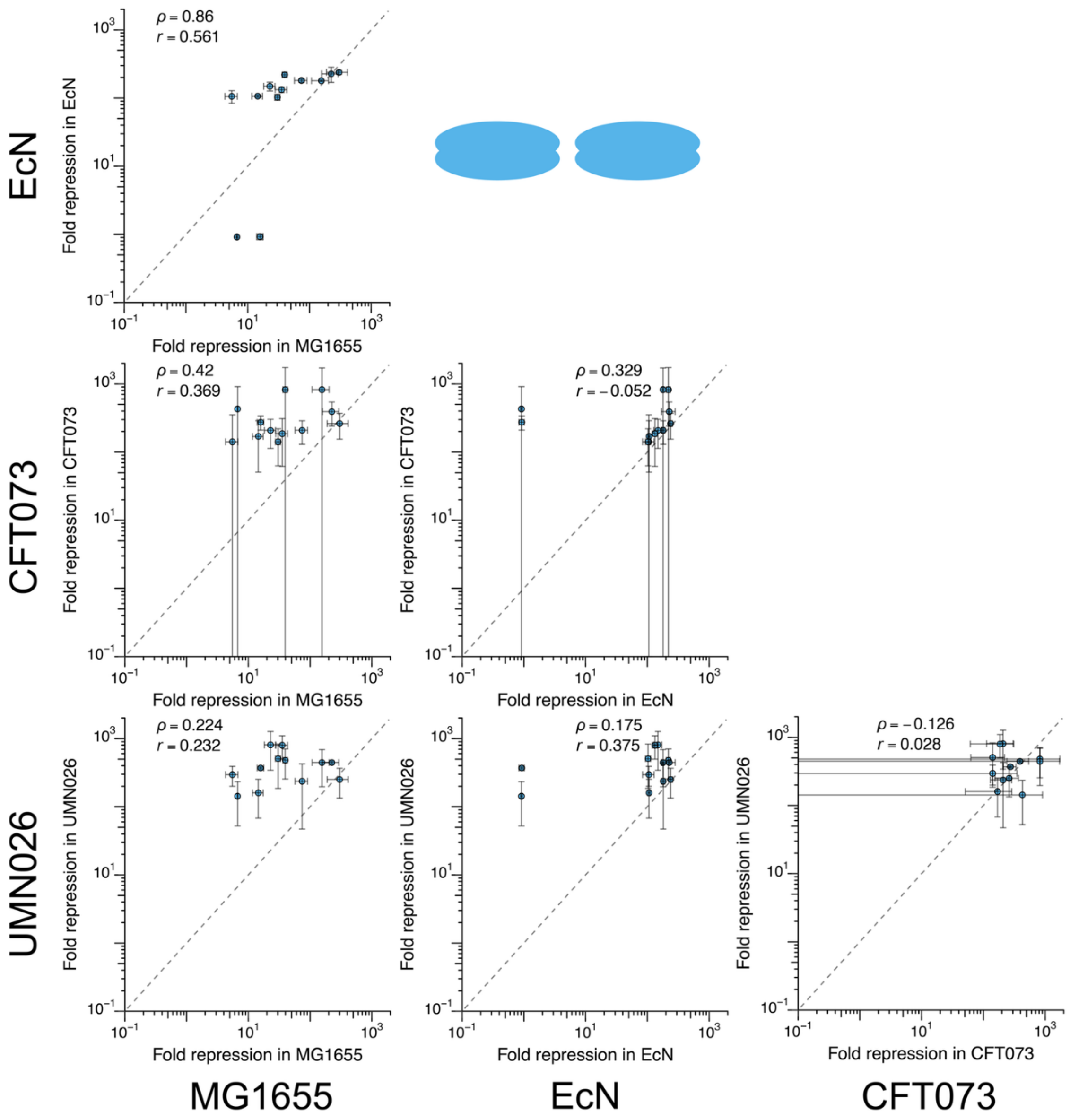

**Figure S17. Correlations of repression for Lb-dCas12a CRISPRi with dual gRNA arrays**

The average fold repression values for all dual gRNA designs of the Lb-dCas12a system for all pairwise combinations of MG1655, EcN, CFT073, and UMN026 strains are shown. The diagonal y = x (dashed line) is included for comparison. Pearson (*r*) and Spearman (*ρ*) correlations were performed on the $\log_{10}$-transformed fold repression values of each dataset and are indicated. The *p*-value for each correlation is provided in **SI File 1**.

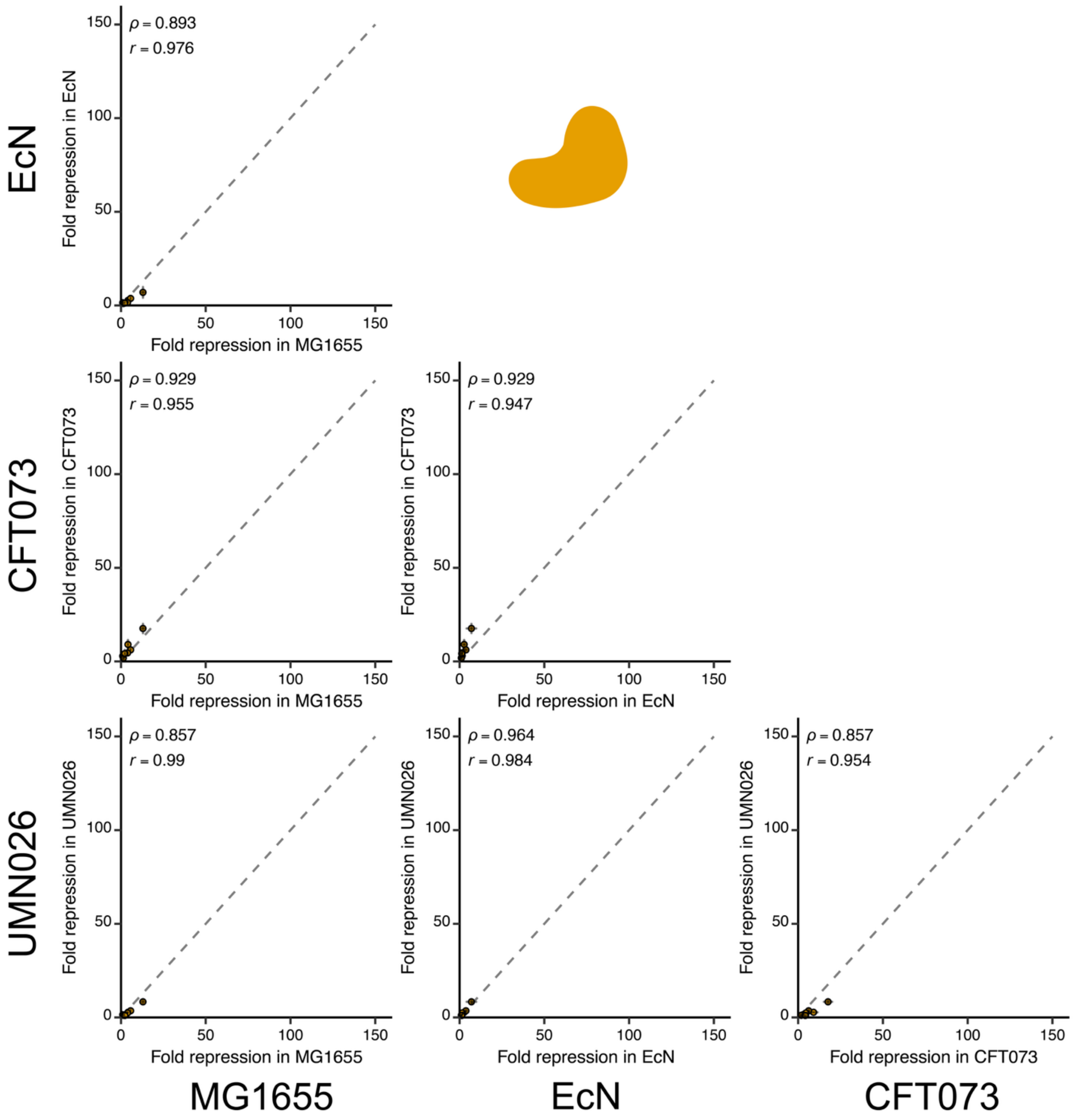

**Figure S18. Repression with Sp-dCas9 and expected active gRNA from prior studies**

The average fold repression values for gRNA targeting either strand of the promoter region (–123N, –89N, –144T, –128T) or non-template strand of the coding sequence (+34N, +334N, +664N) of *eyfp* using Sp-dCas9 CRISPRi for all pairwise combinations of MG1655, EcN, CFT073, and UMN026 strains are shown. The diagonal y = x (dashed line) is included for comparison. Pearson (*r*) and Spearman (*ρ*) correlations were performed on the untransformed fold repression values of each dataset and are indicated. The *p*-value for each correlation is in **SI File 1**.

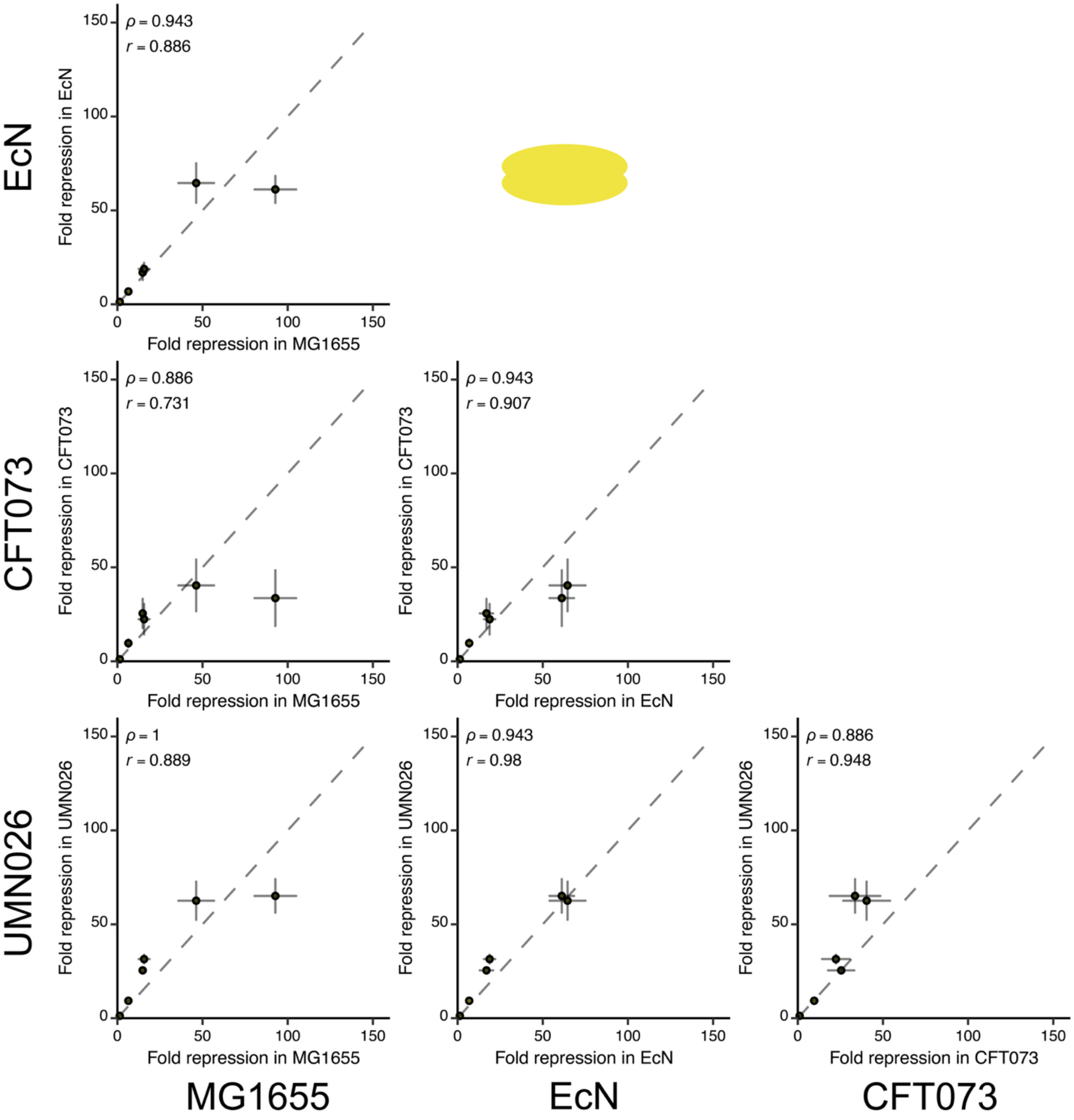

**Figure S19. Repression with Fn-dCas12a and expected active gRNA from prior studies**

The observed average fold repression for gRNA targeting the promoter region (–119N, –175T) or template strand of the coding sequence (+20T, +139T, +339T, +666T) of *sfgfp* using Fn-dCas12a for all pairwise combinations of MG1655, EcN, CFT073, and UMN026 strains are shown. The diagonal y = x (dashed line) is included for comparison. Pearson ($r$) and Spearman ($\rho$) correlations were performed on the untransformed fold repression values of each dataset and are indicated. The $p$-value for each correlation is provided in **SI File 1**.

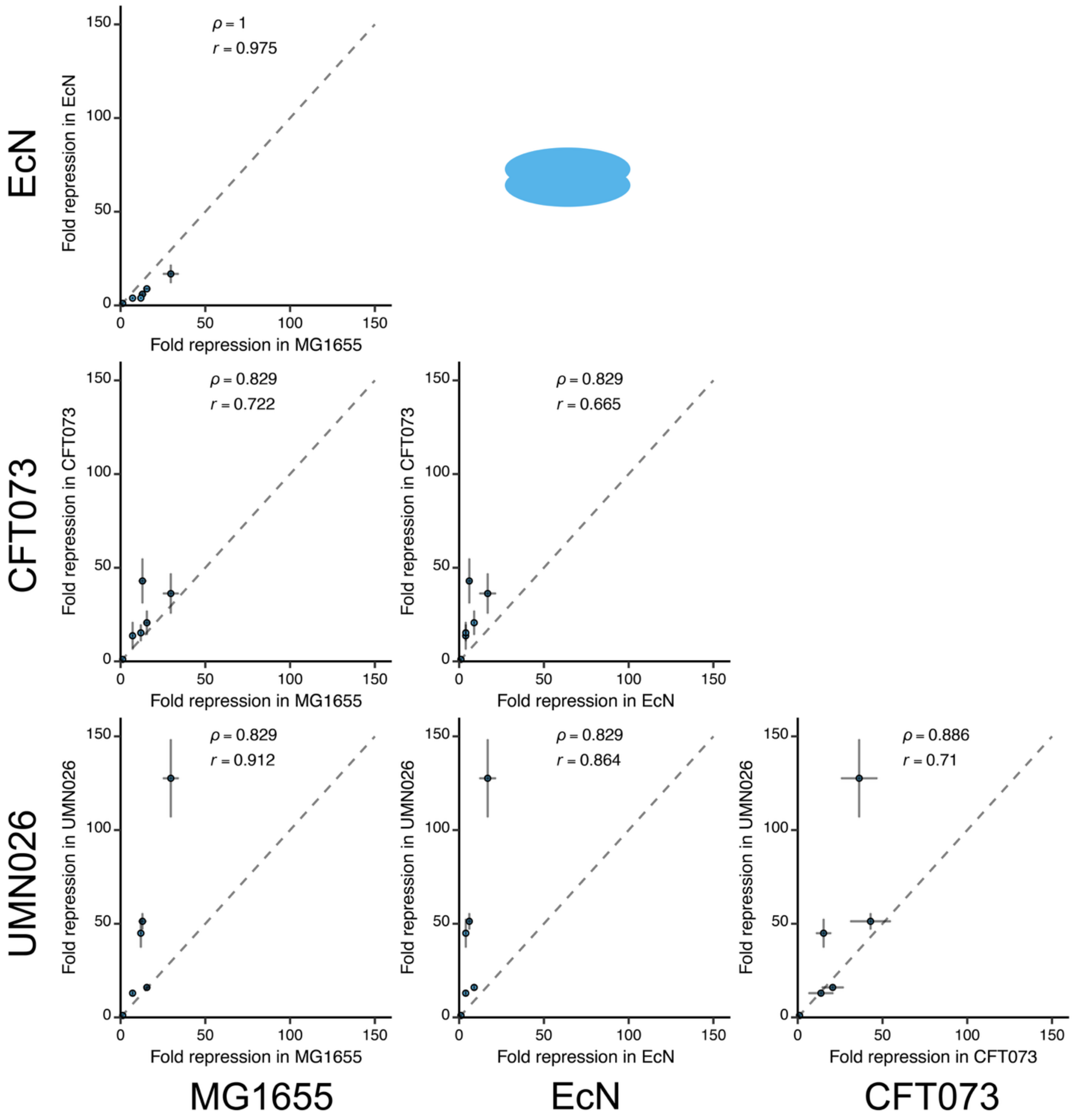

**Figure S20. Repression with Lb-dCas12a and expected active gRNA from prior studies**

The average fold repression values for gRNA targeting the promoter region (–119N, –175T) or template strand of the coding sequence (+20T, +139T, +339T, +666T) of *sfgfp* using Lb-dCas12a CRISPRi for all pairwise combinations of MG1655, EcN, CFT073, and UMN026 strains are shown. The diagonal y = x (dashed line) is included for comparison. Pearson ($r$) and Spearman ($\rho$) correlations were performed on the untransformed fold repression values of each dataset and are indicated. The $p$-value for each correlation is provided in **SI File 1**.

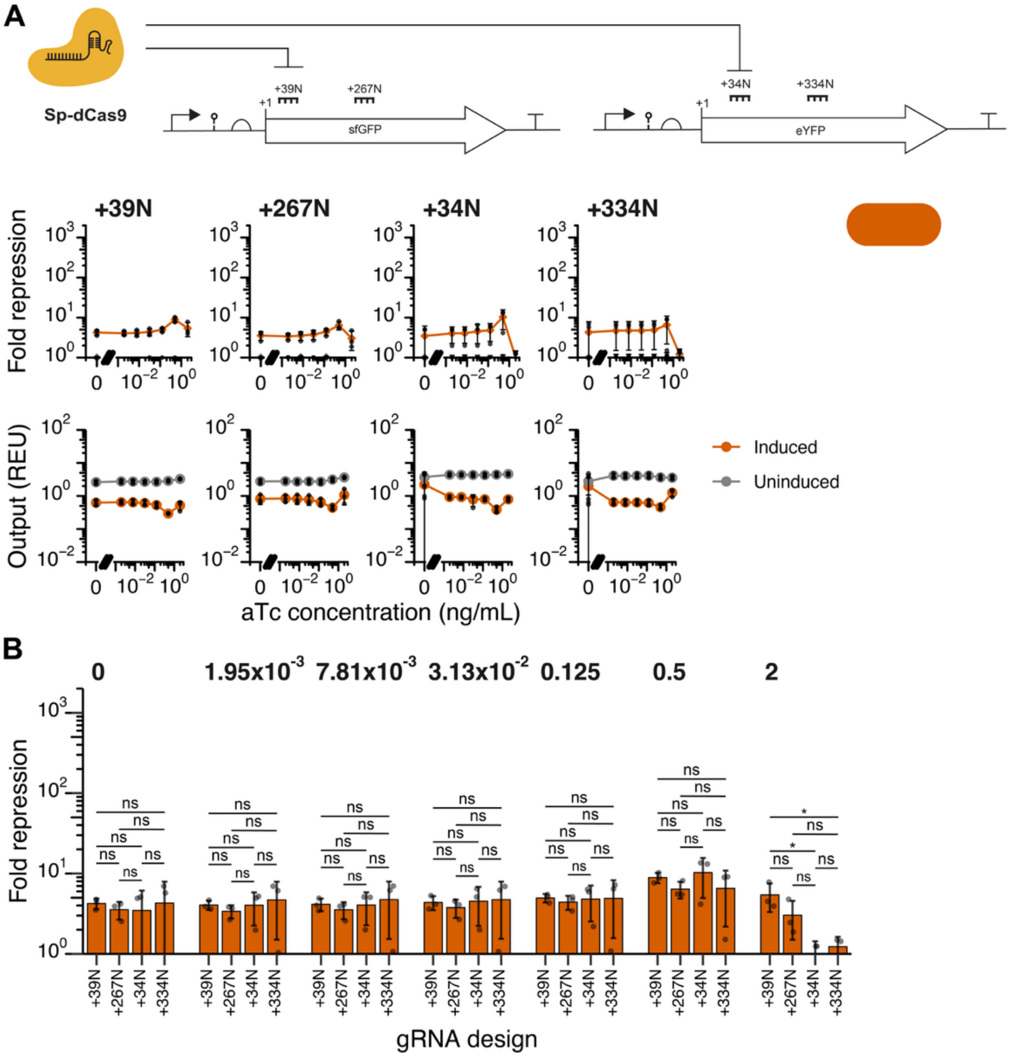

**Figure S21. Titrations of Sp-dCas9 with gRNAs targeting *sfgfp* in MG1655**

**(A)** Binding locations of gRNAs designed to target the non-template (N) strand of the coding sequence of *sfgfp* and *eyfp* using Sp-dCas9. Naming of gRNAs indicates the targeted strand and position relative to the gene start (+1). The expression of Sp-dCas9 in MG1655 was titrated by adding varying aTc in the media with the gRNA either induced with 1 mM IPTG (orange) or uninduced without IPTG (gray). The cell fluorescence was measured using flow cytometry and arbitrary units converted to relative expression units (REU). Fold repression was calculated as the ratio of the output without and with induction of both Sp-dCas9 and the gRNA. **(B)** The repression values for these designs are compared at each aTc concentration listed at top (in ng/mL), with the results of a one-way ANOVA with Tukey post-hoc analysis shown as not significant (ns) or significant (*$p < 0.05$, **$p < 0.01$, and ***$p < 0.001$). Specific $p$-values are given in **SI File 1**. Bars are the arithmetic mean of three identical experiments (small points) performed on three different days with error bars depicting the standard deviation.

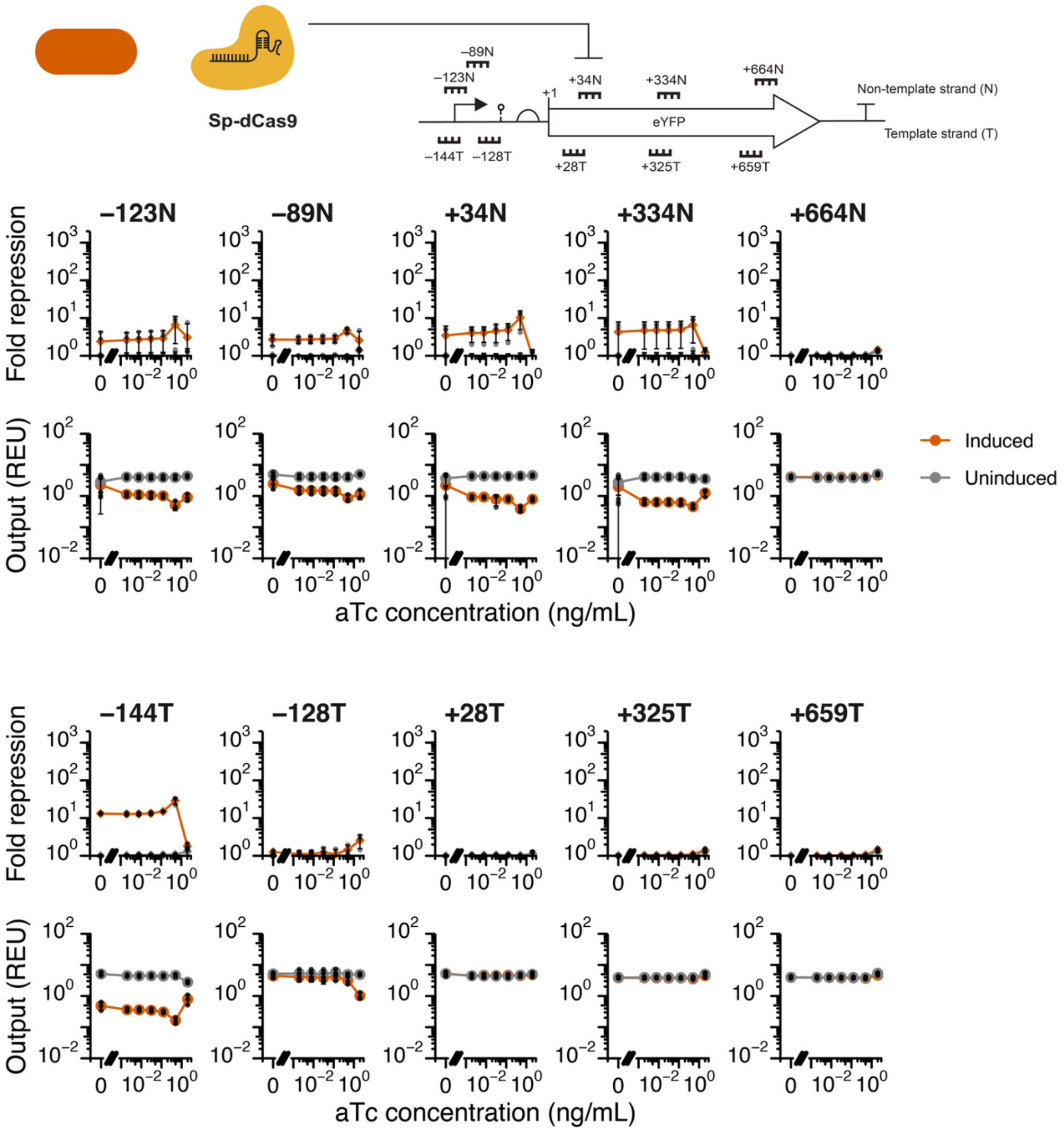

**Figure S22. Titrations of Sp-dCas9 with gRNAs targeting *eyfp* in MG1655**

Genetic schematic illustrates the binding locations of the gRNA designed to target *eyfp* using Sp-dCas9. Name of the gRNA indicates the targeted location relative to the gene start of *eyfp* (+1) and whether the non-template (N) or template (T) strand is targeted. The expression of Sp-dCas9 in MG1655 was titrated by adding varying amounts of aTc into the media with the gRNA either induced with 1 mM IPTG (orange) or uninduced without IPTG (gray). The cell fluorescence was measured using flow cytometry and arbitrary units converted to relative expression units (REU). Fold repression was calculated as the ratio of the output without and with induction of both Sp-dCas9 and the gRNA. Markers are the arithmetic mean of three identical experiments performed on three different days, and error bars show the standard deviation. Lines connect adjacent points of the same titration.

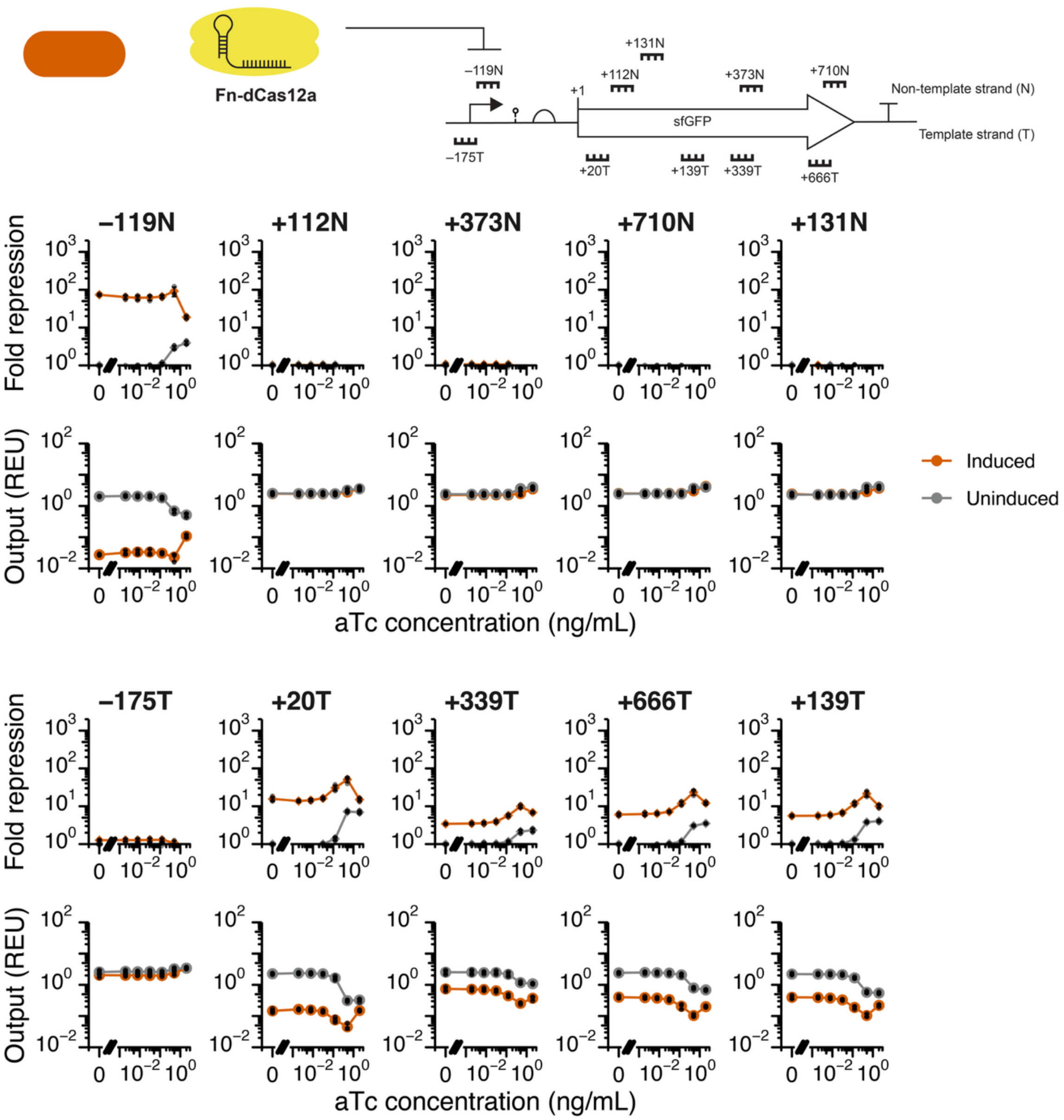

**Figure S23. Titrations of Fn-dCas12a with gRNAs targeting *sfgfp* in MG1655**

Genetic schematic illustrates the binding locations of the gRNA designed to target *sfgfp* using Fn-dCas12a. Name of the gRNA indicates the targeted location relative to the gene start of *sfgfp* (+1) and whether the non-template (N) or template (T) strand is targeted. The expression of Fn-dCas12a in MG1655 was titrated by adding varying amounts of aTc into the media with the gRNA either induced with 1 mM IPTG (orange) or uninduced without IPTG (gray). The cell fluorescence was measured using flow cytometry and arbitrary units converted to relative expression units (REU). Fold repression was calculated as the ratio of the output without and with induction of both Fn-dCas12a and the gRNA. Markers are the arithmetic mean of three identical experiments performed on three different days, and error bars show the standard deviation. Lines connect adjacent points of the same titration.

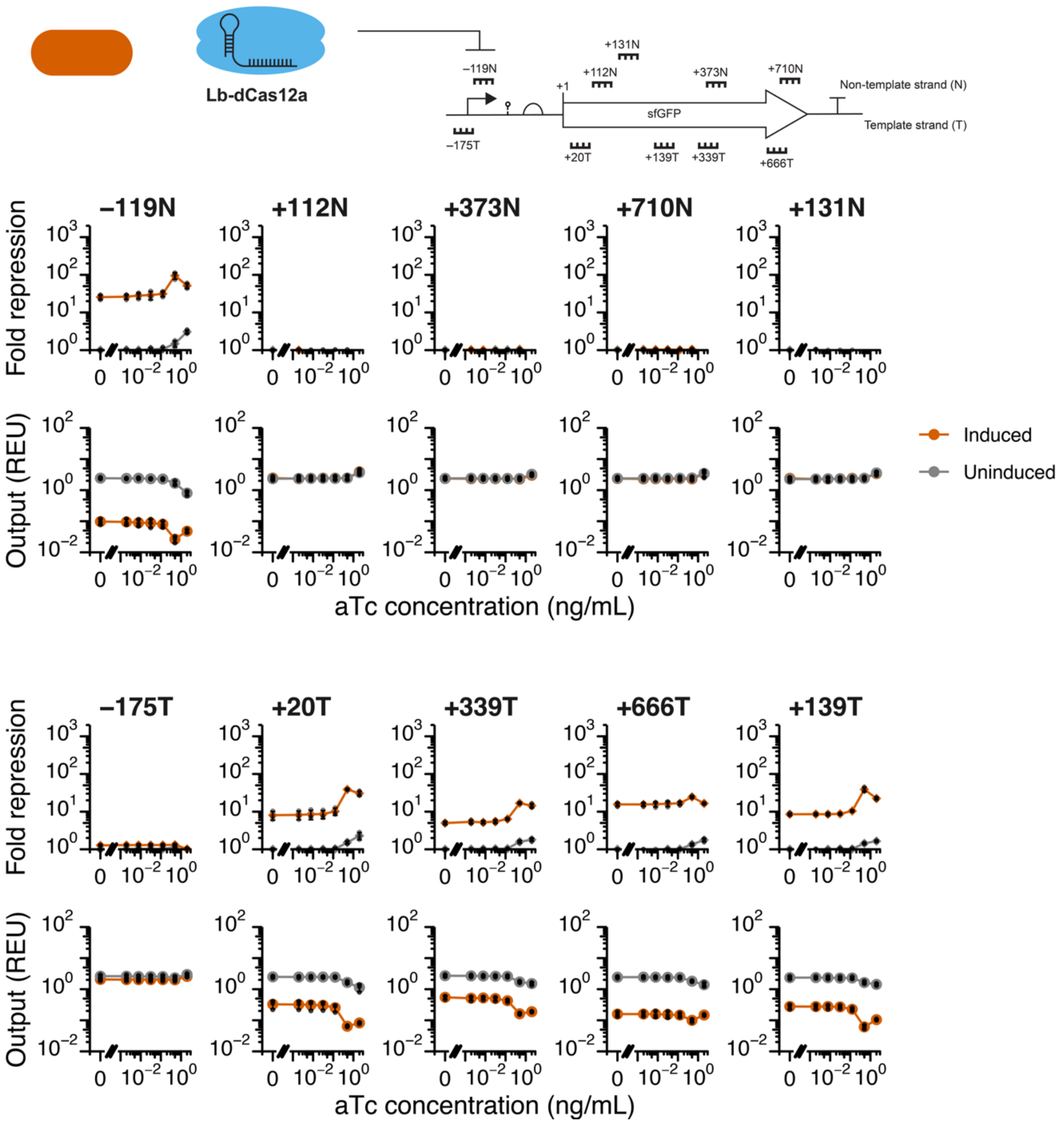

**Figure S24. Titrations of Lb-Cas12a with gRNAs targeting *sfgfp* in MG1655**

Genetic schematic illustrates the binding locations of the gRNA designed to target *sfgfp* using Lb-dCas12a. Name of the gRNA indicates the targeted location relative to the gene start of *sfgfp* (+1) and whether the non-template (N) or template (T) strand is targeted. The expression of Lb-dCas12a in MG1655 was titrated by adding varying amounts of aTc into the media with the gRNA either induced with 1 mM IPTG (orange) or uninduced without IPTG (gray). The cell fluorescence was measured using flow cytometry and arbitrary units converted to relative expression units (REU). Fold repression was calculated as the ratio of the output without and with induction of both Lb-dCas12a and the gRNA. Markers are the arithmetic mean of three identical experiments performed on three different days, and error bars show the standard deviation. Lines connect adjacent points of the same titration.

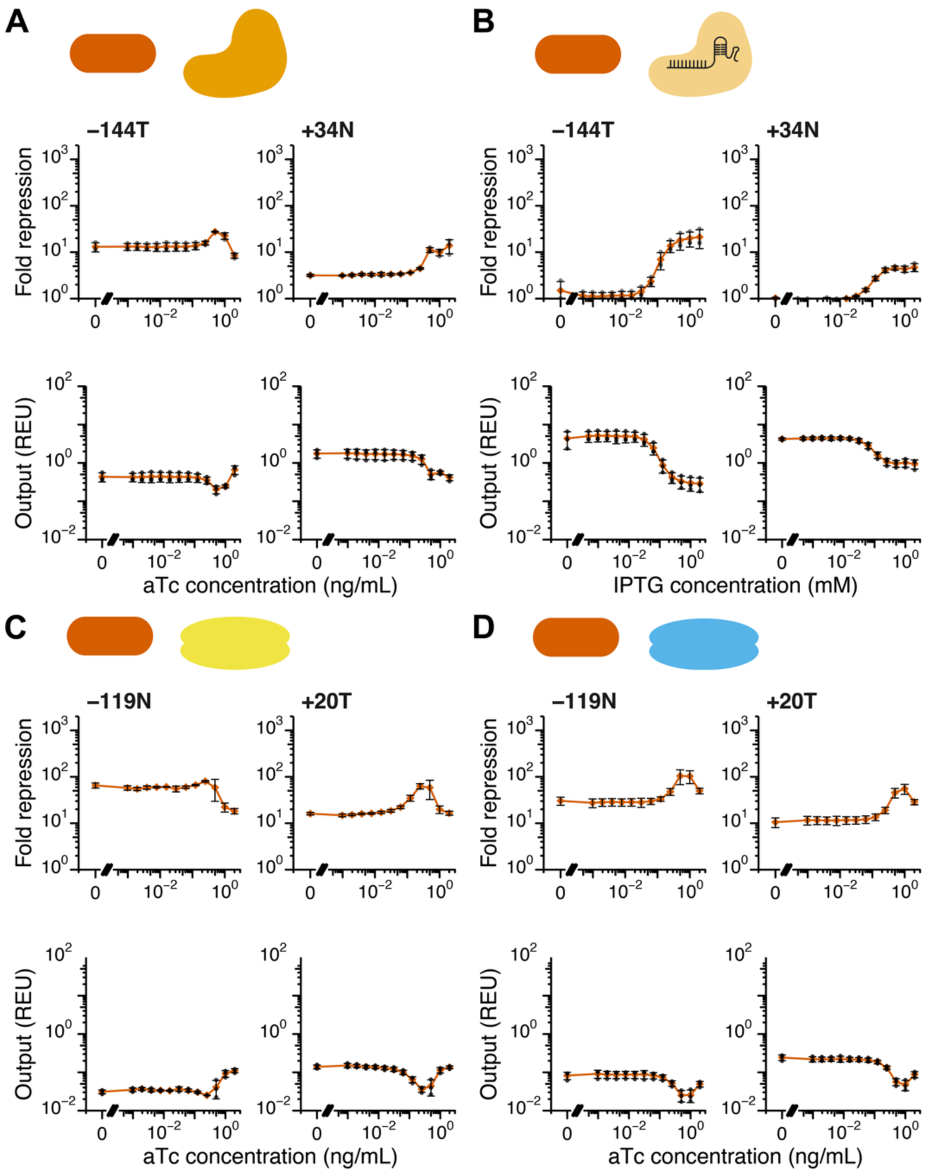

**Figure S25. Higher resolution titrations of select CRISPRi components in MG1655**

Two-fold titrations of the expression of **(A)** Sp-dCas9, **(B)** gRNA designed for Sp-dCas9, **(C)** Fn-dCas12a, and **(D)** Lb-dCas12a CRISPRi were performed in MG1655 with the indicated gRNA designs by adding varying amounts of aTc or IPTG for the dCas or gRNA induction, respectively. Cell fluorescence was measured using flow cytometry and arbitrary units converted to relative expression units (REU). Fold repression was calculated as the ratio of the output without and with induction of both the dCas protein and the gRNA. gRNA were chosen as the designs that showed the greatest repression in MG1655 targeting either the promoter or coding sequence for each CRISPRi system. Markers are the arithmetic mean of three identical experiments (small black points) performed on three different days with error bars depicting the standard deviation. Lines connect adjacent points of the same titration.

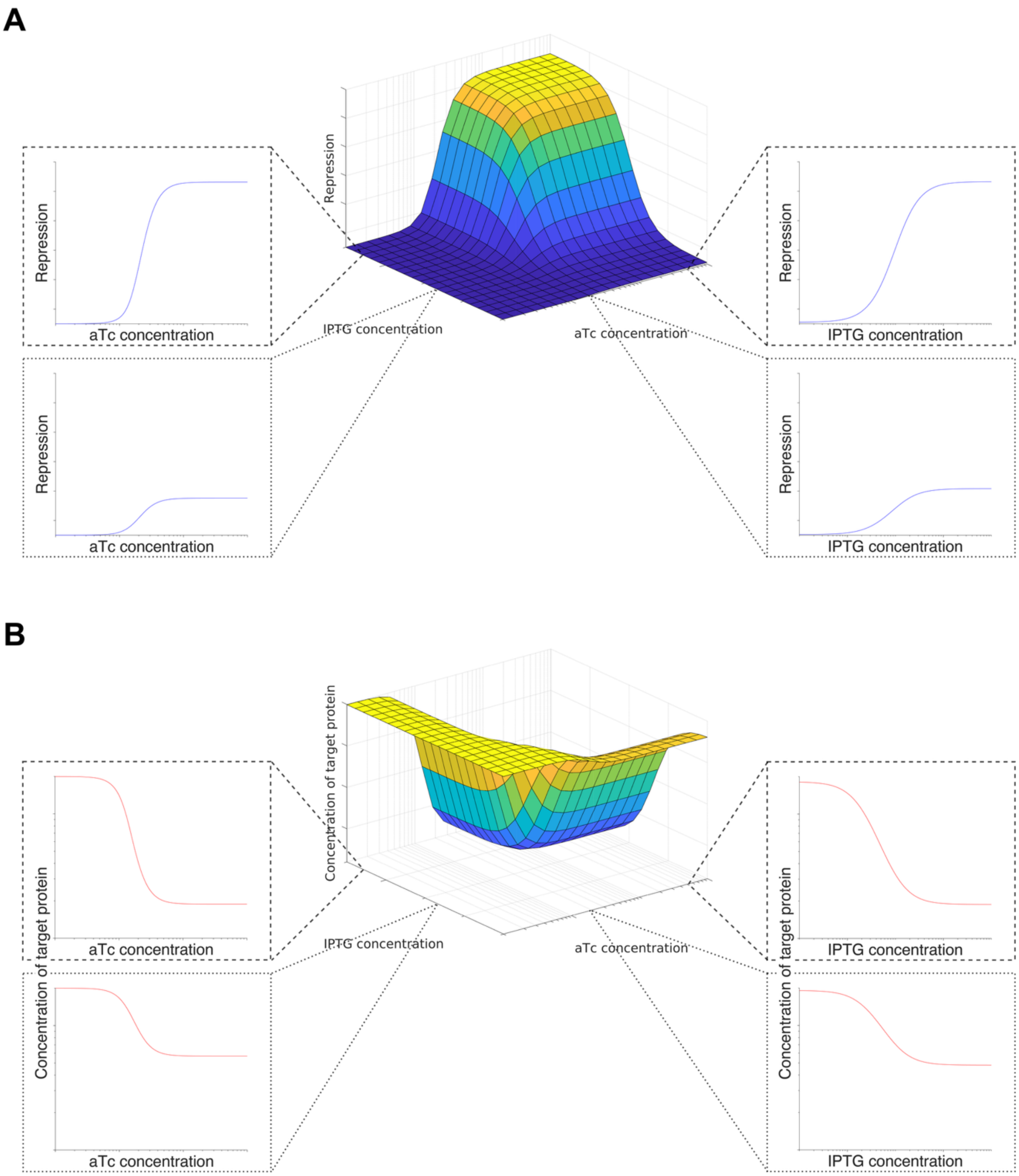

**Figure S26. Simulations of Sp-dCas9 repression using a kinetic model**

A kinetic model was created for our genetic design for the Sp-dCas9 system (**Extended Methods**). Simulations were performed by varying concentrations of aTc and IPTG inducers to tune expression of Sp-dCas9 and the gRNA, respectively. The resulting surface plots are given for **(A)** fold repression and **(B)** the concentration of target protein. Planes are provided at select concentrations of each inducer to illustrate the change in the fold repression and target protein concentration at varying concentrations of one inducer with the other held constant, as was performed in MG1655 using flow cytometry.

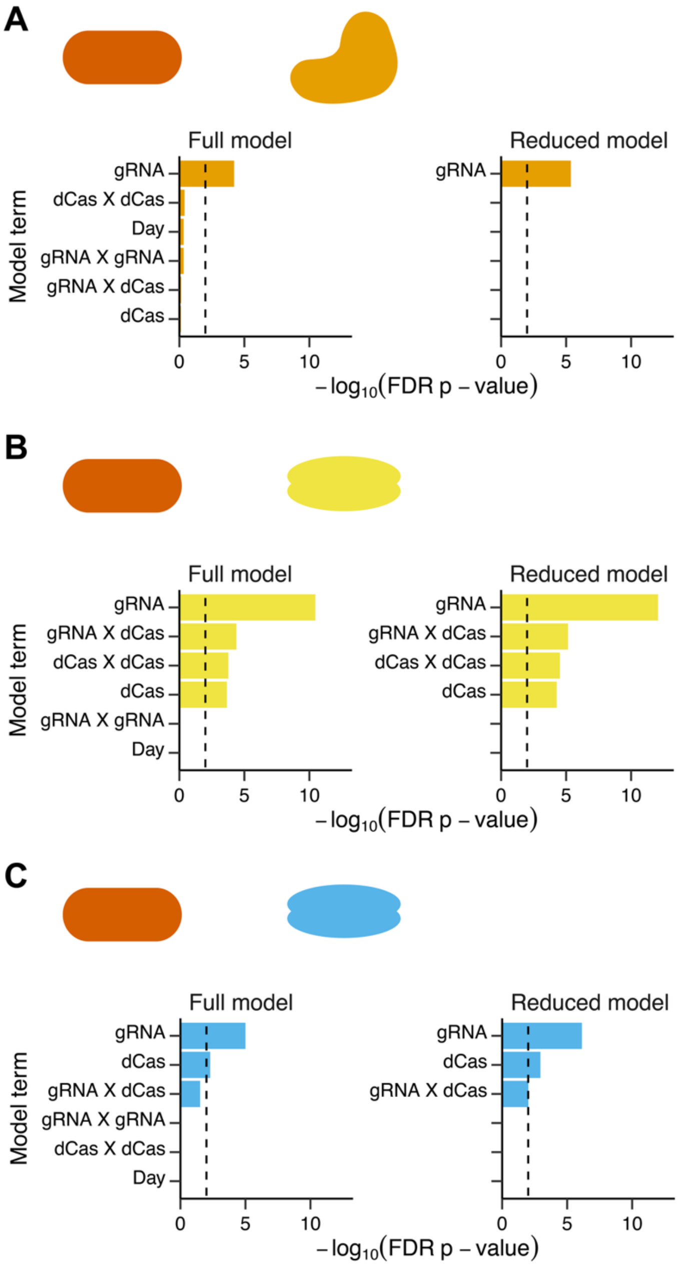

**Figure S27. Design of experiments model for CRISPRi repression in MG1655**

Design of experiments analysis was performed in MG1655 at varying concentrations of **(A)** Sp-dCas9, **(B)** Fn-dCas12a, and **(C)** Lb-dCas12a and the gRNA demonstrating the greatest repression for each system (**Extended Methods**). A polynomial model, including interaction terms and a blocking term for the day of the experiment, was fit to the data collected for each CRISPRi system using JMP Pro v17.0. The false discovery rate (FDR) for each term in the full model and reduced model is given. The reduced model was determined by systematically removing the term with the lowest significance until all remaining terms were significant (FDR < 0.01, dashed line). Estimated terms and *p*-values for the full and reduced models are given in **Tables S5 and S6**.

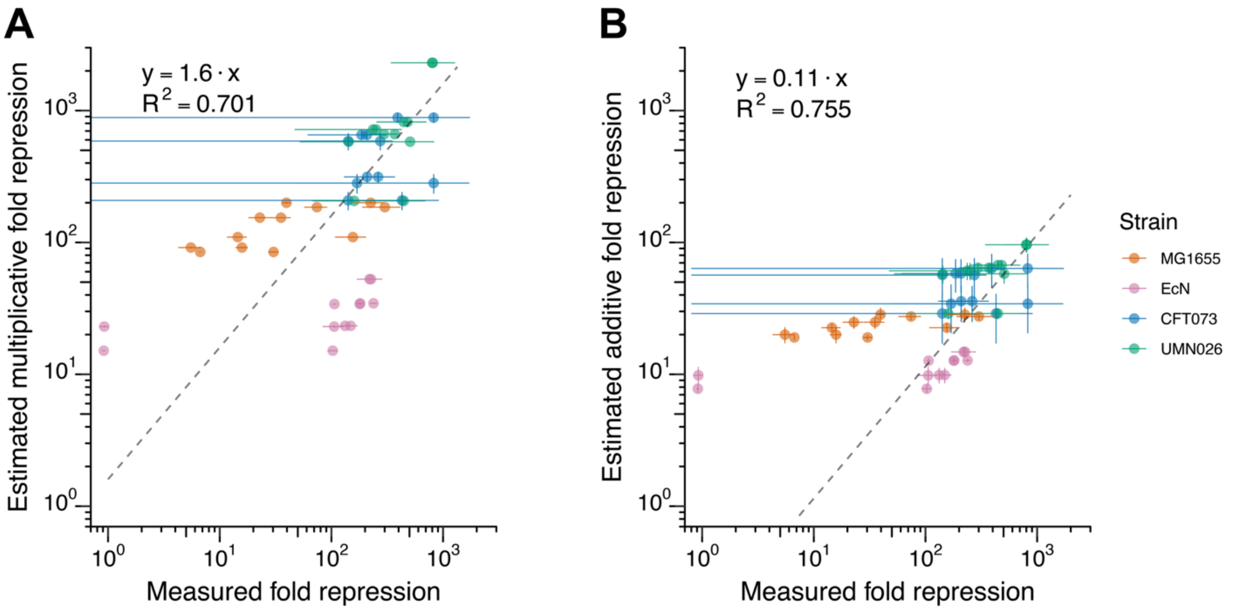

**Figure S28. Analysis of epistasis models for dual gRNA arrays with Lb-dCas12a**

The predicted epistatic effects based on a **(A)** multiplicative and **(B)** additive model were calculated for the fold repression of dual gRNA designs for Lb-dCas12a in each strain by multiplying or adding the fold repression of the Lb-dCas12a single gRNA comprising the designs, respectively. The resulting values are plotted against the measured fold repression through flow cytometry for all strains. A linear fit through the origin (dashed line) of the mean values was performed for each dataset, and the resulting equation and coefficient of determination ($R^2$) given. All flow data are the arithmetic mean of three identical experiments performed on separate days with horizontal error bars depicting the standard deviation. Vertical error bars for the epistatic calculations were determined through error propagation of the standard deviation and mean for both single gRNA comprising the dual gRNA design (**Extended Methods**).

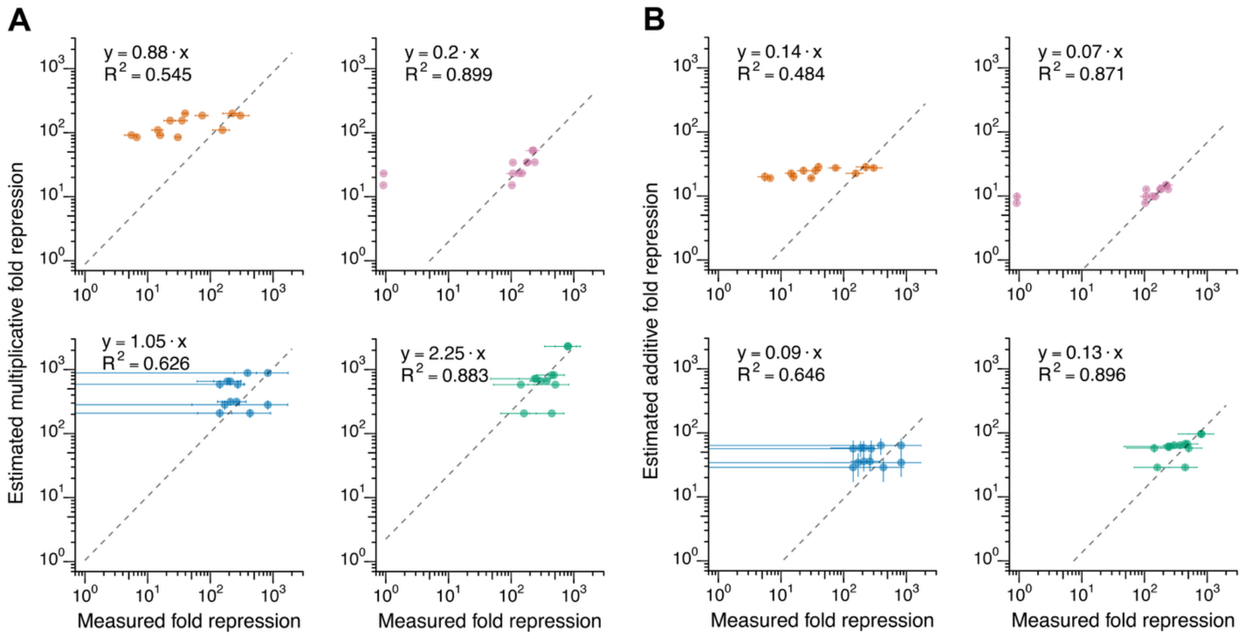

**Figure S29. Epistatic effects for dual gRNA arrays with Lb-dCas12a**

In each strain, the predicted epistatic effects based on a **(A)** multiplicative and **(B)** additive model were calculated for the fold repression dual gRNA designs for Lb-dCas12a by multiplying or adding the fold repression of the Lb-dCas12a single gRNA comprising the designs, respectively. The resulting values are plotted against the measured fold repression from flow cytometry for each strain. A linear fit through the origin (dashed line) of the mean values was performed for each dataset, and the resulting equation and coefficient of determination ($R^2$) given. All flow data are the arithmetic mean of three identical experiments performed on separate days with horizontal error bars depicting the standard deviation. Vertical error bars for the epistatic calculations were determined through error propagation of the standard deviation and mean for both single gRNA comprising the dual gRNA design (**Extended Methods**).

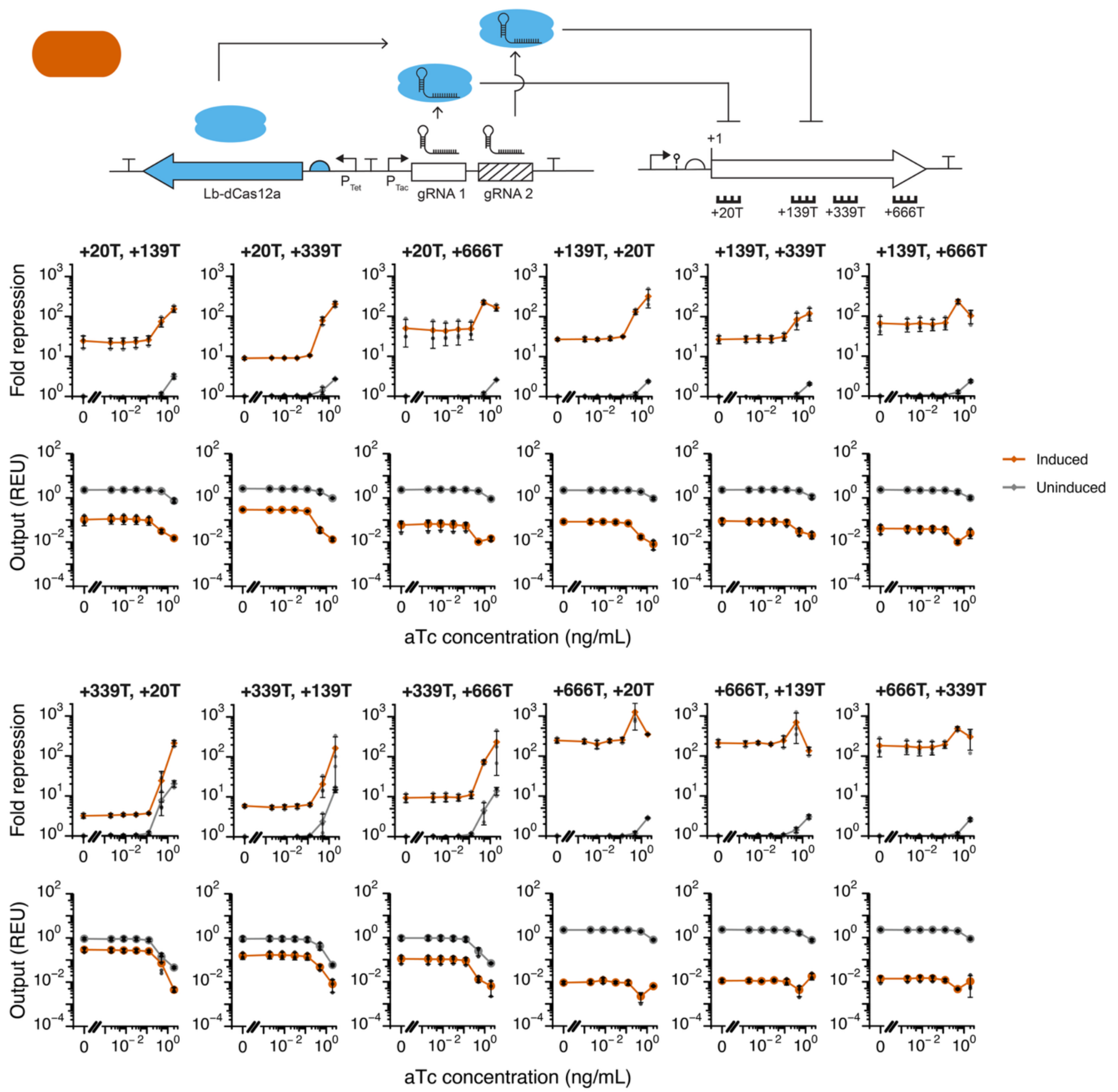

**Figure S30. Titrations of Lb-Cas12a with dual gRNAs targeting *sfgfp* in MG1655**

A genetic schematic illustrates the binding locations of the subset of single gRNA designs targeting *sfgfp* that were used to create dual gRNA designs for Lb-Cas12a. The expression of Lb-dCas12a in MG1655 was titrated by adding varying amounts of aTc into the media with the gRNA either induced with 1 mM IPTG (orange) or uninduced without IPTG (gray). Cell fluorescence was measured using flow cytometry and arbitrary units converted to relative expression units (REU). Fold repression was calculated as the ratio of the output without and with induction of both Lb-dCas12a and the gRNA. Markers are the arithmetic mean of three identical experiments (small black points) performed on three different days ($n$ > 5,000 events per sample) with error bars depicting the standard deviation. Lines connect adjacent points of the same titration.

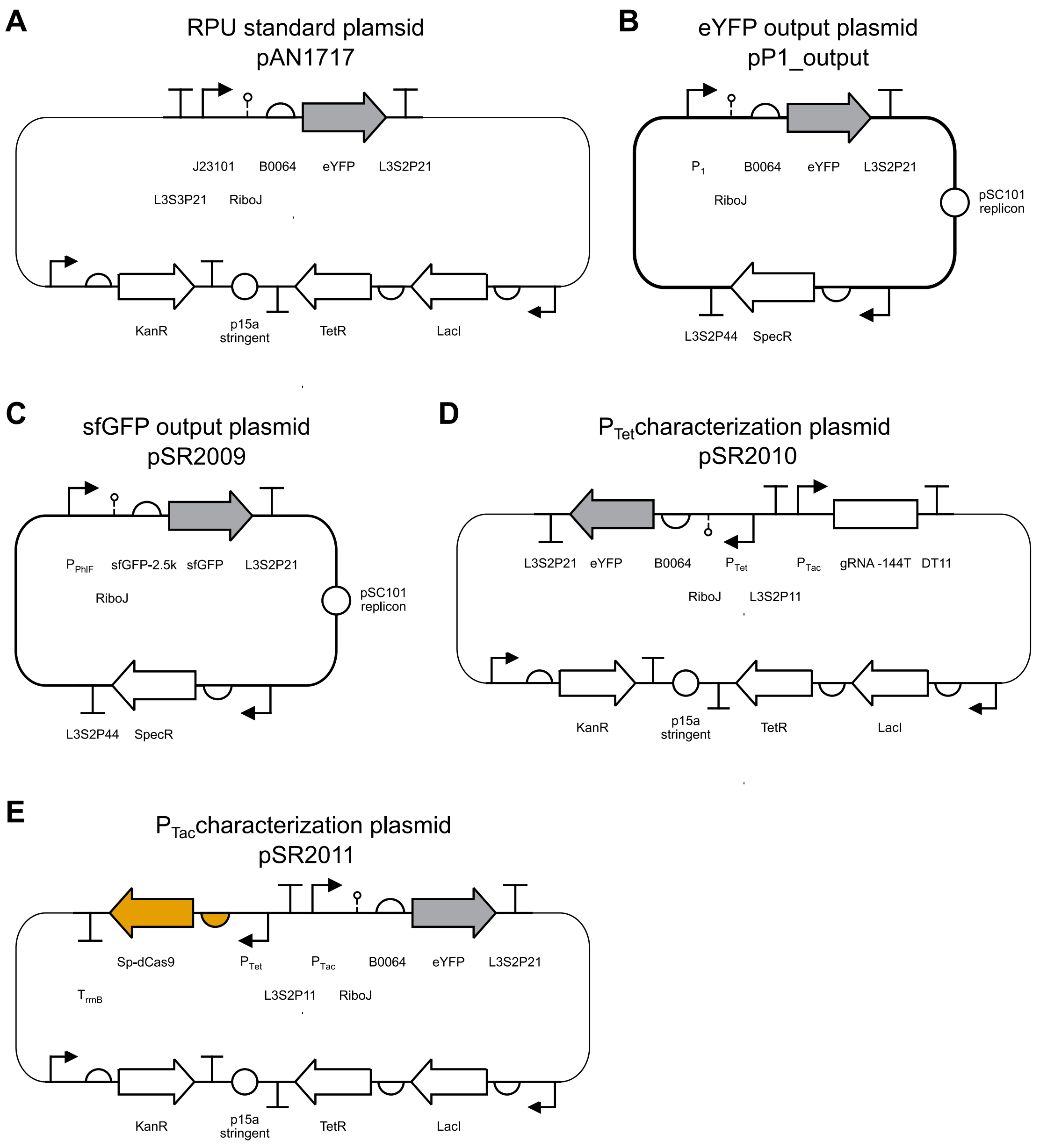

**Figure S31. Plasmid maps of standard, output, and promoter characterization plasmids**

**(A)** Plasmid pAN1717,[2,3] which contains a constitutive *eyfp* cassette, was used for relative promoter unit (RPU) and relative expression unit (REU) determination. **(B)** Plasmid pP1_output contains an *eyfp* gene on a low-copy vector (pSC101 origin) and was used as the output plasmid for Sp-dCas9 CRISPRi. **(C)** pSR2009 contains *sfgfp* on a low-copy vector (pSC101 origin) and was used as the output plasmid for Fn-dCas12a and Lb-dCas12a CRISPRi. **(D)** pSR2010 was used to perform sensor characterization and determine the response function for $P_{Tet}$. It is identical to the CRISPRi plasmid (with gRNA –144T), except the dCas gene was replaced by *eyfp* regulated by $P_{Tet}$. **(E)** pSR2011 was used to determine the response function for $P_{Tac}$. It is identical to the CRISPRi plasmid (with Sp-dCas9), except the gRNA was replaced by *eyfp* regulated by $P_{Tac}$. Details for each of these plasmids are provided in **Table S7**.

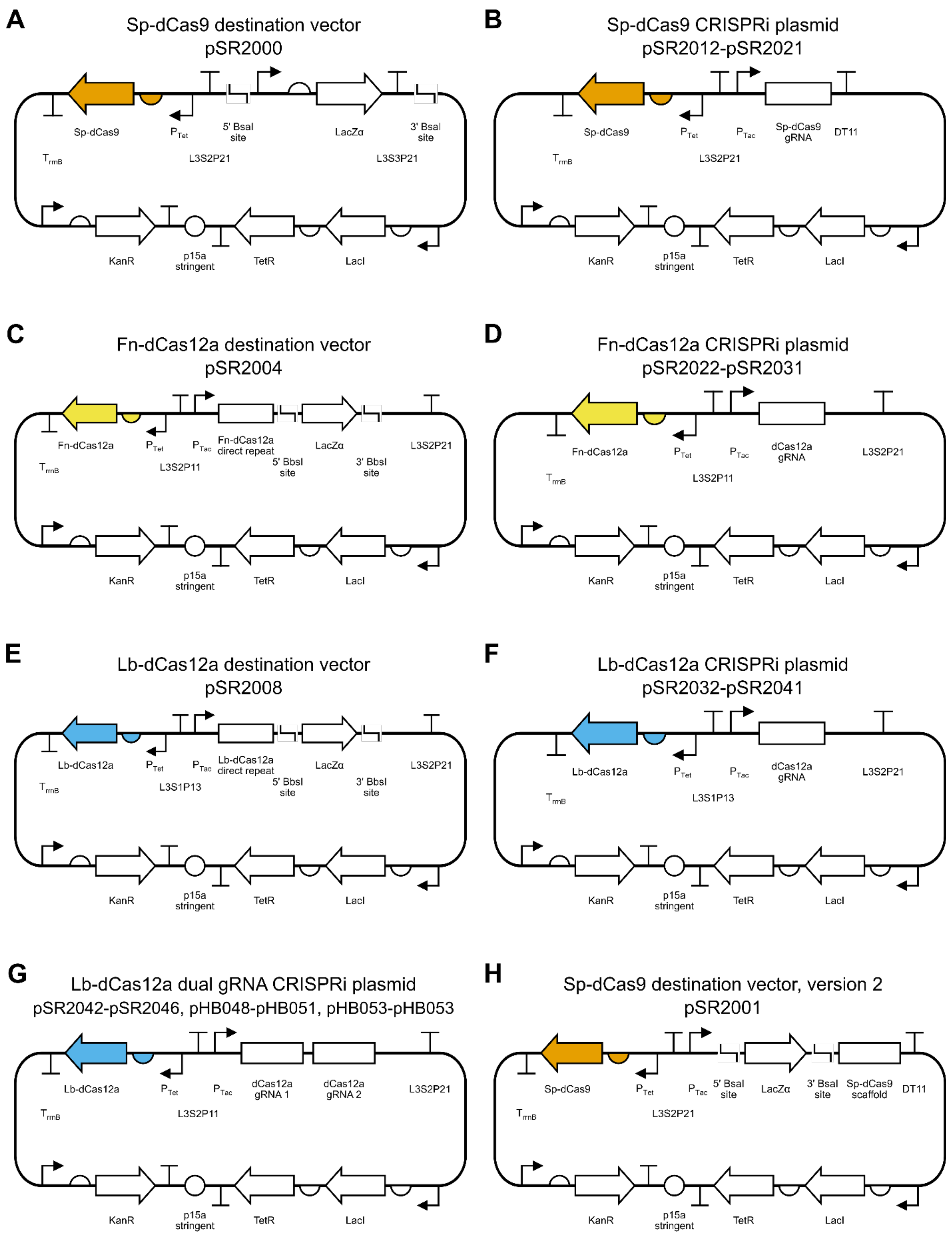

**Figure S32. Plasmid maps of the CRISPRi plasmids.**

Genetic schematics depicting relevant genetic parts are provided for the plasmids used for characterizing the CRISPRi systems. Details for each of these plasmids are provided in **Table S7**. All CRISPRi plasmids contain the dCas protein and gRNA in divergent transcriptional units with a terminator between them. Additionally, they contain a sensor block with TetR and LacI for the $P_{Tet}$ and $P_{Tac}$ inducible promoters, respectively. **(A)** The destination vector for the Sp-dCas9 CRISPRi system (pSR2000) contains the Sp-dCas9 gene and a LacZα cassette surrounded by BsaI recognition sites and 4 nt linker sequences on the low copy number plasmid replicon p15a stringent. **(B)** A representative plasmid used to characterize the Sp-dCas9 system, created by replacing the LacZα cassette with the $P_{Tac}$ promoter, an Sp-dCas9 gRNA, and the DT11 terminator in a Type IIS DNA assembly using BsaI. **(C)** The destination vector for the Fn-dCas12a CRISPRi system (pSR2004) contains the Fn-dCas12a gene and a LacZα cassette surrounded by BbsI recognition sites and linker sequences (downstream of $P_{Tac}$ and the Fn-Cas12a direct repeat sequence) on the low copy number plasmid replicon p15a stringent. **(D)** A representative plasmid used to characterize the Fn-dCas12a system, created by replacing the LacZα cassette with a 20 base pair gRNA sequence for Fn-dCas12a in a Type IIS DNA assembly using BbsI. **(E)** The destination vector for the Lb-dCas12a CRISPRi system (pSR2004) contains the Lb-dCas12a gene and a LacZα cassette surrounded by BbsI recognition sites and linker sequences (downstream of $P_{Tac}$ and the Lb-Cas12a direct repeat sequence) on the low copy number plasmid replicon p15a stringent. **(F)** A representative plasmid used to characterize the Lb-dCas12a system, created by replacing the LacZα cassette with a 20 base pair gRNA sequence for Lb-dCas12a in a Type IIS DNA assembly using BbsI. **(G)** A representative plasmid used to characterize dual gRNA designs for the Lb-dCas12a system, created by replacing the LacZα cassette a dual gRNA array (two 20 base pair spacer sequences separated by a Lb-dCas12a direct repeat sequence) in a Type IIS DNA assembly using BbsI. **(H)** Alternative design of the destination vector for Sp-dCas9 (pSR2001), which has an identical design except the location where the gRNA is inserted was altered to simplify DNA assembly. Here, $P_{Tac}$ is upstream of the LacZα coding sequence, which remains surrounded by BsaI recognition sites and appropriate linker sequences, while the Sp-dCas9 chimeric scaffold sequence and DT11 terminator are downstream of LacZα. This design only requires insertion of the 20 base pair gRNA spacer sequence to target a desired gene.

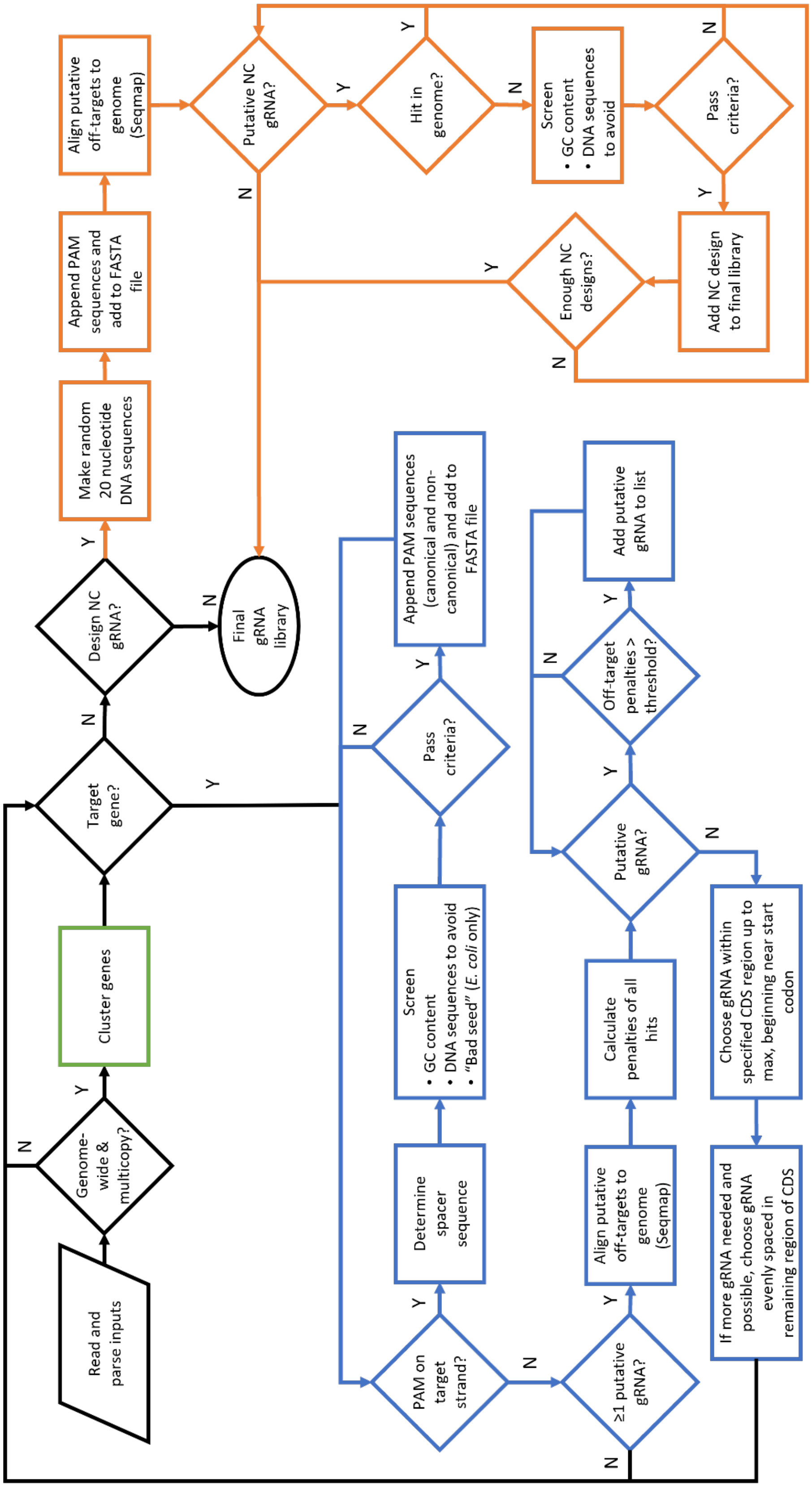

**Figure S33. Flowchart of Python scripts for design of gRNAs.**

A detailed flowchart depicting the Python script set that can be used to design gRNA and gRNA libraries for Sp-dCas9, Fn-dCas12a, or Lb-dCas12a in any organism with an annotated genome, with each colored section indicating a different script in the set. Parameters from a configuration file and other input files (including FASTA files of gene targets and the host organism's genome) are used to customize the design of gRNAs, targeting any number of genes from one to all those in an entire genome (black). If designated, target genes are clustered based on homology from BLAST alignment (green). gRNAs are then designed for each target gene based on input constraints for GC content, DNA sequences that should be avoided, the "bad seed" effect (if applicable),[4] and off-target effects (blue). Different criteria are used to screen preliminary gRNA for each CRISPRi system. Non-targeting negative control (NC) gRNAs are designed, if indicated (orange). A more detailed description this script set, including differences in the design of gRNA for the CRISPRi systems, is provided in **SI Note 4**.

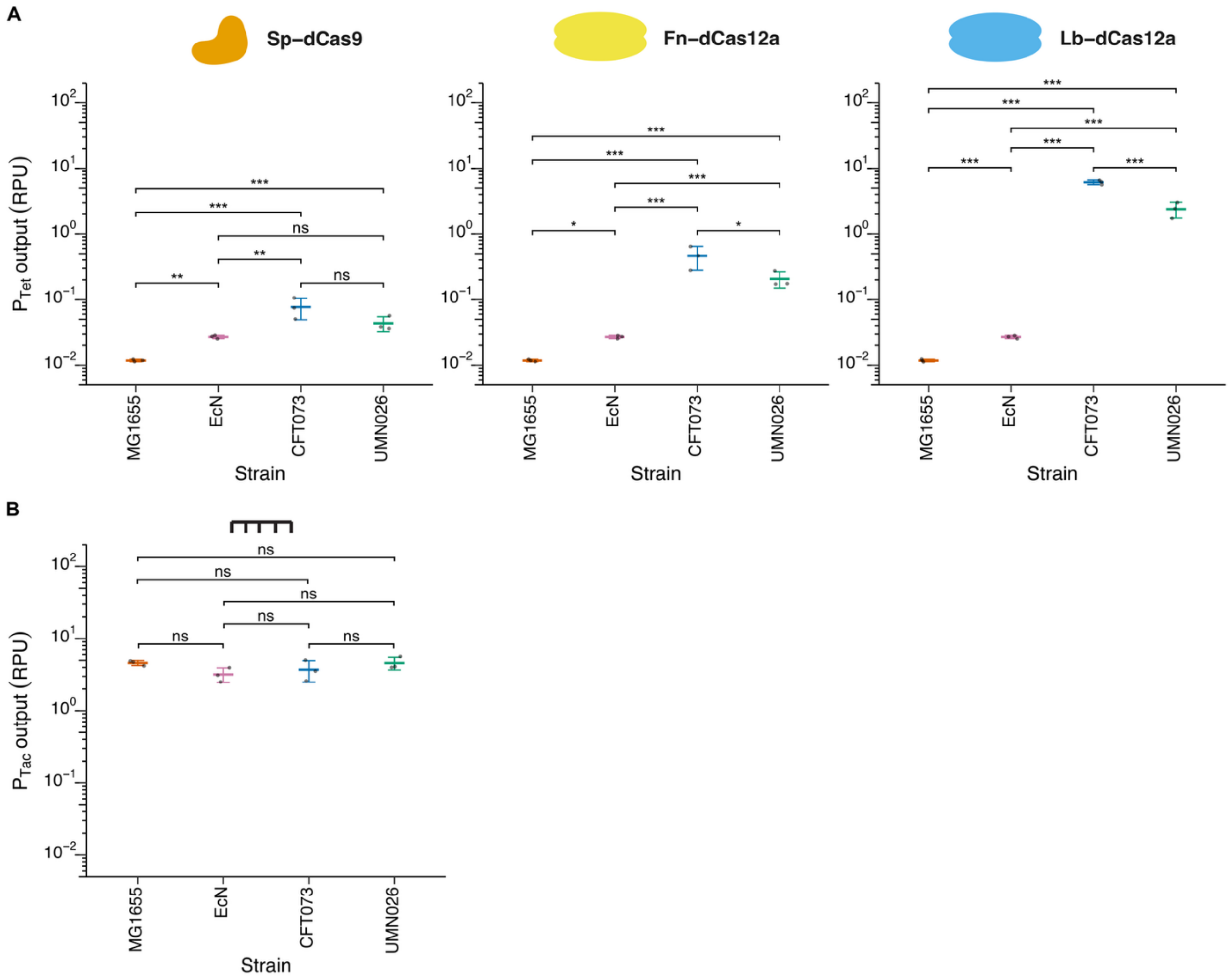

**Figure S34. Comparison of promoter input for CRISPRi component expression**

The outputs from the (**A**) $P_{Tet}$ and (**B**) $P_{Tac}$ promoters chosen to express each dCas protein and gRNA, respectively, are plotted at the induction levels for characterization of the CRISPRi systems in MG1655, EcN, CFT073, and UMN026. Data plotted are those collected from characterization of each inducible promoter using flow cytometry at the inducer concentration chosen for the CRISPRi repression assays, converted to relative promoter units (RPU). For $P_{Tet}$, this was 0.25 ng/mL for MG1655 (all CRISPRi systems), 0.125 ng/mL for EcN (all CRISPRi systems), and 0.25 (Sp-dCas9), 0.5 (Fn-dCas12a), or 2 (Lb-dCas12a) ng/mL aTc for CFT073 and UMN026. For $P_{Tac}$, 1 mM IPTG was used for all strains and CRISPRi systems. Dashes are the mean of three identical experiments (points) performed on three different days with error bars depicting the standard deviation. One-way ANOVA with Tukey post-hoc analysis for the outputs between all combinations of strains for each inducible promoter and CRISPRi systems (for $P_{Tet}$). Results are shown as not significant (ns) or significant (*$p < 0.05$, **$p < 0.01$, and ***$p < 0.001$). Calculated *p*-values are reported in **SI File 1**.

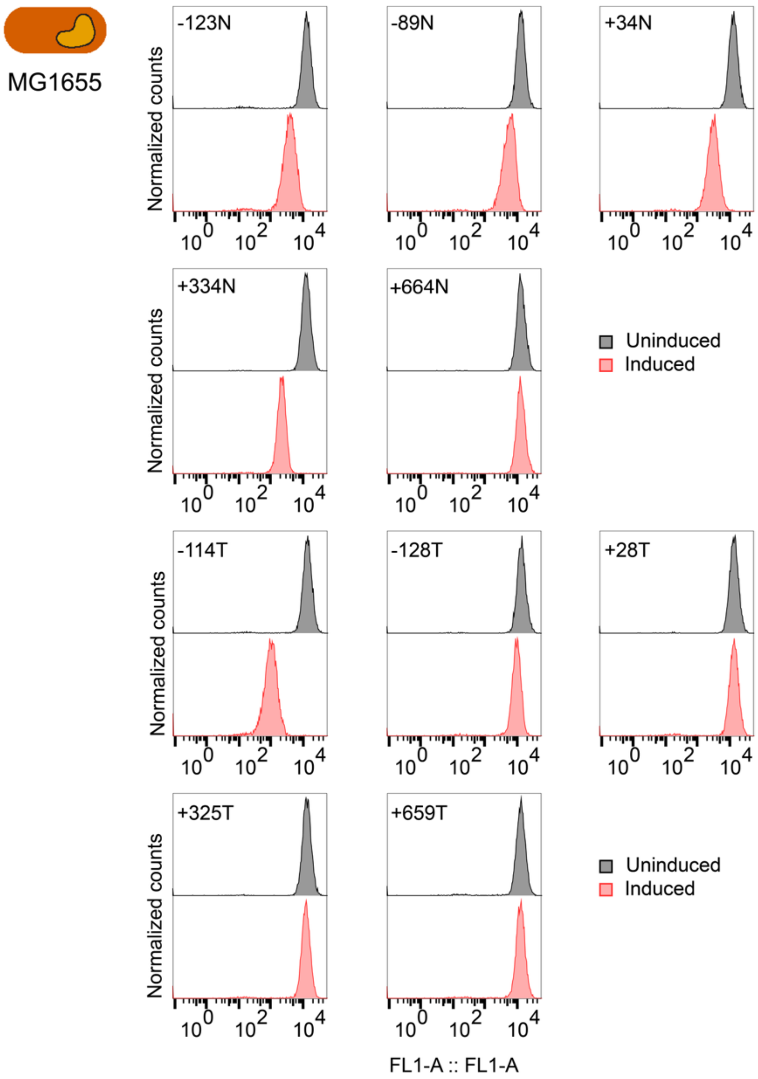

**Figure S35. Histograms of Sp-dCas9 CRISPRi repression in MG1655**

Representative histograms of the normalized cell count for the cell fluorescence used to characterize CRISPRi repression for the ten gRNA designs for Sp-dCas9 in MG1655. Histograms are provided for the same biological sample with the CRISPRi components (Sp-dCas9 and gRNA) uninduced (gray) and induced (red). Fluorescence was measured using the FL1-A channel of the BD Accuri C6 cytometer. Histograms display gated cell events for MG1655. Each histogram includes 4,500 to 10,000 cells.

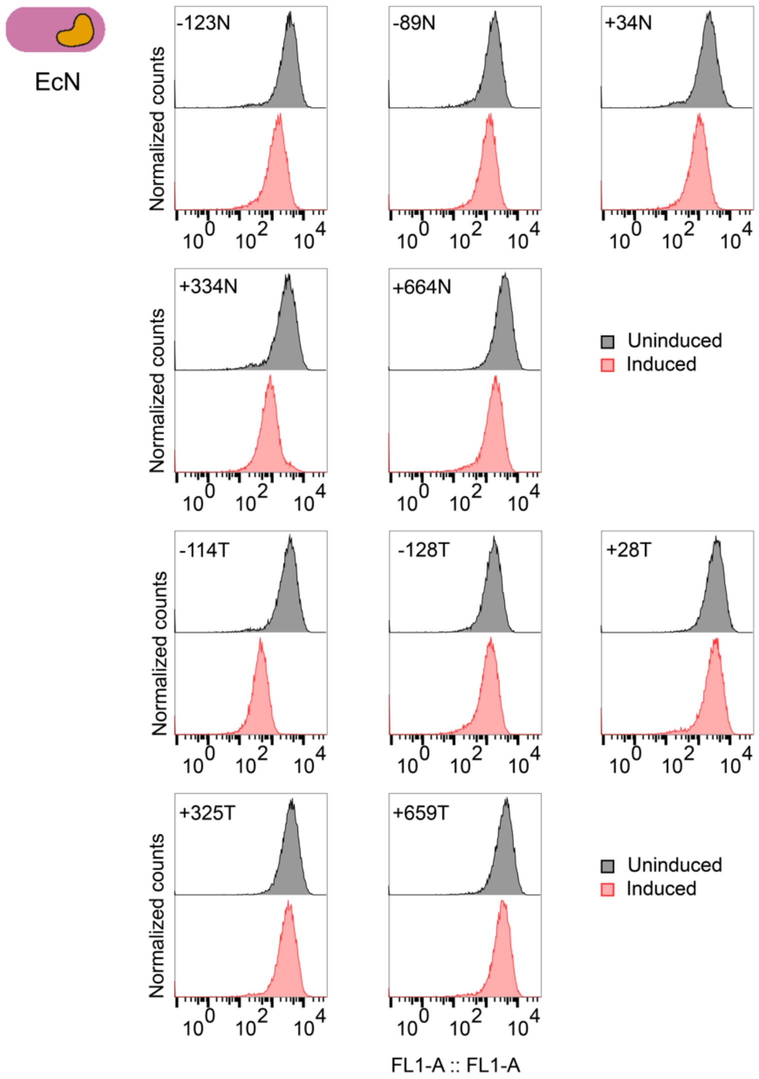

**Figure S36. Histograms of Sp-dCas9 CRISPRi repression in EcN**

Representative histograms of the normalized cell count for the cell fluorescence used to characterize CRISPRi repression for the ten gRNA designs for Sp-dCas9 in EcN. Histograms are provided for the same biological sample with the CRISPRi components (Sp-dCas9 and gRNA) uninduced (gray) and induced (red). Fluorescence was measured using the FL1-A channel of the BD Accuri C6 cytometer. Histograms display gated cell events for EcN. Each histogram includes 4,500 to 10,000 cells.

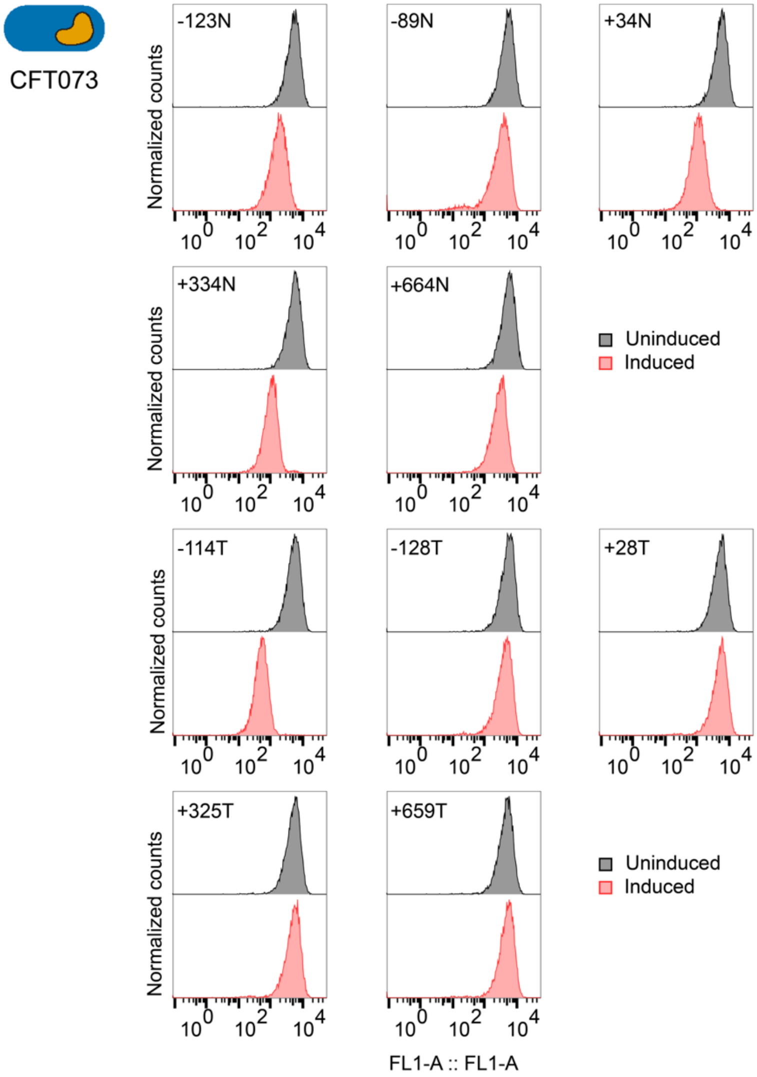

**Figure S37. Histograms of Sp-dCas9 CRISPRi repression in CFT073**

Representative histograms of the normalized cell count for the cell fluorescence used to characterize CRISPRi repression for the ten gRNA designs for Sp-dCas9 in CFT073. Histograms are provided for the same biological sample with the CRISPRi components (Sp-dCas9 and gRNA) uninduced (gray) and induced (red). Fluorescence was measured using the FL1-A channel of the BD Accuri C6 cytometer. Histograms display gated cell events for CFT073. Each histogram includes 4,500 to 10,000 cells.

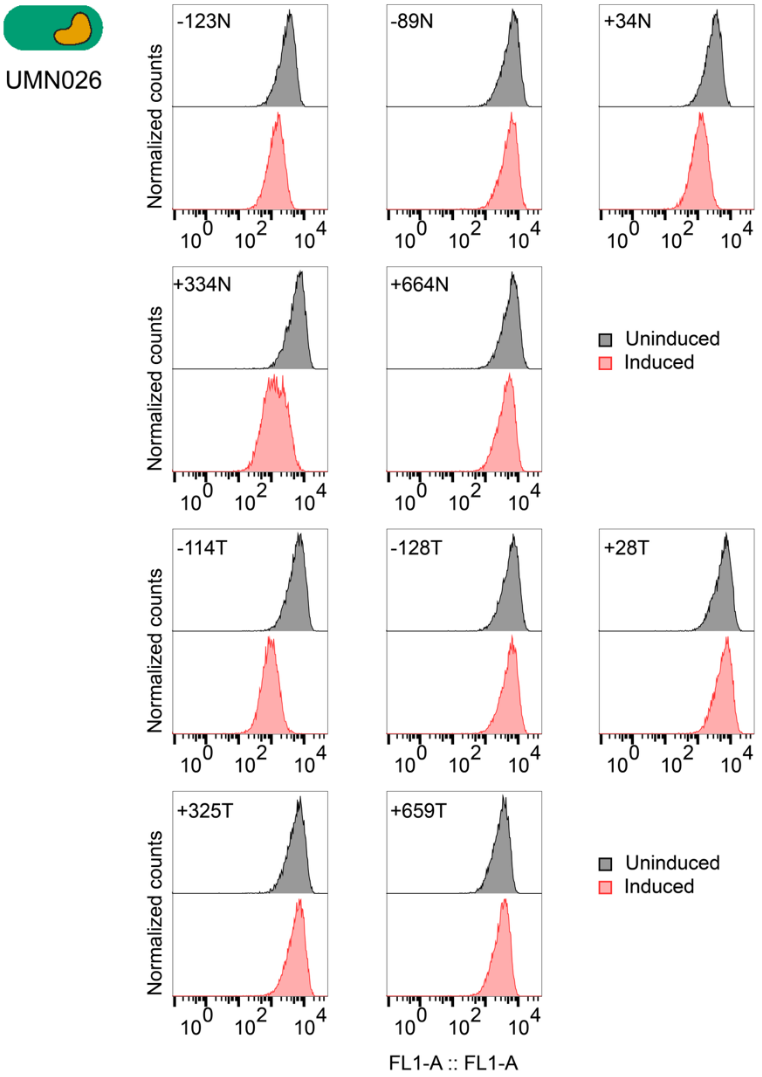

**Figure S38. Histograms of Sp-dCas9 CRISPRi repression in UMN026**

Representative histograms of the normalized cell count for the cell fluorescence used to characterize CRISPRi repression for the ten gRNA designs for Sp-dCas9 in UMN026. Histograms are provided for the same biological sample with the CRISPRi components (Sp-dCas9 and gRNA) uninduced (gray) and induced (red). Fluorescence was measured using the FL1-A channel of the BD Accuri C6 cytometer. Histograms display gated cell events for UMN026. Each histogram includes 4,500 to 10,000 cells.

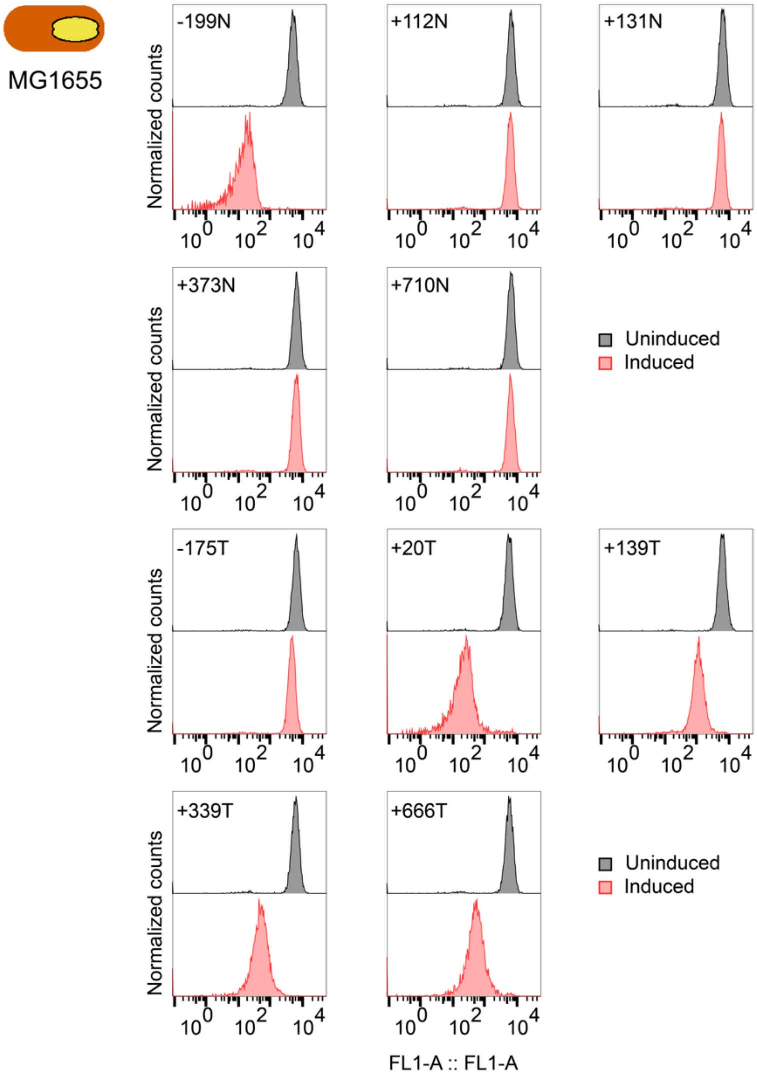

**Figure S39. Histograms of Fn-dCas12a CRISPRi repression in MG1655**

Representative histograms of the normalized cell count for the cell fluorescence used to characterize CRISPRi repression for the ten gRNA designs for Fn-dCas12a in MG1655. Histograms are provided for the same biological sample with the CRISPRi components (Fn-dCas12a and gRNA) uninduced (gray) and induced (red). Fluorescence was measured using the FL1-A channel of the BD Accuri C6 cytometer. Histograms display gated cell events for MG1655. Each histogram includes 4,500 to 10,000 cells.

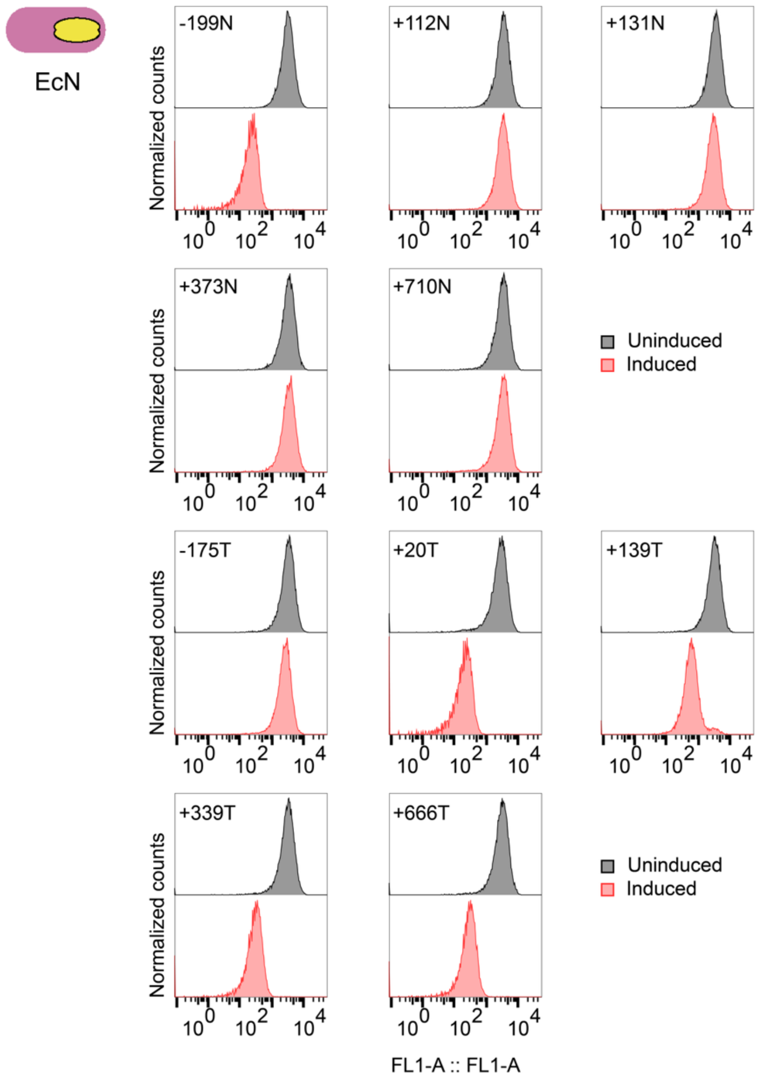

**Figure S40. Histograms of Fn-dCas12a CRISPRi repression in EcN**

Representative histograms of the normalized cell count for the cell fluorescence used to characterize CRISPRi repression for the ten gRNA designs for Fn-dCas12a in EcN. Histograms are provided for the same biological sample with the CRISPRi components (Fn-dCas12a and gRNA) uninduced (gray) and induced (red). Fluorescence was measured using the FL1-A channel of the BD Accuri C6 cytometer. Histograms display gated cell events for EcN. Each histogram includes 4,500 to 10,000 cells.

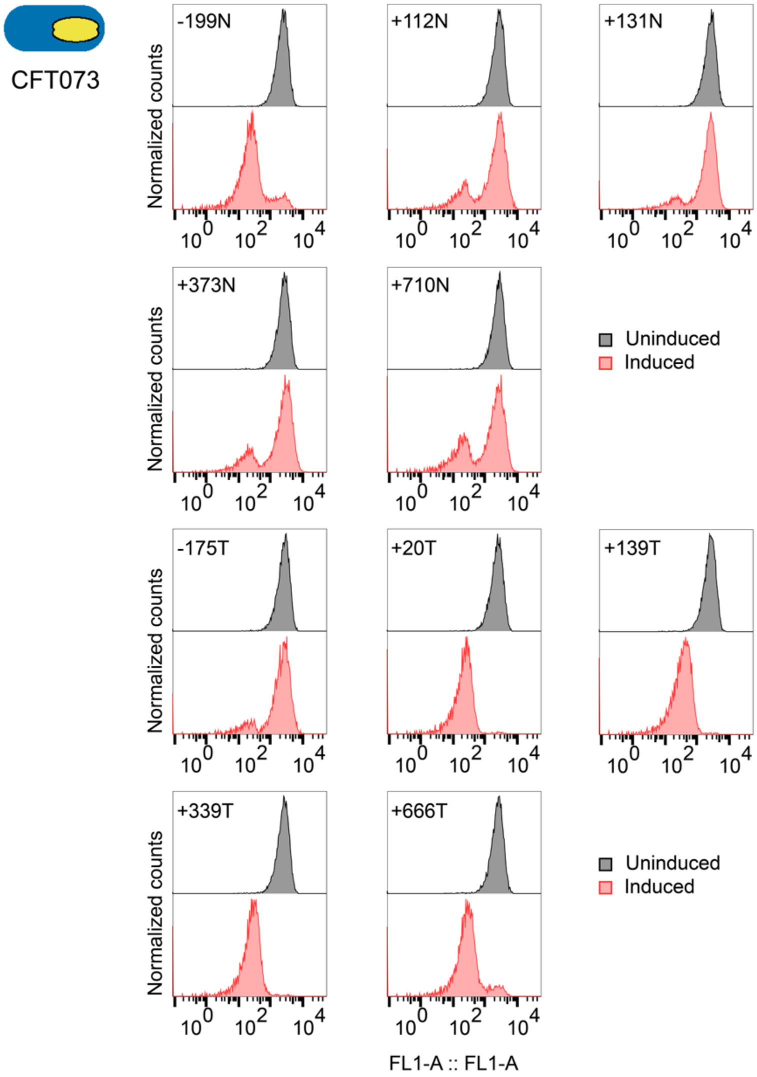

**Figure S41. Histograms of Fn-dCas12a CRISPRi repression in CFT073**

Representative histograms of the normalized cell count for the cell fluorescence used to characterize CRISPRi repression for the ten gRNA designs for Fn-dCas12a in CFT073. Histograms are provided for the same biological sample with the CRISPRi components (Fn-dCas12a and gRNA) uninduced (gray) and induced (red). Fluorescence was measured using the FL1-A channel of the BD Accuri C6 cytometer. Histograms display gated cell events for CFT073. Each histogram includes 4,500 to 10,000 cells.

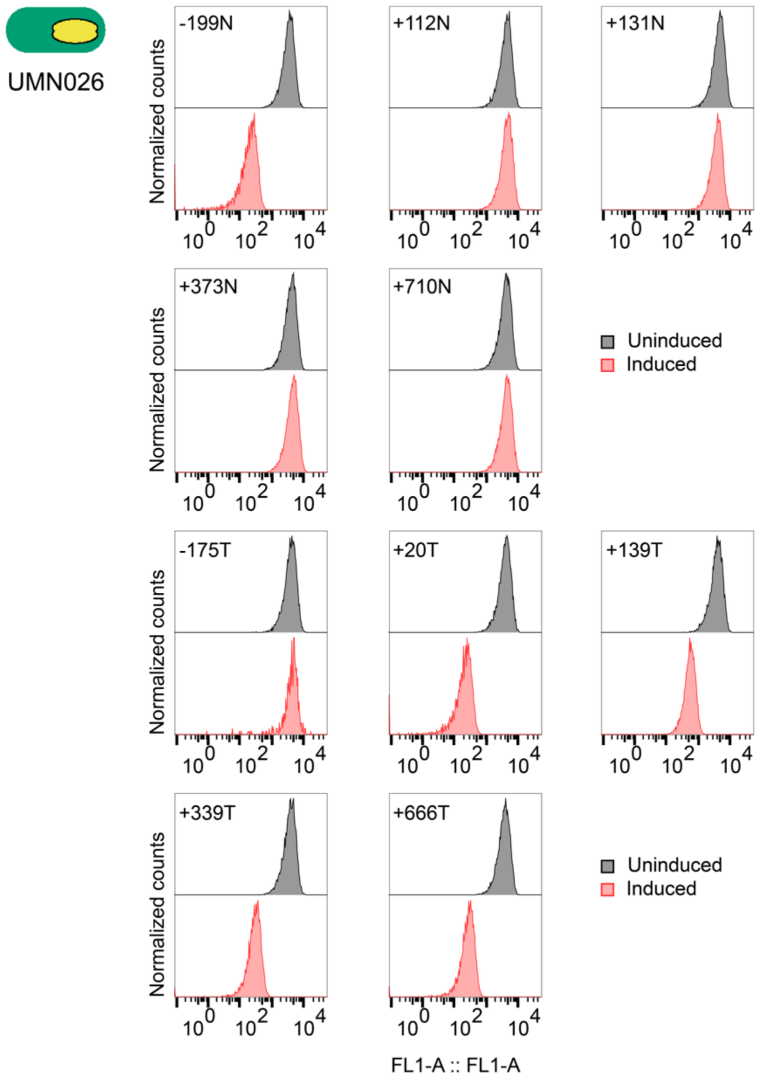

**Figure S42. Histograms of Fn-dCas12a CRISPRi repression in UMN026**

Representative histograms of the normalized cell count for the cell fluorescence used to characterize CRISPRi repression for the ten gRNA designs for Fn-dCas12a in UMN026 Histograms are provided for the same biological sample with the CRISPRi components (Fn-dCas12a and gRNA) uninduced (gray) and induced (red). Fluorescence was measured using the FL1-A channel of the BD Accuri C6 cytometer. Histograms display gated cell events for UMN026. Each histogram includes 4,500 to 10,000 cells.

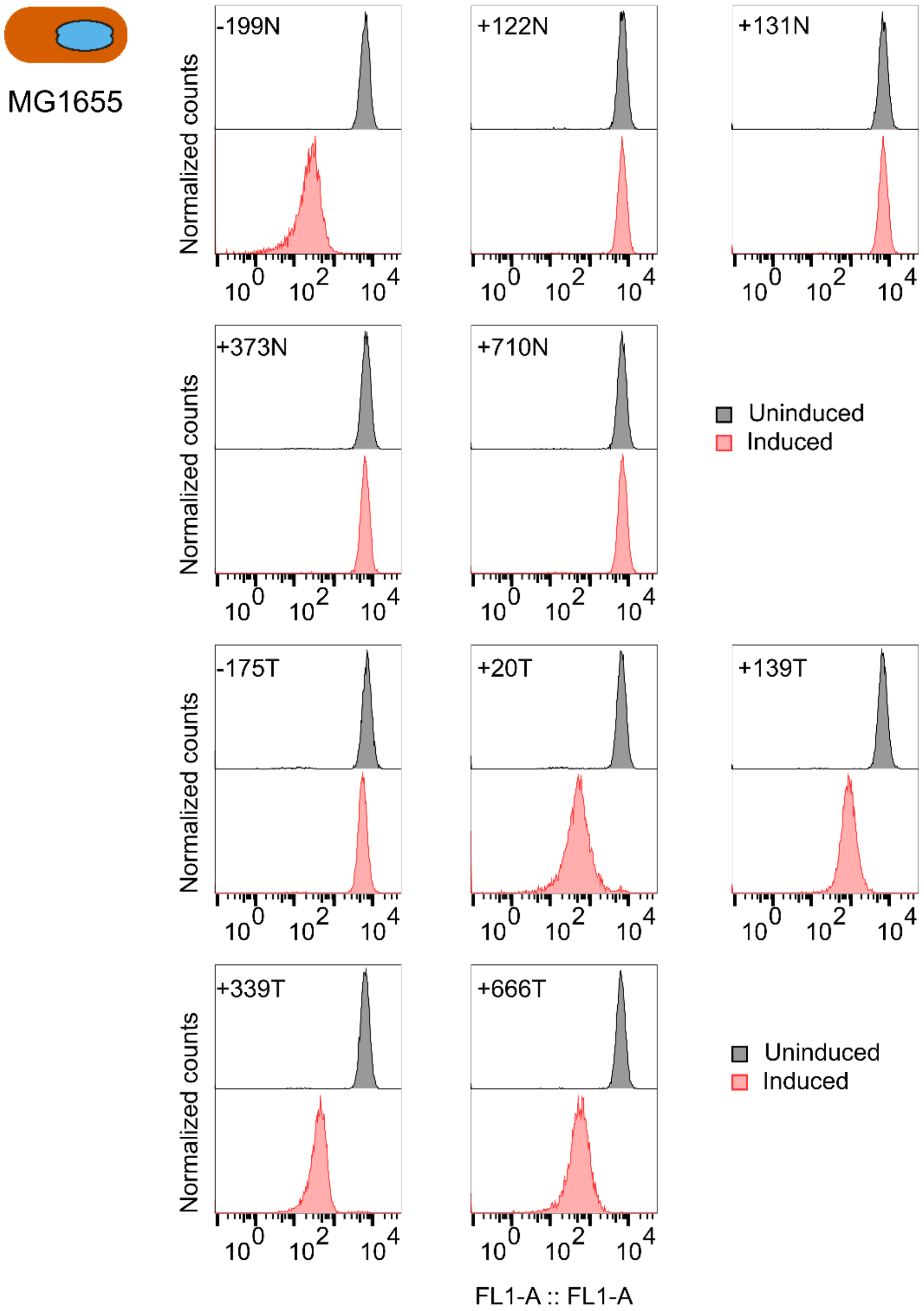

**Figure S43. Histograms of Lb-dCas12a CRISPRi repression in MG1655**

Representative histograms of the normalized cell count for the cell fluorescence used to characterize CRISPRi repression for the ten gRNA designs for Lb-dCas12a in MG1655. Histograms are provided for the same biological sample with the CRISPRi components (Lb-dCas12a and gRNA) uninduced (gray) and induced (red). Fluorescence was measured using the FL1-A channel of the BD Accuri C6 cytometer. Histograms display gated cell events for MG1655. Each histogram includes 4,500 to 10,000 cells.

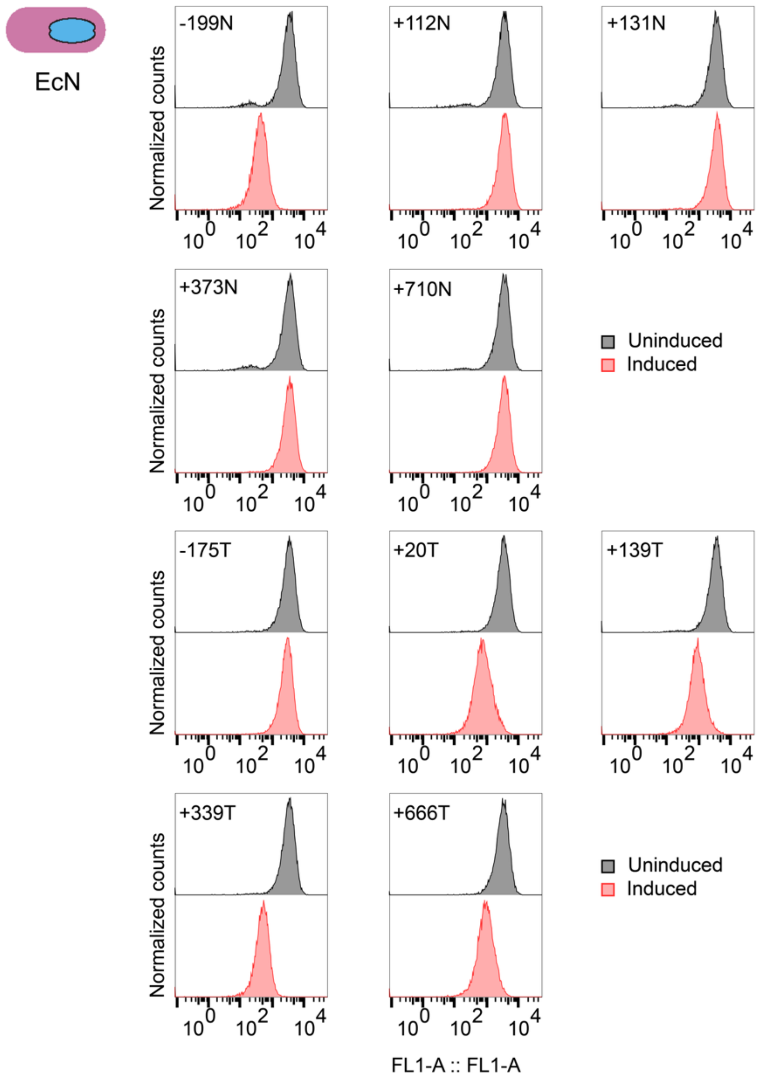

**Figure S44. Histograms of Lb-dCas12a CRISPRi repression in EcN**

Representative histograms of the normalized cell count for the cell fluorescence used to characterize CRISPRi repression for the ten gRNA designs for Lb-dCas12a in EcN. Histograms are provided for the same biological sample with the CRISPRi components (Lb-dCas12a and gRNA) uninduced (gray) and induced (red). Fluorescence was measured using the FL1-A channel of the BD Accuri C6 cytometer. Histograms display gated cell events for EcN. Each histogram includes 4,500 to 10,000 cells.

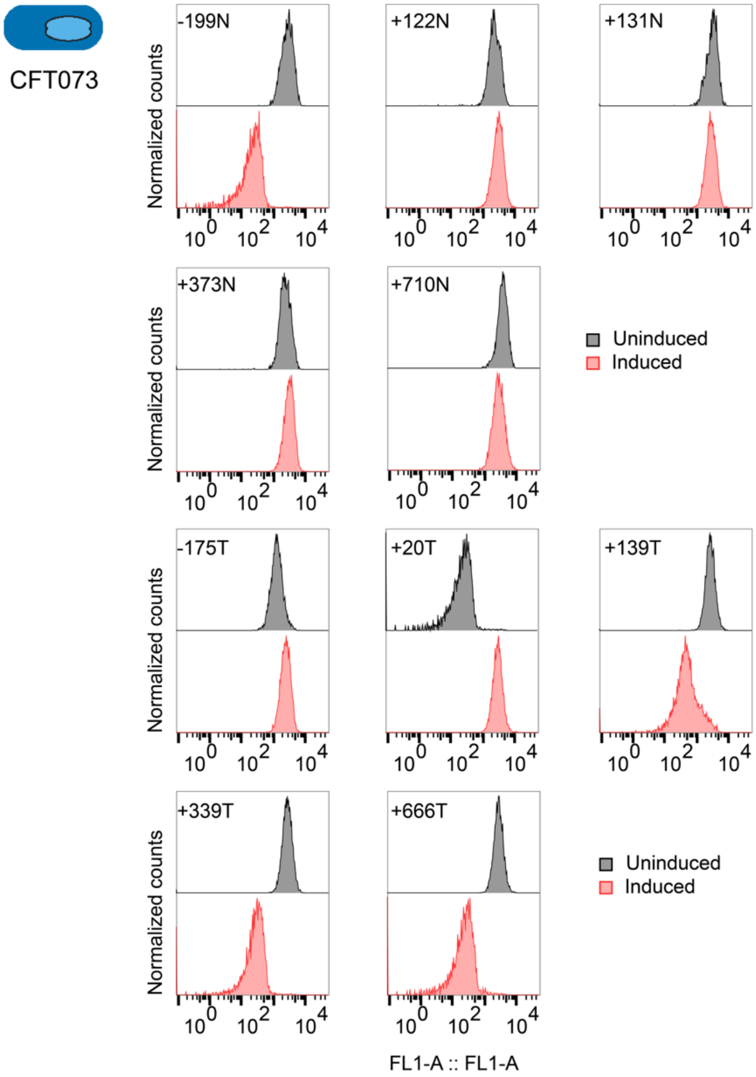

**Figure S45. Histograms of Lb-dCas12a CRISPRi repression in CFT073**

Representative histograms of the normalized cell count for the cell fluorescence used to characterize CRISPRi repression for the ten gRNA designs for Lb-dCas12a in CFT073. Histograms are provided for the same biological sample with the CRISPRi components (Lb-dCas12a and gRNA) uninduced (gray) and induced (red). Fluorescence was measured using the FL1-A channel of the BD Accuri C6 cytometer. Histograms display gated cell events for CFT073. Each histogram includes 4,500 to 10,000 cells.

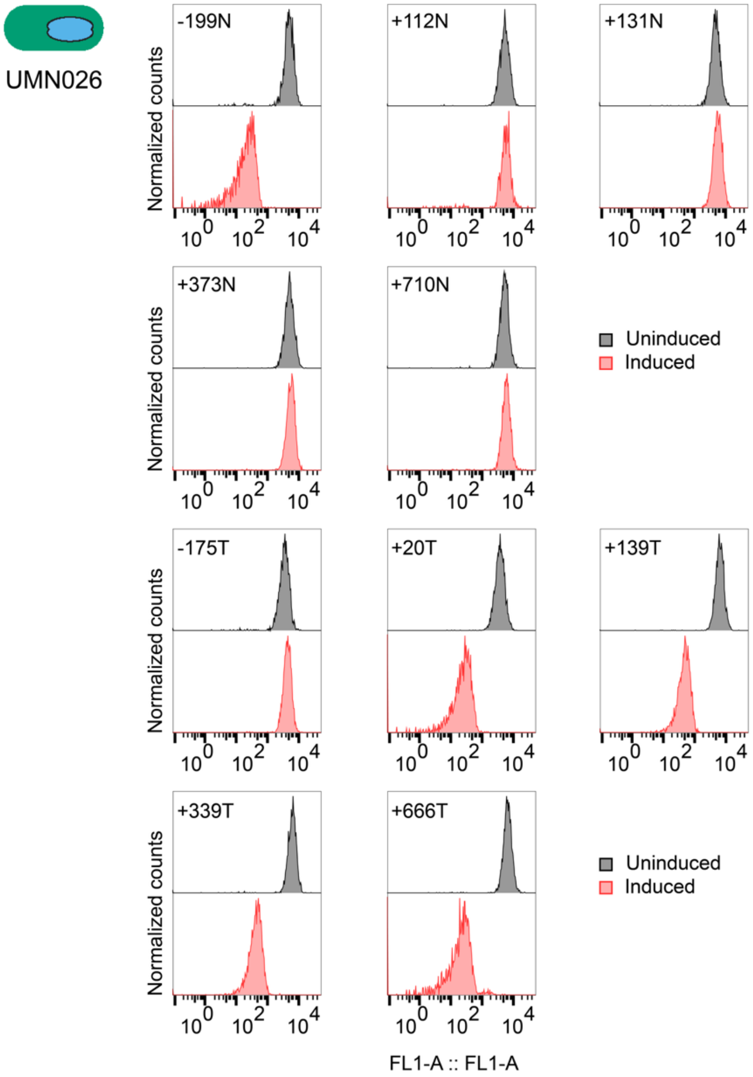

**Figure S46. Histograms of Lb-dCas12a CRISPRi repression in UMN026**

Representative histograms of the normalized cell count for the cell fluorescence used to characterize CRISPRi repression for the ten gRNA designs for Lb-dCas12a in UMN026. Histograms are provided for the same biological sample with the CRISPRi components (Lb-dCas12a and gRNA) uninduced (gray) and induced (red). Fluorescence was measured using the FL1-A channel of the BD Accuri C6 cytometer. Histograms display gated cell events for UMN026. Each histogram includes 4,500 to 10,000 cells.

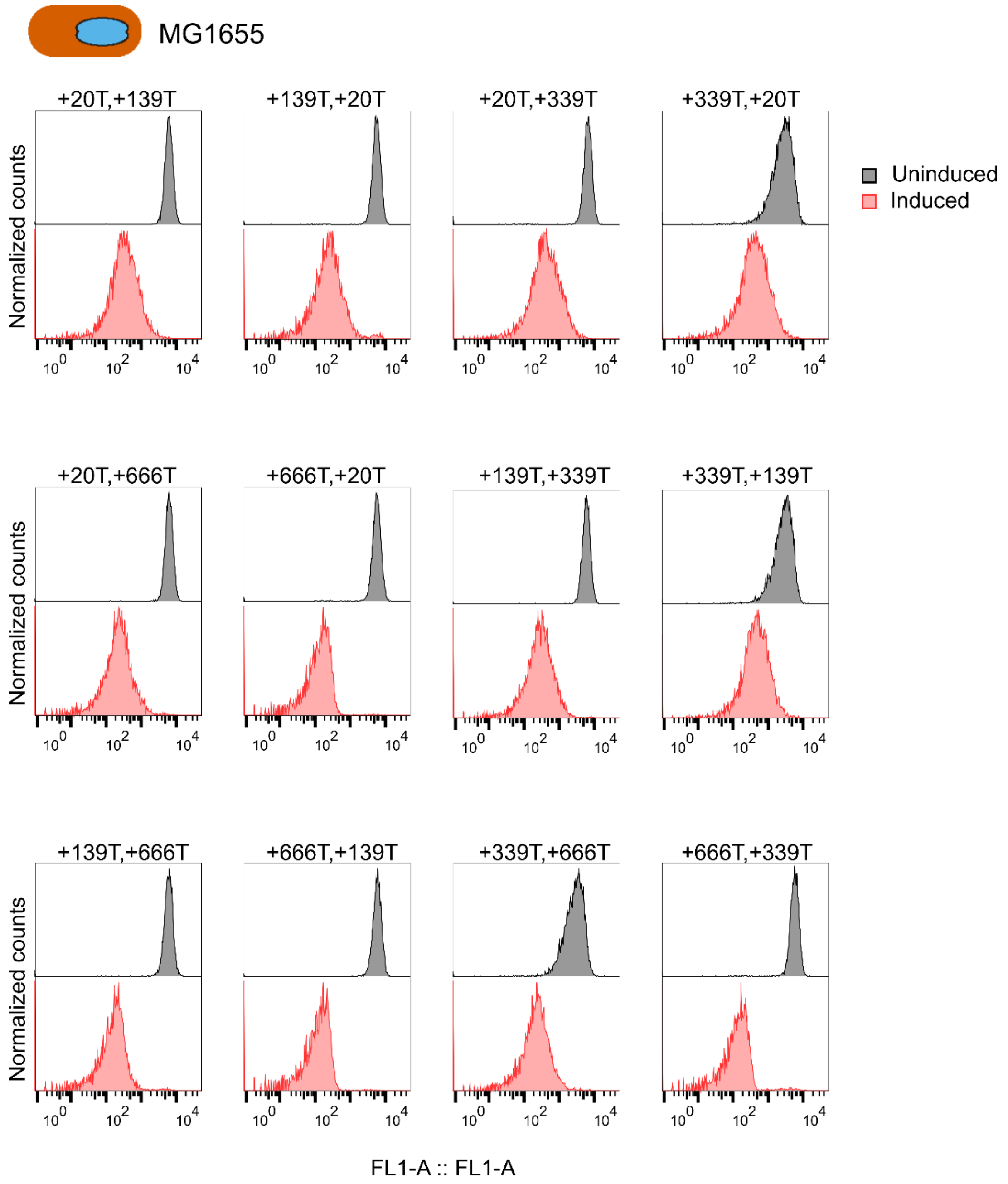

**Figure S47. Histograms of dual gRNA Lb-dCas12a CRISPRi repression in MG1655**

Representative histograms of the normalized cell count for the cell fluorescence used to characterize CRISPRi repression for the twelve dual gRNA designs for Lb-dCas12a in MG1655. Histograms are provided for the same biological sample with the CRISPRi components (Lb-dCas12a and dual gRNA array) uninduced (gray) and induced (red). Fluorescence was measured using the FL1-A channel of the BD Accuri C6 cytometer. Histograms display gated cell events for MG1655. Each histogram includes 4,500 to 10,000 cells.

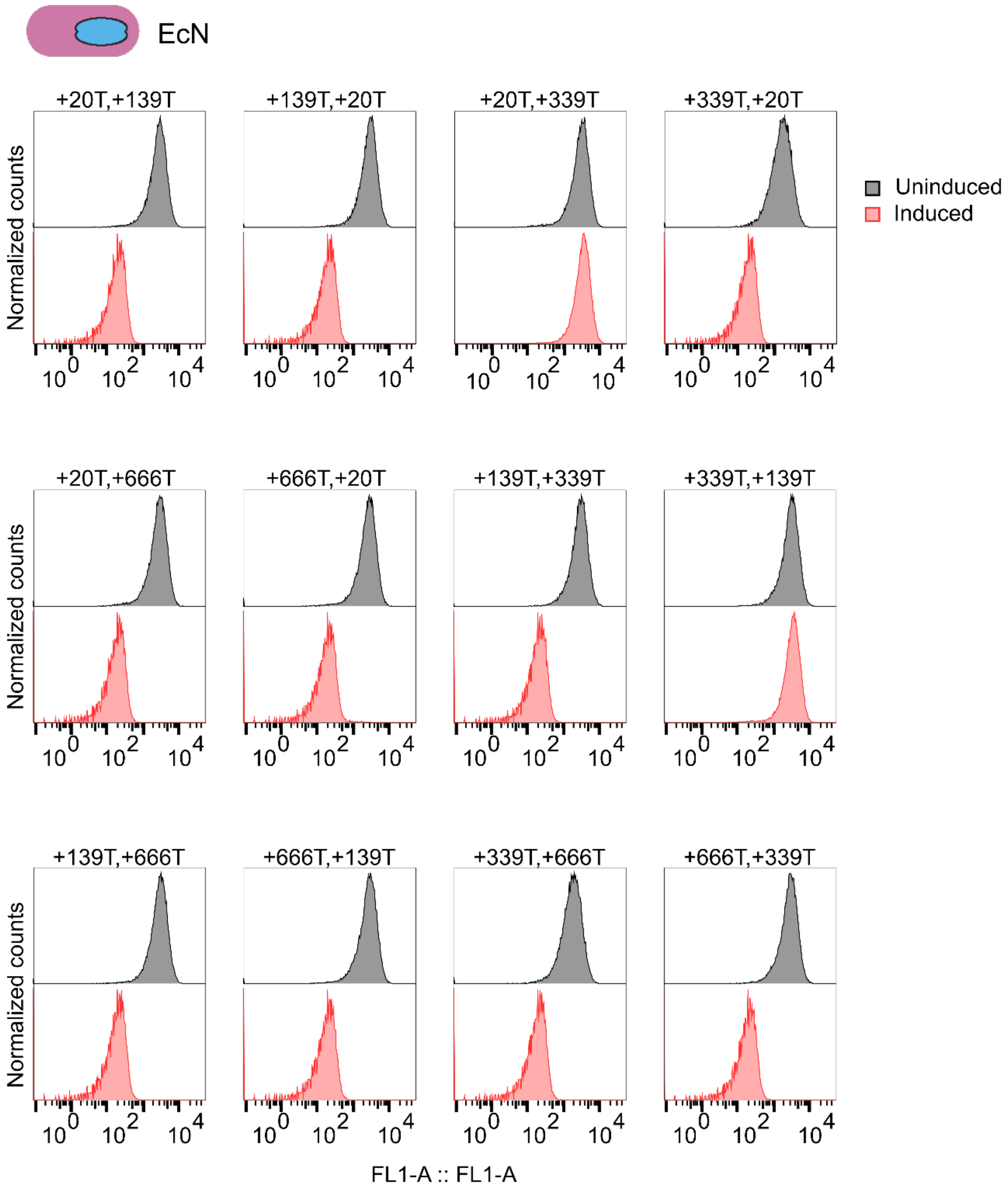

**Figure S48. Histograms of dual gRNA Lb-dCas12a CRISPRi repression in EcN**

Representative histograms of the normalized cell count for the cell fluorescence used to characterize CRISPRi repression for the twelve dual gRNA designs for Lb-dCas12a in EcN. Histograms are provided for the same biological sample with the CRISPRi components (Lb-dCas12a and dual gRNA array) uninduced (gray) and induced (red). Fluorescence was measured using the FL1-A channel of the BD Accuri C6 cytometer. Histograms display gated cell events for EcN. Each histogram includes 4,500 to 10,000 cells.

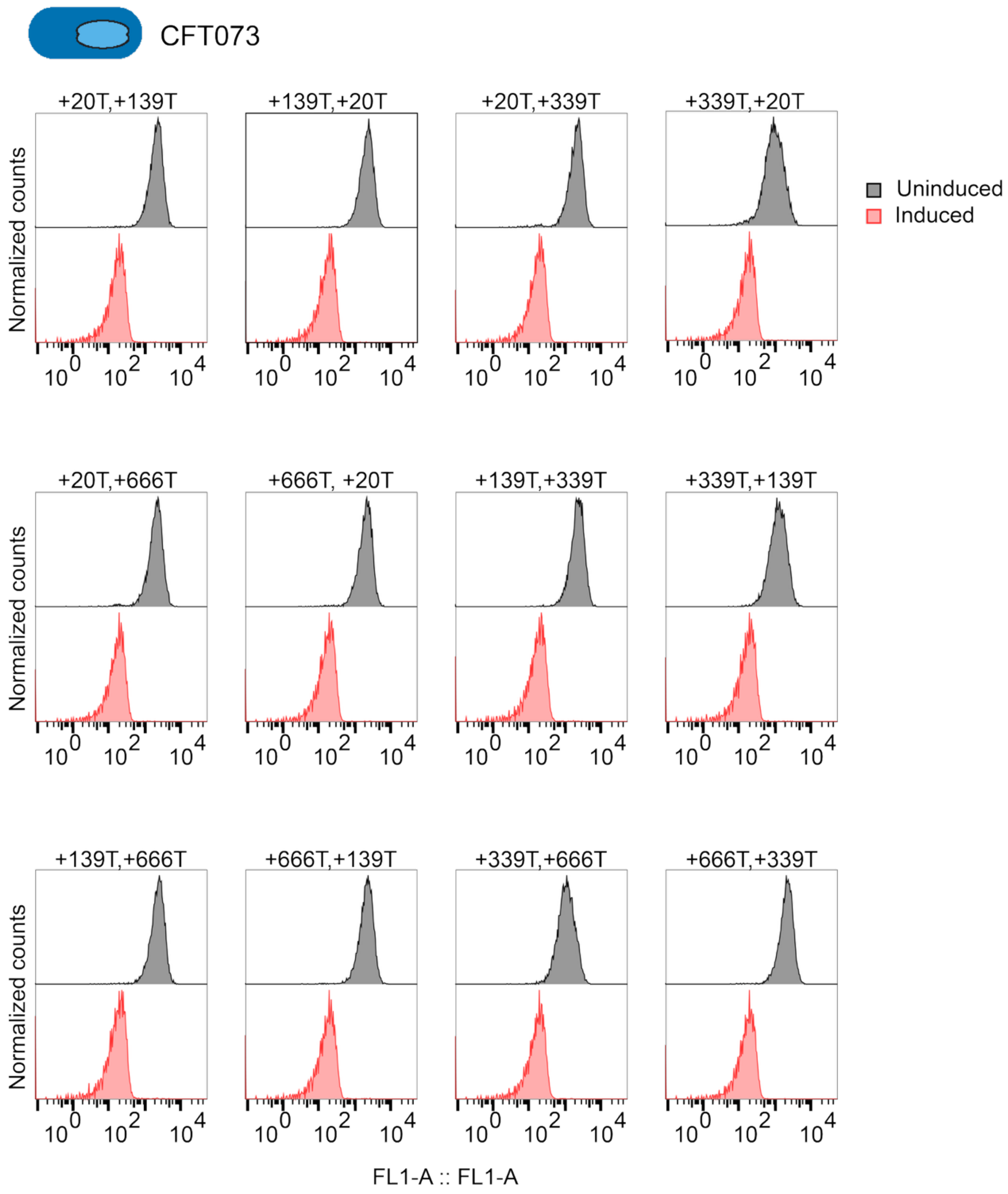

**Figure S49. Histograms of dual gRNA Lb-dCas12a CRISPRi repression in CFT073**

Representative histograms of the normalized cell count for the cell fluorescence used to characterize CRISPRi repression for the twelve dual gRNA designs for Lb-dCas12a in CFT073. Histograms are provided for the same biological sample with the CRISPRi components (Lb-dCas12a and dual gRNA array) uninduced (gray) and induced (red). Fluorescence was measured using the FL1-A channel of the BD Accuri C6 cytometer. Histograms display gated cell events for CFT073. Each histogram includes 4,500 to 10,000 cells.

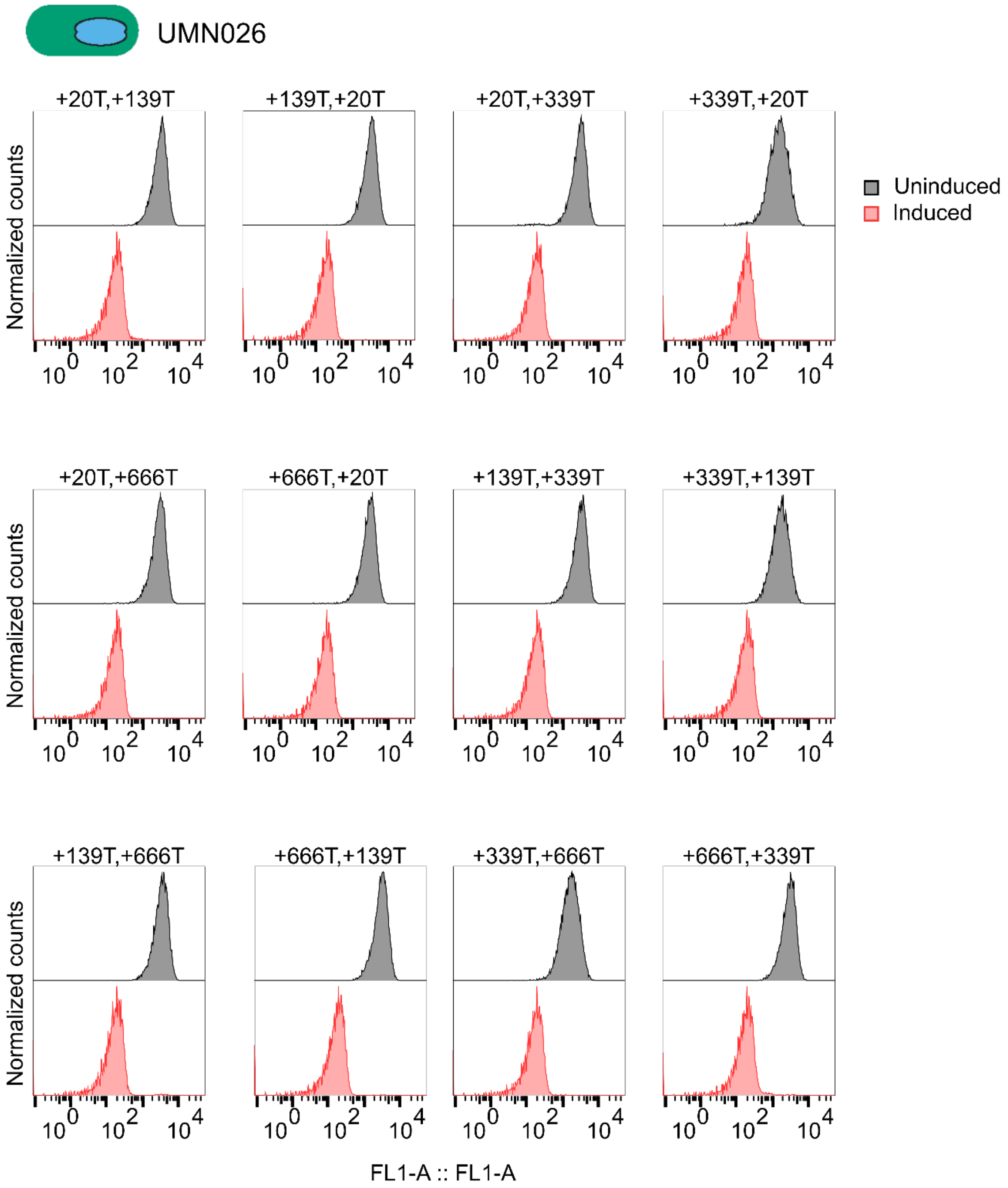

**Figure S50. Histograms of dual gRNA Lb-dCas12a CRISPRi repression in UMN026**

Representative histograms of the normalized cell count for the cell fluorescence used to characterize CRISPRi repression for the twelve dual gRNA designs for Lb-dCas12a in UMN026. Histograms are provided for the same biological sample with the CRISPRi components (Lb-dCas12a and dual gRNA array) uninduced (gray) and induced (red). Fluorescence was measured using the FL1-A channel of the BD Accuri C6 cytometer. Histograms display gated cell events for UMN026. Each histogram includes 4,500 to 10,000 cells.

### Supplementary Tables

#### Table S1. *Escherichia coli* strains used for phylogenetic tree analysis

| Strain | RefSeq Accession Number | Phylogroup[a] | Ecological niche[b] |
|---|---|---|---|
| 11128 | GCF_000010765.1 | B1 | EHEC |
| 11368 | GCF_000091005.1 | B1 | EHEC |
| 2011C-3493 | GCF_000299455.1 | B1 | EAHEC |
| 42 | GCF_008042015.2 | C | EAEC |
| 55989 | GCF_003028695.1 | B2 | EAEC |
| 83972 | GCF_000148365.1 | B2 | ABU |
| APEC O1 | GCF_030347155.1 | B2 | APEC |
| APEC O78 | GCF_000332755.1 | C | APEC |
| BL21 | GCF_019754155.1 | A | Laboratory |
| BW25113 | GCF_004355105.2 | A | Laboratory |
| CB9615 | GCF_000025165.1 | E | EPEC |
| CE10 | GCF_000227625.1 | F | NMEC |
| CFT073 | GCF_000007445.1 | B2 | UPEC |
| DH10B | GCF_000019425.1 | A | Laboratory |
| E2348/69 | GCF_900149915.1 | B2 | EPEC |
| E24377A | GCF_000017745.1 | B1 | ETEC |
| E42 | GCA_013694085.1 | D | EAEC |
| EC958 | GCF_000285655.3 | B2 | UPEC |
| EcN | GCF_021559835.1 | B2 | Commensal |
| ECOR 48 | GCA_002190935.1 | E | UPEC |
| ECOR 70 | GCA_002190075.1 | C | Commensal |
| *Escherichia fergusonii* | GCF_020097475.1 | N/A | Pathogen |
| HS | GCF_000017765.1 | A | Commensal |
| IAI1 | GCF_000026265.1 | B1 | Commensal |
| IAI39 | GCF_000026345.1 | F | UPEC |
| IHE3034 | GCF_000025745.1 | B2 | NMEC |
| JJ1886 | GCF_000493755.1 | B2 | UPEC |
| LF82 | GCF_021398935.1 | B2 | AIEC |
| MG1655 | GCF_000005845.2 | A | Laboratory |
| O104:H4 | GCF_022869985.1 | B1 | EAEC |
| O157:H7 | GCF_000008865.2 | E | EHEC |
| PCN033 | GCF_000219515.2 | D | ExPEC |
| PMV-1 | GCF_000493595.2 | B2 | ExPEC |
| RM12579 | GCF_000245515.1 | E | EPEC |
| SE15 | GCF_000010485.1 | B2 | Commensal |
| SMS-3-5 | GCF_000019645.1 | F | Environmental |
| UM146 | GCF_000148605.1 | B2 | AIEC |
| UMN026 | GCF_000026325.1 | D | UPEC |
| UMNF18 | GCF_000220005.1 | A | ETEC |
| UMNK88 | GCF_000212715.2 | A | ETEC |
| UTI89 | GCF_015644765.1 | B2 | UPEC |

[a] Phylogroups were determined using ClermonTyping[1]

[b] Acronyms for specific ecological niches: ABU = asymptomatic bacteriuria, AIEC = adherent-invasive *E. coli*, APEC = avian-pathogenic *E. coli*, EAEC = enteroaggregative *E. coli*, EAHEC = enteroaggregative and hemorrhagic *E. coli*, EHEC = enterohemorrhagic *E. coli*, EIEC = enteroinvasive *E. coli*, EPEC = enteropathogenic *E. coli*, ETEC = enterotoxigenic *E. coli*, ExPEC = extraintestinal pathogenic *E. coli*, NMEC = neonatal meningitis *E. coli*, and UPEC, uropathogenic *E. coli*

**Table S2. DNA sequences of genetic parts**

| Part Name | Type of Part | DNA Sequence | Reference |
|---|---|---|---|
| $P_{Tet}$ | Promoter | TCCACCGTTGGCTTTTTTCCCTATCAGTGATAGAGATTGA CATCCCTATCAGTGATAGAGATAATGAGCAC | [5] |
| $P_{Tac}$ | Promoter | AACGATCGTTGGCTGTGTTGACAATTAATCATCGGCTCGT ATAATGTGTGGAATTGTGAGCGCTCACAATT | [6] |
| J23101 | Promoter | TTTACAGCTAGCTCAGTCCTAGGTATTATGCTAGC | [7] |
| P1 | Promoter | TTGACAGGTCCTAAAGGATAGTCTATAATGCTAGC | [7] |
| $P_{PhlF}$ | Promoter | CGACGTACGGTGGAAtctgattcgttaccaattgacATGATACGAAA CGTACCGTATCGTTAAGGT | [5] |
| Bba_B0064 | Synthetic RBS | AAAGAGGGGAAA | [8] |
| sfGFP-2.5k | Synthetic RBS | CTCTCGAAGGCCCCTGACACA | This study, created using Salis lab's RBS calculator[9–12] |
| Bba_B0034[a] | Synthetic RBS | AAAGAGGAGAAA | [13] |
| dCas12a RBS | Synthetic RBS | AGGTGCCCTCAAAGGAGCTGGATTA | This study, created using Salis lab's RBS calculator[9–12] |
| L3S2P21 | Terminator | CTCGGTACCAAATTCCAGAAAAGAGGCCTCCCGAAAGGG GGGCCTTTTTTCGTTTTGGTCC | [14] |
| L3S3P21 | Terminator | GGGAGACCAGAAACAAAAAAAGGCCCCCCGTTAGGGAG GCCTTCAATAATTGG | [14] |
| L3S2P11 | Terminator | CTCGGTACCAAATTCCAGAAAAGAGACGCTTTCGAGCGT CTTTTTTCGTTTTGGTC | [14] |
| L3S2P44 | Terminator | CTCGGTACCAAACCAATTATTGAAGACGCTGAAAAGCGT CTTTTTTTGTTTCGGTCC | [14] |
| L3S1P13 | Terminator | GACGAACAATAAGGCCTCCCTAACGGGGGGCCTTTTTTA TTGATAACAAAA | [14] |
| $T_{rrnB}$ | Terminator | CAAATAAAACGAAAGGCTCAGTCGAAAGACTGGGCCTTT CGTTTTATCTGTTGTTTGTCGGTGAACGCTCTC | [15] |
| DT11 | Double terminator | AACGCATGAGAAAGCCCCCGGAAGATCACCTTCCGGGG GCTTTTTTATTGCGCTCCTTGGCCCTCCATCCTTAGATAG CTCGGTACCAAATTCCAGAAAAGAGGCCTCCCGAAAGGG GGGCCTTTTTTCGTTTTGGTCC | [14] |
| Spacer DNA[b] | Spacer DNA | AAGCTTGGCTGTTTTGGCGGATGAGAGAAGATTTTCAGC CTGATACAGATTAAATCAGAACGCAGAAGCGGTCTGATAA AACAGAATT | [16] |
| Sp-dCas9 scaffold | Chimeric sgRNA scaffold | GTTTTAGAGCTAGAAATAGCAAGTTAAAATAAGGCTAGTC CGTTATCAACTTGAAAAAGTGG | [17] |
| Fn-dCas12a direct repeat | Direct repeat | GTCTAAGAACTTTAAATAATTTCTACTGTTGTAGAT | [18] |

| | | | |
|---|---|---|---|
| Lb-dCas12a direct repeat | Direct repeat | GTTTCAAAGATTAAATAATTTCTACTAAGTGTAGAT | [18] |
| RiboJ | Ribozyme (genetic insulator) | AGCTGTCACCGGATGTGCTTTCCGGTCTGATGAGTCCGT GAGGACGAAACAGCCTCTACAAATAATTTTGTTTAA | [3] |
| *sfgfp* | coding sequence | ATGAGCAAAGGAGAAGAACTTTTCACTGGAGTTGTCCCA ATTCTTGTTGAATTAGATGGTGATGTTAATGGGCACAAAT TTTCTGTCCGTGGAGAGGGTGAAGGTGATGCTACAAACG GAAAACTCACCCTTAAATTTATTTGCACTACTGGAAAACTA CCTGTTCCGTGGCCAACACTTGTCACTACTCTGACCTATG GTGTTCAATGCTTTTCCCGTTATCCGGATCACATGAAACG TCATGACTTTTTCAAGAGTGCCATGCCTGAAGGTTATGTA CAGGAACGCACTATATCTTTCAAAGATGACGGGACCTAC AAGACGCGTGCTGAAGTCAAGTTTGAAGGTGATACCCTT GTTAATCGTATCGAGTTAAAGGGTATTGATTTTAAAGAAG ATGGAAACATTCTTGGACACAAACTCGAGTACAACTTTAA CTCACACAATGTATACATCACGGCAGACAAACAAAAGAAT GGAATCAAAGCTAACTTCAAAATTCGCCACAACGTTGAAG ATGGTTCCGTTCAACTAGCAGACCATTATCAACAAAATAC TCCAATTGGCGATGGCCCTGTCCTTTTACCAGACAACCAT TACCTGTCGACACAATCTGTCCTTTCGAAAGATCCTAACG AAAAGCGTGACCACATGGTCCTTCTTGAGTTTGTAACTGC TGCTGGGATTACACATGGCATGGATGAGCTCTACAAA | [19] |
| *eyfp* | coding sequence | ATGGTGAGCAAGGGCGAGGAGCTGTTCACCGGGGTGGT GCCCATCCTGGTCGAGCTGGACGGCGACGTAAACGGCC ACAAGTTCAGCGTGTCCGGCGAGGGCGAGGGCGATGCC ACCTACGGCAAGCTGACCCTGAAGTTCATCTGCACCACA GGCAAGCTGCCCGTGCCCTGGCCCACCCTCGTGACCAC CTTCGGCTACGGCCTGCAATGCTTCGCCCGCTACCCCGA CCACATGAAGCTGCACGACTTCTTCAAGTCCGCCATGCC CGAAGGCTACGTCCAGGAGCGCACCATCTTCTTCAAGGA CGACGGCAACTACAAGACCCGCGCCGAGGTGAAGTTCG AGGGCGACACCCTGGTGAACCGCATCGAGCTGAAGGGC ATCGACTTCAAGGAGGACGGCAACATCCTGGGGCACAAG CTGGAGTACAACTACAACAGCCACAACGTCTATATCATGG CCGACAAGCAGAAGAACGGCATCAAGGTGAACTTCAAGA TCCGCCACAACATCGAGGACGGCAGCGTGCAGCTCGCC GACCACTACCAGCAGAACACCCCAATCGGCGACGGCCC CGTGCTGCTGCCCGACAACCACTACCTTAGCTACCAGTC CGCCCTGAGCAAAGACCCCAACGAGAAGCGCGATCACAT GGTCCTGCTGGAGTTCGTGACCGCCGCCGGGATCACTC TCGGCATGGACGAGCTGTACAAGTAA | [20] |
| *tetR* | coding sequence | ATGTCCAGATTAGATAAAAGTAAAGTGATTAACAGCGCAT TAGAGCTGCTTAATGAGGTCGGAATCGAAGGTTTAACAA CCCGTAAACTCGCCCAGAAGCTAGGTGTAGAGCAGCCTA CATTGTATTGGCATGTAAAAAATAAGCGGGCTTTGCTCGA CGCCTTAGCCATTGAGATGTTAGATAGGCACCATACTCAC TTTTGCCCTTTAGAAGGGGAAAGCTGGCAAGATTTTTTAC GTAATAACGCTAAAAGTTTTAGATGTGCTTTACTAAGTCAT CGCGATGGAGCAAAAGTACATTTAGGTACACGGCCTACA GAAAAACAGTATGAAACTCTCGAAAATCAATTAGCCTTTTT ATGCCAACAAGGTTTTTCACTAGAGAATGCATTATATGCA CTCAGCGCTGTGGGGCATTTTACTTTAGGTTGCGTATTG GAAGATCAAGAGCATCAAGTCGCTAAAGAAGAAAGGGAA ACACCTACTACTGATAGTATGCCGCCATTATTACGACAAG CTATCGAATTATTTGATCACCAAGGTGCAGAGCCAGCCTT | [5] |

| | | | |
|---|---|---|---|
| | | CTTATTCGGCCTTGAATTGATCATATGCGGATTAGAAAAA CAACTTAAATGTGAAAGTGGGTCCTAA | |
| *lacI* | coding sequence | ATGAAACCAGTAACGTTATACGATGTCGCAGAGTATGCC GGTGTCTCTTATCAGACCGTTTCCCGCGTGGTGAACCAG GCCAGCCACGTTTCTGCGAAAACGCGGGAAAAAGTGGAA GCGGCGATGGCGGAGCTGAATTACATTCCCAACCGCGT GGCACAACAACTGGCGGGCAAACAGTCGTTGCTGATTGG CGTTGCCACCTCCAGTCTGGCCCTGCACGCGCCGTCGC AAATTGTCGCGGCGATTAAATCTCGCGCCGATCAACTGG GTGCCAGCGTGGTGGTGTCGATGGTAGAACGAAGCGGC GTCGAAGCCTGTAAAGCGGCGGTGCACAATCTTCTCGCG CAACGCGTCAGTGGGCTGATCATTAACTATCCGCTGGAT GACCAGGATGCCATTGCTGTGGAAGCTGCCTGCACTAAT GTTCCGGCGTTATTTCTTGATGTCTCTGACCAGACACCCA TCAACAGTATTATTTTCTCCCATGAGGACGGTACGCGACT GGGCGTGGAGCATCTGGTCGCATTGGGTCACCAGCAAAT CGCGCTGTTAGCGGGCCCATTAAGTTCTGTCTCGGCGCG TCTGCGTCTGGCTGGCTGGCATAAATATCTCACTCGCAAT CAAATTCAGCCGATAGCGGAACGGGAAGGCGACTGGAG TGCCATGTCCGGTTTTCAACAAACCATGCAAATGCTGAAT GAGGGCATCGTTCCCACTGCGATGCTGGTTGCCAACGAT CAGATGGCGCTGGGCGCAATGCGCGCCATTACCGAGTC CGGGCTGCGCGTTGGTGCGGATATCTCGGTAGTGGGAT ACGACGATACCGAAGATAGCTCATGTTATATCCCGCCGTT AACCACCATCAAACAGGATTTTCGCCTGCTGGGGCAAAC CAGCGTGGACCGCTTGCTGCAACTCTCTCAGGGCCAGG CGGTGAAGGGCAATCAGCTGTTGCCAGTCTCACTGGTGA AAAGAAAAACCACCCTGGCGCCCAATACGCAAACCGCCT CTCCCCGCGCGTTGGCCGATTCATTAATGCAGCTGGCAC GACAGGTTTCCCGACTGGAAAGCGGGCAGTGA | 5 |
| *sp-dcas9* | coding sequence | TTAGTCACCTCCTAGCTGACTCAAATCAATGCGTGTTTCA TAAAGACCAGTGATGGATTGATGGATAAGAGTGGCATCT AAAACTTCTTTTGTAGACGTATATCGTTTACGATCAATTGT TGTATCAAAATATTTAAAAGCAGCGGGAGCTCCAAGATTC GTCAACGTAAATAAATGAATAATATTTTCTGCTTGTTCACG TATTGGTTTGTCTCTATGTTTGTTATATGCACTAAGAACTT TATCTAAATTGGCATCTGCTAAAATAACACGCTTAGAAAAT TCACTGATTTGCTCAATAATCTCATCTAAATAATGCTTATG CTGCTCCACAAACAATTGTTTTTGTTCGTTATCTTCTGGAC TACCCTTCAACTTTTCATAATGACTAGCTAAATATAAAAAA TTCACATATTTGCTTGGCAGAGCCAGCTCATTTCCTTTTT GTAATTCTCCGGCACTAGCCAGCATCCGTTTACGACCGT TTTCTAACTCAAAAAGACTATATTTAGGTAGTTTAATGATT AAGTCTTTTTTAACTTCCTTATATCCTTTAGCTTCTAAAAA GTCAATCGGATTTTTTTCAAAGGAACTTCTTTCCATAATTG TGATCCCTAGTAACTCTTTAACGGATTTTAACTTCTTCGAT TTCCCTTTTTCCACCTTAGCAACCACTAGGACTGAATAAG CTACCGTTGGACTATCAAAACCACCATATTTTTTTGGATC CCAGTCTTTTTTACGAGCAATAAGCTTGTCCGAATTTCTTT TTGGTAAAATTGACTCCTTGGAGAATCCGCCTGTCTGTAC TTCTGTTTTCTTGACAATATTGACTTGGGGCATGGACAAT ACTTTGCGCACTGTGGCAAAATCTCGCCCTTTATCCCAGA CAATTTCTCCAGTTTCCCCATTAGTTTCGATTAGAGGGCG TTTGCGAATCTCTCCATTTGCAAGTGTAATTTCTGTTTTGA AGAAGTTCATGATATTAGAGTAAAAGAAATATTTTGCGGT TGCTTTGCCTATTTCTTGCTCAGACTTAGCAATCATTTTAC | 15 |

| | | |
|---|---|---|
| | | GAACATCATAAACTTTATAATCACCATAGACAAACTCCGA<br>TTCAAGTTTTGGATATTTCTTAATCAAAGCAGTTCCAACGA<br>CGGCATTTAGATACGCATCATGGGCATGATGGTAATTGTT<br>AATCTCACGTACTTTATAGAATTGGAAATCTTTTCGGAAGT<br>CAGAAACTAATTTAGATTTTAAGGTAATCACTTTAACCTCT<br>CGAATAAGTTTATCATTTTCATCGTATTTAGTATTCATGCG<br>ACTATCCAAAATTTGTGCCACATGCTTAGTGATTTGGCGA<br>GTTTCAACCAATTGGCGTTTGATAAAACCAGCTTTATCAA<br>GTTCACTCAAACCTCCACGTTCAGCTTTCGTTAAATTATC<br>AAACTTACGTTGAGTGATTAACTTGGCGTTTAGAAGTTGT<br>CTCCAATAGTTTTTCATCTTTTTGACTACTTCTTCACTTGG<br>AACGTTATCCGATTTACCACGATTTTTATCAGAACGCGTT<br>AAGACCTTATTGTCTATTGAATCGTCTTTAAGGAAACTTTG<br>TGGAACAATGGCATCGACATCATAATCACTTAAACGATTA<br>ATATCTAATTCTTGGTCCACATACATGTCTCTTCCATTTTG<br>GAGATAATAGAGATAGAGCTTTTCATTTTGCAATTGAGTA<br>TTTTCAACAGGATGCTCTTTAAGAATCTGACTTCCTAATTC<br>TTTGATACCTTCTTCGATTCGTTTCATACGCTCTCGCGAA<br>TTTTTCTGGCCCTTTTGAGTTGTCTGATTTTCACGTGCCAT<br>TTCAATAACGATATTTTCTGGCTTATGCCGCCCCATTACTT<br>TGACCAATTCATCAACAACTTTTACAGTCTGTAAAATACCT<br>TTTTTAATAGCAGGGCTACCAGCTAAATTTGCAATATGTT<br>CATGTAAACTATCGCCTTGTCCAGACACTTGTGCTTTTTG<br>AATGTCTTCTTTAAATGTCAAACTATCATCATGGATCAGCT<br>GCATAAAATTGCGATTGGCAAAACCATCTGATTTCAAAAA<br>ATCTAATATTGTTTTGCCAGATTGCTTATCCCTAATACCAT<br>TAATCAATTTTCGAGACAAACGTCCCCAACCAGTATAACG<br>GCGACGTTTAAGCTGTTTCATCACCTTATCATCAAAGAGG<br>TGAGCATATGTTTTAAGTCTTTCCTCAATCATCTCCCTATC<br>TTCAAATAAGGTCAATGTTAAAACAATATCCTCTAAGATAT<br>CTTCATTTTCTTCATTATCCAAAAAATCTTTATCTTTAATAA<br>TTTTTAGCAAATCATGGTAGGTACCTAATGAAGCATTAAAT<br>CTATCTTCAACTCCTGAAATTTCAACACTATCAAAACATTC<br>TATTTTTTTGAAATAATCTTCTTTTAATTGCTTAACGGTTAC<br>TTTTCGATTTGTTTTGAAGAGTAAATCAACAATGGCTTTCT<br>TCTGTTCACCTGAAAGAAATGCTGGTTTTCGCATTCCTTC<br>AGTAACATATTTGACCTTTGTCAATTCGTTATAAACCGTAA<br>AATACTCATAAAGCAAACTATGTTTTGGTAGTACTTTTTCA<br>TTTGGAAGATTTTTATCAAAGTTTGTCATGCGTTCAATAAA<br>TGATTGAGCTGAAGCACCTTTATCGACAACTTCTTCAAAA<br>TTCCATGGGGTAATTGTTTCTTCAGACTTCCGAGTCATCC<br>ATGCAAAACGACTATTGCCACGCGCCAATGGACCAACAT<br>AATAAGGAATTCGAAAAGTCAAGATTTTTTCAATCTTCTCA<br>CGATTGTCTTTTAAAAATGGATAAAAGTCTTCTTGTCTTCT<br>CAAAATAGCATGCAGCTCACCCAAGTGAATTTGATGGGG<br>AATAGAGCCGTTGTCAAAGGTCCGTTGCTTGCGCAGCAA<br>ATCTTCACGATTTAGTTTCACCAATAATTCCTCAGTACCAT<br>CCATTTTTTCTAAAATTGGTTTGATAAATTTATAAAATTCTT<br>CTTGGCTAGCTCCCCCATCAATATAACCTGCATATCCGTT<br>TTTTGATTGATCAAAAAAGATTTCTTTATACTTTTCTGGAA<br>GTTGTTGTCGAACTAAAGCTTTTAAAAGAGTCAAGTCTTG<br>ATGATGTTCATCGTAGCGTTTAATCATTGAAGCTGATAGG<br>GGAGCCTTAGTTATTTCAGTATTTACTCTTAGGATATCTGA<br>AAGTAAAATAGCATCTGATAAATTCTTAGCTGCCAAAAAC<br>AAATCAGCATATTGATCTCCAATTTGCGCCAATAAATTATC<br>TAAATCATCATCGTAAGTATCTTTTGAAAGCTGTAATTTAG |

| | | | |
|---|---|---|---|
| | | CATCTTCTGCCAAATCAAAATTTGATTTAAAATTAGGGGTC<br>AAACCCAATGACAAAGCAATGAGATTCCCAAATAAGCCAT<br>TTTTCTTCTCACCGGGGAGCTGAGCAATGAGATTTTCTAA<br>TCGTCTTGATTTACTCAATCGTGCAGAAAGAATCGCTTTA<br>GCATCTACTCCACTTGCGTTAATAGGGTTTTCTTCAAATA<br>ATTGATTGTAGGTTTGTACCAACTGGATAAATAGTTTGTC<br>CACATCACTATTATCAGGATTTAAATCTCCCTCAATCAAAA<br>AATGACCACGAAACTTAATCATATGCGCTAAGGCCAAATA<br>GATTAAGCGCAAATCCGCTTTATCAGTAGAATCTACCAAT<br>TTTTTTCGCAGATGATAGATAGTTGGATATTTCTCATGATA<br>AGCAACTTCATCTACTATATTTCCAAAAATAGGATGACGTT<br>CATGCTTCTTGTCTTCTTCCACCAAAAAAGACTCTTCAAG<br>TCGATGAAAGAAACTATCATCTACTTTCGCCATCTCATTT<br>GAAAAAATCTCCTGTAGATAACAAATACGATTCTTCCGAC<br>GTGTATACCTTCTACGAGCTGTCCGTTTGAGACGAGTCG<br>CTTCCGCTGTCTCTCCACTGTCAAATAAAAGAGCCCCTAT<br>AAGATTTTTTTTGATACTGTGGCGGTCTGTATTTCCCAGA<br>ACCTTGAACTTTTTAGACGGAACCTTATATTCATCAGTGAT<br>CACCGCCCATCCGACGCTATTTGTGCCGATAGCTAAGCC<br>TATTGAGTATTTCTTATCCAT | |
| *fn-dcas12a* | coding sequence | TTAGTTGTTTCTATTCTGAACAAATTCAAAATACTCTTCGT<br>TTTTTATTACCAAATTCAACTTCTTTCCTTCCTGGTTATTTT<br>TGATTCTACCCAACAACATTAAGCCCTTCAAACCGATATG<br>GTAAGCACCGTTCGCATCCGCGTCCTGAGGCATGTTTTT<br>AGGAGCTTGTCTACTATCGAAAAAGTTGCCATTGACGTCA<br>GCCACCGGACTGATTAAGTAATCCAACTCTGTTCCTGTCT<br>TAGAGTTTCGCATCTGCAAGATAGTGTTTAATACAGATGT<br>CAATTTTGCGAAAAATTTTTTGTCAGATTCTCCACAAATTG<br>CTGCTTTAATGCACTCGCCGTGGCCGTATTCTATAGAATA<br>GTCCTTTAATAATTTTTCCAATTCTTTCGTCGGATACACCT<br>CACGAGTATCCCAGTTATGGTTCTTATCTGAGTTTCTAAA<br>GTTGATCAATCGAGAGCCAAAAGATGCAATAGTCCATTTG<br>CCTTTAGCTGCTTTATCGCCGAAATTTTTGTAATCAAATGA<br>GAACTCGAAGTATCCTTTGTCTAAATTGTAACAAATTTTAT<br>CAAACTTAGAGAAAAACTCCTGACTTTTAGACACACTCTC<br>GTATTTCGGGTACAACTGGTTCACGAATCCCGTTACTGGA<br>CATATCTTACTTGTAAATCCAGCAGGTACATAATAGATAAT<br>ACCTGTTTGCTTGCCCATTTTCTTGAACGTCTCAAAAGGA<br>GCCGTCAATTGGTATGCACGTAAGACGCCGCCAGTTTTG<br>TCAAATTCGTTGTCTTTAAATACTAAATAATTCAACTTCTC<br>AATTAACATTTTTTCCAATTTCTGGTACACCTGTTTCTCAA<br>CCTTGAATCTTCCTCTTTTAAAGCCGAAGTTTAAATCTTCG<br>AATACCACAATCGCGTTATATTCGATCACCAATTTTGCTAT<br>TTCGTGTACTACCTGTGATAAGTATCCCTCCTTCATCTCC<br>TTAATATTATTAATCTTTTTCCAATCTTTACGAGCAGAATC<br>TCTGTCCTTTTCGATAGCTGCCAACTTATCGTGATAATTT<br>GTTTTCATACGATCATTTCCAATTATATTAAATGTATCCTG<br>TTTGATAATGTTACCCTTGCCATCAACTAACGTATAATAAG<br>CCAAGTGTCGTTCGCCTCTCGCGATAGACAATATGTGGA<br>CGTCGTTAGCCTTTTCCTTCAACAATAAATTGATTTCGTC<br>GTTAAATTTATTCGCTCCAGAACTCTTAAAATTAATAGTGA<br>TAGGACAATGGAAAAAAAACTTATCCTCTGTAAATCGCTT<br>ATCTTTTATTAAGTCATACTCAAATACAGACTCTTTTTTTG<br>GATTGTCTTTGTTTTTGTTCGCGATCGCCTCTTTCGCAGG<br>GTGTGTGATTTTCTTTGGGATAGACTGCTTTCGGTAAAAT<br>AACTCTGCCTCTCCATTTAACTTATACACTACATCTTGCAA | Codon optimized for *S. aureus.* Amino acid sequence from [18] |

| | | |
|---|---|---|
| | | GTTACGTTCATCGAACAAAGCTTTCCAGTACAACGTATGT<br>AAGTTAGGTCGTCCCTTACTGTACGCAGAGAAATCTTTAT<br>TATAGATTTGAAACAAGTACAATTTTCCTTGATTCACCACA<br>GAATCAATATAAGATTCTGATATGTTTTCAAACGTCAACTT<br>GTAACCCTGATTCTCTACTTCTCTGTAAAACTCGTCGATA<br>CTGTTGTATCTCTGAGTGTCACTGAATCGGAAACCGAAGT<br>CCTTCCACTCAGGATGTTTACTAATTGACTGCTTATAAAAA<br>TCTATGAATTTTCGACAGTCCTCAATATTGAACTCGAATTT<br>TTCGTAGCCTTTTTGAGGAGAGCCATTCTTAGTATGAGTA<br>CTATGATTACGAATTCTCAAAATGTCCTCTGATGGATTATA<br>AAACTTTATACTTTTAGCTGAGAAGAAAACCTTTGGTAACA<br>TTTTATTAGCGCCCGGTAACAACTTATAAACTATCTTCTTA<br>TAACCCTCGCCTTTATTTTCCTTGATTGCTTTGTCATCAAA<br>TATTTTATTATTCTTTTTATTCATGACACCCAAATAGTATTT<br>GTCGTCCTTGATAAACAAAATTGCCGTATTATCCGGCTCT<br>TTGTTTTTATCCCAACCATTAGCCAAAGTTGAATTCTCAAA<br>ATTTAATTTGAATTTCTCGTCAGAGTAAGGCTTTTGAGTAA<br>TGTAATTTCGTATCTTATTGTACAATGGTACTATGTTAGCT<br>AACTCAAAATAGCATTCTTCAAAGACTAAGTAGAAGTGCT<br>CGTCCTTATCCAAGATGTTCGCCTTGTCCTCTGACTGTGA<br>TATGTGAAATATCTTCAACTTGTGCAATAAGTTGTTTGTTT<br>GGTCTAACAAGTCCTTGATTGCTTTTACATCATCTTCCGC<br>TGACGCCTGCAACAAATCTTTCTTACCTTGGTTCTGATAC<br>TTAATTGAAATTTGCGCTAAGTTATCCTTATTCTGTGCAAT<br>TTCATCGAAAATCATTGGAATTGCCGCAAAGTTCGCTAAA<br>ATTTCCTCGAAACGACACTGTTTGTCTATATCACGATGCT<br>TATTGAACTCCTCTAACGCCAATTTGATTGTTTCCAAAGAT<br>AAGTACTTCGCTTTCTCCGTCTTTTTCGCTATCAACTCCT<br>GCTCCTTTTTACTAGGATTATCTAAATTCTTCGGCGCTATT<br>TGCTGTGTGATATACTCTAAGACTGCCGTTCCGATCACAC<br>TGTAATCGTCGAACACTTGTTGAGATAAGTCCGTCAAAGA<br>CTTGTCGTTCTTGAAATAAATTTTACTTAAATCTAATTTTTG<br>CGCCTTTAAGTCATCGAACAACAAACTTAAAGTCTCTTTTA<br>TACTTTTTTCTTCTACTGTTTTGAATGCTGCTATTTGCTCA<br>TAGAAAGACTGCATTGTTGTCACCACGTCTGAGTCGTCCT<br>CCAACTTGTCAATTACAAAACTCTTTGATTCTGTATCACTT<br>AATATTTGTTTAAATAAGACAGACATTTTATACTTCTTTAAT<br>GTTTTGTCATTTATCTGTTGACTATACAAGTTGATATATTC<br>ATTTATTCCCTTTCGTTTAGTGTTCTCGCCGTTTACGAATT<br>TACCTCCTATGATCGTGTTGAACTTAGTAATACCTGATTG<br>GTTTAAATAATTGTTGAAATTCGCGATCTCGAATACCTCAT<br>CTAAAGAAAACACTCTCTGGTTAACTTCAGACGTTTTATA<br>GTCAATATCGAAAGTTAACTCCTCCGCCAAGTCTTTCTTT<br>ATTTGTTCATAATTTATTGCCTCCGGCGCCTTGTCTTTTAA<br>TGATTCGTACTTCGCCTTATTCTCTAAAAATTTCGGCAAAT<br>TGTCGTCCACAATTCTGTAAATAATACTTGTAGGTATATCA<br>TTAGATGAGTAGACATTTTTACGATTCTCATGGAATCCCTT<br>GAAATATGTTGTCCATCCCTTAAAACTTTTGATGATCTCTA<br>ATGCCTCGTCAATGTCAGTGATATCACTATTTGCCTTGAA<br>CAACTCTATTCCGTTGTCTTTACTTTGTTTCAACCATAAGA<br>TTAAATCAGATTCTTGTCCCTTCTTTGCATCAATCAAATTC<br>TGATTGAATAAGTTCTTAAATTTTTCAGAGTCTTTAATGTA<br>TTCAGAGATCTGTTTTTTGATCGTATCTTTCGCTGATTTGA<br>AATCTTTTTGCAAGTTGTCGTCGTCACTTTTTTTCAATTTG<br>AAGTACACATCTGAATAATTCTGCAATAAGTCCTCACTAAT<br>GCAAACACTAGATAATATCTCTTCAATAAAAAATTGATGGT |

| | | | |
|---|---|---|---|
| | | ATTTGTCAATTATCTGCTTCGCCTTTTTGTAGTCTTTCGCT<br>CTTTTTTCGTCGTCTAAAATCAAACCACGAGCCTTGATATT<br>CTCCAATGTTTTTCCTTGTGGAATCAACTCAAATCTCAATG<br>TCTTTGACAATGAGTATTTGTTGACGAATTCTTGATAGATA<br>GACAT | |
| *lb-dcas12a* | coding sequence | CTAATGTTTGACAGATGTTTGCGCATATTCCAACCACTCT<br>TTATTTGAGATTGCTATCTTTACCTTATCTAATTTTTCATCT<br>TCTGCCTTTTTGAATTGACCGATTGCCCATAAGACCTTAC<br>GAGCAATATTGTATGCTCCATTTGCATCCGCGTTCTTCGG<br>CAATATAGCATTTTCCTGCGCTTCATAATTTCGACTGTCG<br>TAGAATATACCGTCTGAATTCTTGACCGGAGATATTAAGA<br>AATCCACATCTGTACGACCCGTAATACTGTTTCGCATTTG<br>CAACATTAAAGACATTAACGCCATGAAAGAACTATAGAAC<br>GCCTTGTCAGATTGTTCACACAACAATGCTCGTATATCTC<br>CTTGCTGATAATTGATTCCATATTTGTTGAACAATTCCTTG<br>TAAGCTGACGTTAAGCAAACTTCTTCCCAATCGAACACGT<br>TATTTTTTTTTGGGTTTCTAAATATACGGATTCGGTTTCCA<br>TATGAATACAATTTCCACTTTTTGATGTAATCCGCATCCGT<br>ACGTGAGAAGTTCTTATAGTCTAATGCGAATTCAAATAAG<br>TCCTCTTCTGGAACATACATGATACGATCGAATGAAGAAA<br>TGAATTTTTTAGAGTCCGCAATAGACGTATACTTCGTCTTT<br>AATAAATTTACAAAGCCTGTTGACGGGTCAATTTTACTCG<br>TCAACCACGCTGGGATGTAAAAAATAAAGCCATTCTGAGT<br>ACTCATTGATTTAAAACTCTCGAACTTATTTGTGATTTGGT<br>AGCCCTTTAACGCTCCGCCAGTAGCACACGGATTTGATTT<br>TTTATCGACCATGTAATTCAACTTATCTATTAACATTTTCT<br>CAAATTTTTGGTACACCTGTTTTTCTACTTTAACTCTTGAA<br>TTTTTAAAGCCACTATTTAAATCTTCTAAAGCGATTACTGC<br>GTCGTATTTCTCGACTAACTCGCAAATCTTATGGACTACC<br>TGAGATATATAACCTGCCTTCAACTCTTTGATATTCTCTAT<br>ACTTGTCCAGTTCTGTCTTGCCTCAAATCGTTCTTTCTCCT<br>TTTTATCTAATAAACTATGATAGTCTGTCTTTATTCTGATA<br>CCGTTGAAGTTATTAATAATTTCGTTTAATGAGTACTGTTC<br>GACGATGTTTCCTTTTCCATCGACCACAACGATATATAAC<br>AAGTTTCGCTCACCTCTCGCTATGCCGATTACATACGGGT<br>TATCATCGTGCTTCAACAAAACTCGCACTTCTGTATTAATC<br>TTGAATATGTTTTTAGGGCACTTATTGATTGCTATAGGTAT<br>GTGCAATTCGTATTGATCTTCTGAGAATCGCTTATCCTTAT<br>ACACGTCGTAAGACAATGTAGTTGTTTTTTTTGGATTGTC<br>CGGATTCTTGTTAGCAATAGGTGAATTAGCTGGGTGAACT<br>ACTAACTCCTCCTTTTTCAAACTCGCTCGTCGCATAAACA<br>ACTCCGCACCTCCAGATAATCGTATCTGTCCATGGTTGTT<br>TTCATCAAACAACAATTTGAAGTACATCGTATGCAAGTTA<br>GGAGTGCCATGAGATTTATCACTAAAGTCCTTATTGTAAA<br>TCTGAAACATGTACAACTTACCTTCCTCAACCAACTTGTC<br>TACTTCTTTTTTAGACGCACTTTCGAAAGATACCTTATAGC<br>CCTGCTCTTCCACTTCTCTATAAAATCCAGCTATATCCTTG<br>TACTTTTCTGTTTCACTAAAGTTAAAGTCATAAGCATTACT<br>CCACTTTGGATAACGTGATATAGAATCCTTGAAGAAATCA<br>ATTAACTTGTGGCAGTCATTTAAGTTAAACATATCACCTTT<br>TTTGAACGTACCATTTTTGTAAATTTTCTGAATGTCCTCAG<br>AAGGGTTATAATAAGCCATCCATTTTTTACTAAAGAACACT<br>TTCGGTAACATTTTATTCGGGCCAGGCAATAACTTGTAGT<br>TGATTTTCTCATAGTTACCGTTTACGTCATCCTTATCAATT<br>TTCTGTAAGCATTTTGCGTATTTTTTGTCCATTATCGCTAA<br>GTAGTATTTAGAACCATATCGCAAGATTGTTGCTCGGTAA | Codon optimized for *S. aureus.* Amino acid sequence from [18] |

| | | | |
|---|---|---|---|
| | | TCAGTTTCCTTGTCCTTATCCCAACCGCCCATGAATTGAG GATTCTGAAAGTACAACTTAAACTTGTCCTTTGAGTATGG TTTTTGTGTCACATAATTACGAATCGCATCATAAATATGGT CAACCTTCAATAAAATGTCATATGCTAAAACGAAATCACC GTAGAAACTTTCGTCACGGTTCGTCTCTTTGCCTTCTCCG AAGAAAGCTTTAATGTAGTTTTCGAAACTCTTAACTGAGT CCAATAAATCTTTCATTATAGCGACAACAGCATCATTTTTT TTCAAAGATTTTTCTAAAACGAAGTCAGCATCAAACAACTT CTCAGAACTACCGTAAACTTTGTATATCTCATCGACCTTTT GTATGATAATTTCTTTCAACTTCTCCACAACACTCAAATCT GCATCCGCATATTCCTGCAACTGCTCCAAAGAGAAAGAG CCTATTTTCTTAAATGACTTACGTCGATCATCTTCGTATTT CTCTGTTACTACCGCTTTCTTCTTTAAGTGTATGTCGTCAT ATTCAGCATTCCACTTATCACGAATAACATTCCATTCACCA AAAATGTCCTTACTAATCGTTGAGATTGCCGGTCCATTCT TCACGAATATGCCCGCACTTGAGTATTCATCGAAGTTCTT AAACAATTTCTCTAACTTCTTGATAGATGAAAAAATTTCAC TGTTCTTGTTTAATGTATTTCGAAAGACTTCCAACACTTCC TCATCAGACGTGTATCCCTCACCGTAGAAAGATAAAGATT CTCGGTCACTCAAAACTTGCTTATACAACGGCTTGAATTT AGGTAACTTTTGTTTTGTCTTTTGGTTGTACAAGTTTATGT ATTCGTTCAAGCCTTTTATCTTTTCGCCAGATTCTGTGAC GAATCCACCTATTATCGCATTATACACGTCGATGCCTTCC TGTGTTAACACAAAATTAAAAAACTCTCCTTCAAAAAAATC CTCGACGTCATAGTCAGAGTTTAAAATTTTTTCTTTAATCT CTTGAACCTCGTGCTTGTCAAATATTGCGTCAACTTTCTC GAAAATATCCATGTTACTGATGTATCTAGTTAAGTTTTCGT TGATGCATCTAAAAGCTATTGACGTTGATTTCGCTTCCTC ACTAAACATGTTCTCTCTGTTGTCAAAGAAACCAGTGAAA GCCGTCGTAAAGCCGTTGAAACTATTGACTAATGCGATCT CGTCTTTATCATCCAAAAATTCTGGTAATATCGTCTCAATA ATATCTTTTTTAAATAAACTTTTGTAACCTTCATTGCCTTTA AAAGCCTTAGCTATCTCCTTTCGTAAATTAATTTCCAAATT CTCCAATTCCTTGTTCTCTTTCTCTGTTCTTGTCTTTTTTCT GAATAAAGATATGTAATTATTCAAGTTTTTTAACTTAATTG AATGTAAAACATCGTTGATGAAACTCAAATAATAACGATC CAACAACTTCTTGACACCCTTATAATCCTCAGCTCTCTTTT CATCTTCCACCAATAAACGTTTATTGTCTATATTTTCTTGA GTCTTGCCTACCGGAATCGCTTTAAAACGCAATGTCTTTG ATAATGAGTAGCAATTTGTAAATTTTTCTAACTTTGACAT | |
| *kanR* | coding sequence | ATGACCATGATTACGCCAAGCTTGCATGCCTGCAGGTCG ACTCTAGAGGATCCCCGGGTACCGAGCTCGAATTCACTG GCCGTCGTTTTACAACGTCGTGACTGGGAAAACCCTGGC GTTACCCAACTTAATCGCCTTGCAGCACATCCCCCTTTCG CCAGCTGGCGTAATAGCGAAGAGGCCCACACCGATCGC CCTTCCCAACAGTTGCGCAGCCTGAATGGCTAA | [3] |
| *specR* | coding sequence | TTAGAAAAACTCATCGAGCATCAAATGAAACTGCAATTTA TTCATATCAGGATTATCAATACCATATTTTTGAAAAAGCCG TTTCTGTAATGAAGGAGAAAACTCACCGAGGCAGTTCCAT AGGATGGCAAGATCCTGGTATCGGTCTGCGATTCCGACT CGTCCAACATCAATACAACCTATTAATTTCCCCTCGTCAA AAATAAGGTTATCAAGTGAGAAATCACCATGAGTGACGAC TGAATCCGGTGAGAATGGCAAAAGCTTATGCATTTCTTTC CAGACTTGTTCAACAGGCCAGCCATTACGCTCGTCATCA AAATCACTCGCATCAACCAAACCGTTATTCATTCGTGATT GCGCCTGAGCGAGACGAAATACGCGATCGCTGTTAAAAG | [3] |

| | | | |
|---|---|---|---|
| | | GACAATTACAAACAGGAATCGAATGCAACCGGCGCAGGA ACACTGCCAGCGCATCAACAATATTTTCACCTGAATCAGG ATATTCTTCTAATACCTGGAATGCTGTTTTCCCGGGGATC GCAGTGGTGAGTAACCATGCATCATCAGGAGTACGGATA AAATGCTTGATGGTCGGAAGAGGCATAAATTCCGTCAGC CAGTTTAGTCTGACCATCTCATCTGTAACATCATTGGCAA CGCTACCTTTGCCATGTTTCAGAAACAACTCTGGCGCATC GGGCTTCCCATACAATCGATAGATTGTCGCACCTGATTG CCCGACATTATCGCGAGCCCATTTATACCCATATAAATCA GCATCCATGTTGGAATTTAATCGCGGCCTGGAGCAAGAC GTTTCCCGTTGAATATGGCTCAT | |
| *lacZα* | coding sequence | ATGAGGGAAGCGGTGATCGCCGAAGTATCGACTCAACTA TCAGAGGTAGTTGGCGTCATCGAGCGCCATCTCGAACCG ACGTTGCTGGCCGTACATTTGTACGGCTCCGCAGTGGAT GGCGGCCTGAAGCCACACAGTGATATTGATTTGCTGGTT ACGGTGACCGTAAGGCTTGATGAAACAACGCGGCGAGCT TTGATCAACGACCTTTTGGAAACTTCGGCTTCCCCTGGAG AGAGCGAGATTCTCCGCGCTGTAGAAGTCACCATTGTTG TGCACGACGACATCATTCCGTGGCGTTATCCAGCTAAGC GCGAACTGCAATTTGGAGAATGGCAGCGCAATGACATTC TTGCAGGTATCTTCGAGCCAGCCACGATCGACATTGATC TGGCTATCTTGCTGACAAAAGCAAGAGAACATAGCGTTG CCTTGGTAGGTCCAGCGGCGGAGGAACTCTTTGATCCGG TTCCTGAACAGGATCTATTTGAGGCGCTAAATGAAACCTT AACGCTATGGAACTCGCCGCCCGACTGGGCTGGCGATG AGCGAAATGTAGTGCTTACGTTGTCCCGCATTTGGTACA GCGCAGTAACCGGCAAAATCGCGCCGAAGGATGTCGCT GCCGACTGGGCAATGGAGCGCCTGCCGGCCCAGTATCA GCCCGTCATACTTGAAGCTAGACAGGCTTATCTTGGACA AGAAGAAGATCGCTTGGCCTCGCGCGCAGATCAGTTGGA AGAATTTGTCCACTACGTGAAAGGCGAGATCACCAAGGT AGTCGGCAAATAA | [8] |
| p15a stringent | plasmid replicon | GACCTCAGCGCTAGCGGAGTGTATACTGGCTTACTATGT TGGCACTGATGAGGGTGTCAGTGAAGTGCTTCATGTGGC AGGAGAAAAAAGGCTGCACCGGTGCGTCAGCAGAATATG TGATACAGGATATATTCCGCTTCCTCGCTCACTGACTCGC TACGCTCGGTCGTTCGACTGCGGCGAGCGGAAATGGCTT ACGAACGGGGCGGAGATTTCCTGGAAGATGCCAGGAAG ATACTTAACAGGGAAGTGAGAGGGCCGCGGCAAAGCCG TTTTTCCATAGGCTCCGCCCCCCTGACAAGCATCACGAA ATCTGACGCTCAAATCAGTGGTGGCGAAACCCGACAGGA CTATAAAGATACCAGGCGTTTCCCCCTGGCGGCTCCCTC GTGCGCTCTCCTGTTCCTGCCTTTCGGTTTACCGGTGTC ATTCCGCTGTTATGGCCGCGTTTGTCTCATTCCACGCCTG ACACTCAGTTCCGGGTAGGCAGTTCGCTCCAAGCTGGAC TGTATGCACGAACCCCCCGTTCAGTCCGACCGCTGCGCC TTATCCGGTAACTATCGTCTTGAGTCCAACCCGGAAAGAC ATGCAAAAGCACCACTGGCAGCAGCCACTGGTAATTGAT TTAGAGGAGTTAGTCTTGAAGTCATGCGCCGGTTAAGGC TAAACTGAAAGGACAAGTTTTGGTGACTGCGCTCCTCCAA GCCAGTTACCTCGGTTCAAAGAGTTGGTAGCTCAGAGAA CCTTCGAAAAACCGCCCTGCAAGGCGGTTTTTTCGTTTTC AGAGCAAGAGATTACGCGCAGACCAAAACGATCTCAAGA AGATCATCTTATTAA | [3] |
| pSC101 | plasmid replicon | CTGTCAGACCAAGTTTACGAGCTCGCTTGGACTCCTGTT GATAGATCCAGTAATGACCTCAGAACTCCATCTGGATTTG | [3] |

| | | |
|---|---|---|
| | | TTCAGAACGCTCGGTTGCCGCCGGGCGTTTTTTATTGGT<br>GAGAATCCAAGCACTAGGGACAGTAAGACGGGTAAGCCT<br>GTTGATGATACCGCTGCCTTACTGGGTGCATTAGCCAGT<br>CTGAATGACCTGTCACGGGATAATCCGAAGTGGTCAGAC<br>TGGAAAATCAGAGGGCAGGAACTGCTGAACAGCAAAAAG<br>TCAGATAGCACCACATAGCAGACCCGCCATAAAACGCCC<br>TGAGAAGCCCGTGACGGGCTTTTCTTGTATTATGGGTAG<br>TTTCCTTGCATGAATCCATAAAAGGCGCCTGTAGTGCCAT<br>TTACCCCCATTCACTGCCAGAGCCGTGAGCGCAGCGAAC<br>TGAATGTCACGAAAAAGACAGCGACTCAGGTGCCTGATG<br>GTCGGAGACAAAAGGAATATTCAGCGATTTGCCCGAGCT<br>TGCGAGGGTGCTACTTAAGCCTTTAGGGTTTTAAGGTCT<br>GTTTTGTAGAGGAGCAAACAGCGTTTGCGACATCCTTTTG<br>TAATACTGCGGAACTGACTAAAGTAGTGAGTTATACACAG<br>GGCTGGGATCTATTCTTTTTATCTTTTTTTATTCTTTCTTTA<br>TTCTATAAATTATAACCACTTGAATATAAACAAAAAAAACA<br>CACAAAGGTCTAGCGGAATTTACAGAGGGTCTAGCAGAA<br>TTTACAAGTTTTCCAGCAAAGGTCTAGCAGAATTTACAGA<br>TACCCACAACTCAAAGGAAAAGGACATGTAATTATCATTG<br>ACTAGCCCATCTCAATTGGTATAGTGATTAAAATCACCTA<br>GACCAATTGAGATGTATGTCTGAATTAGTTGTTTTCAAAG<br>CAAATGAACTAGCGATTAGTCGCTATGACTTAACGGAGCA<br>TGAAACCAAGCTAATTTTATGCTGTGTGGCACTACTCAAC<br>CCCACGATTGAAAACCCTACAAGGAAAGAACGGACGGTA<br>TCGTTCACTTATAACCAATACGCTCAGATGATGAACATCA<br>GTAGGGAAAATGCTTATGGTGTATTAGCTAAAGCAACCAG<br>AGAGCTGATGACGAGAACTGTGGAAATCAGGAATCCTTT<br>GGTTAAAGGCTTTGAGATTTTCCAGTGGACAAACTATGCC<br>AAGTTCTCAAGCGAAAAATTAGAATTAGTTTTTAGTGAAG<br>AGATATTGCCTTATCTTTTCCAGTTAAAAAAATTCATAAAA<br>TATAATCTGGAACATGTTAAGTCTTTTGAAAACAAATACTC<br>TATGAGGATTTATGAGTGGTTATTAAAAGAACTAACACAA<br>AAGAAAACTCACAAGGCAAATATAGAGATTAGCCTTGATG<br>AATTTAAGTTCATGTTAATGCTTGAAAATAACTACCATGAG<br>TTTAAAAGGCTTAACCAATGGGTTTTGAAACCAATAAGTA<br>AAGATTTAAACACTTACAGCAATATGAAATTGGTGGTTGA<br>TAAGCGAGGCCGCCCGACTGATACGTTGATTTTCCAAGT<br>TGAACTAGATAGACAAATGGATCTCGTAACCGAACTTGAG<br>AACAACCAGATAAAAATGAATGGTGACAAAATACCAACAA<br>CCATTACATCAGATTCCTACCTACGTAACGGACTAAGAAA<br>AACACTACACGATGCTTTAACTGCAAAAATTCAGCTCACC<br>AGTTTTGAGGCAAAATTTTTGAGTGACATGCAAAGTAAGC<br>ATGATCTCAATGGTTCGTTCTCATGGCTCACGCAAAAACA<br>ACGAACCACACTAGAGAACATACTGGCTAAATACGGAAG<br>GATCTGAGGTTCTTATGGCTCTTGTATCTATCAGTGAAGC<br>ATCAAGACTAACAAACAAAAGTAGAACAACTGTTCACCGT<br>TAGATATCAAAGGGAAAACTGTCCATATGCACAGATGAAA<br>ACGGTGTAAAAAAGATAGATACATCAGAGCTTTTACGAGT<br>TTTTGGTGCATTTAAAGCTGTTCACCATGAACAGATCGAC<br>AATGTAAC |

[a] The Bba_B0034 RBS was used to control the expression of *sp-dcas9*.

[b] This sequence of DNA was used between the dCas12a single or dual gRNA and the L3S2P21 terminator as buffer to limit interaction of the terminator hairpin with the spacer sequence.

**Table S3. Sensor response function parameters for inducible promoters**

| Strain | Inducible promoter | $K_d$ (ng/mL or mM)[a] | n | $y_{min}$ (RPU) | $y_{max}$ (RPU) | Fold induction[b] |
|---|---|---|---|---|---|---|
| MG1655 | $P_{Tet}$ | 1.950 | 3.112 | 0.00284 | 7.445 | 2619 |
| EcN | $P_{Tet}$ | 0.963 | 2.835 | 0.00553 | 7.766 | 1403 |
| CFT073 | $P_{Tet}$ | 1.262 | 3.051 | 0.00318 | 8.051 | 2529 |
| UMN026 | $P_{Tet}$ | 2.474 | 2.313 | 0.00863 | 7.092 | 822 |
| MG1655 | $P_{Tac}$ | 0.296 | 1.701 | 0.00768 | 5.814 | 757 |
| EcN | $P_{Tac}$ | 0.0199 | 3.508 | 0.01112 | 3.496 | 314 |
| CFT073 | $P_{Tac}$ | 0.268 | 1.782 | 0.00636 | 4.596 | 722 |
| UMN026 | $P_{Tac}$ | 0.158 | 2.061 | 0.01583 | 5.211 | 329 |

[a] $K_d$ for sensor with $P_{Tet}$ is in units of ng/mL aTc. $K_d$ for sensor with $P_{Tac}$ is in units of mM IPTG.

[b] Fold induction was calculated as $y_{max}$ / $y_{min}$ using the fitted parameters

**Table S4. gRNAs with significant differences in fold repression between strains**

| CRISPRi system | gRNA name | Strain pair | *p*-value[a] |
|---|---|---|---|
| Sp-dCas9 | -123N | CFT073, EcN | 0.005051 |
| Sp-dCas9 | -123N | EcN, MG1655 | 0.011155 |
| Sp-dCas9 | -89N | UMN026, CFT073 | 0.017447 |
| Sp-dCas9 | -89N | CFT073, EcN | 0.036205 |
| Sp-dCas9 | +34N | CFT073, EcN | 0.003924 |
| Sp-dCas9 | +34N | UMN026, CFT073 | 0.006277 |
| Sp-dCas9 | +664N | CFT073, MG1655 | 0.00104 |
| Sp-dCas9 | +664N | CFT073, EcN | 0.014332 |
| Sp-dCas9 | +664N | UMN026, CFT073 | 0.015031 |
| Sp-dCas9 | -144T | CFT073, EcN | 0.019228 |
| Fn-dCas12a | -119N | CFT073, MG1655 | 0.005909 |
| Fn-dCas12a | -119N | UMN026, CFT073 | 0.047519 |
| Fn-dCas12a | +131N | UMN026, CFT073 | 0.007473 |
| Fn-dCas12a | +131N | CFT073, EcN | 0.048433 |
| Fn-dCas12a | +339T | CFT073, MG1655 | 0.047045 |
| Lb-dCas12a | -119N | UMN026, EcN | 4.24E-05 |
| Lb-dCas12a | -119N | UMN026, MG1655 | 0.000488 |
| Lb-dCas12a | -119N | UMN026, CFT073 | 0.001164 |
| Lb-dCas12a | -119N | CFT073, EcN | 0.024952 |
| Lb-dCas12a | +112N | CFT073, EcN | 0.034179 |
| Lb-dCas12a | +20T | UMN026, EcN | 5.14E-06 |
| Lb-dCas12a | +20T | CFT073, EcN | 1.08E-05 |
| Lb-dCas12a | +20T | UMN026, MG1655 | 0.000138 |
| Lb-dCas12a | +20T | CFT073, MG1655 | 0.000423 |
| Lb-dCas12a | +20T | EcN, MG1655 | 0.006791 |
| Lb-dCas12a | +339T | UMN026, EcN | 0.012417 |
| Lb-dCas12a | +339T | CFT073, EcN | 0.017343 |
| Lb-dCas12a | +666T | CFT073, EcN | 0.001941 |
| Lb-dCas12a | +666T | UMN026, EcN | 0.013248 |
| Lb-dCas12a | +666T | EcN, MG1655 | 0.01782 |
| Lb-dCas12a | +139T | UMN026, EcN | 6.99E-07 |
| Lb-dCas12a | +139T | CFT073, EcN | 6.87E-05 |
| Lb-dCas12a | +139T | UMN026, MG1655 | 7.44E-05 |
| Lb-dCas12a | +139T | EcN, MG1655 | 0.000243 |
| Lb-dCas12a | +139T | UMN026, CFT073 | 0.000266 |

[a] *p*-values were determined from a one-way ANOVA with Tukey post-hoc analysis on the fold repression data across the strains for each gRNA design in each CRISPRi system, with significance determined if $p < 0.05$ (Methods).

**Table S5. Parameters for the full linear regression models of CRISPRi repression**

| CRISPRi system | Term | Estimate | Standard error | t ratio | Probability > \|t\| |
|---|---|---|---|---|---|
| Sp-dCas9 | Intercept | 6.409419 | 2.198337 | 2.92 | 0.008 |
| Sp-dCas9 | gRNA | 5.152237 | 0.906794 | 5.68 | <.0001 |
| Sp-dCas9 | gRNA X gRNA | 2.657024 | 2.23189 | 1.19 | 0.2465 |
| Sp-dCas9 | dCas | 0.273539 | 0.9068 | 0.3 | 0.7657 |
| Sp-dCas9 | dCas X dCas | -3.46166 | 2.231852 | -1.55 | 0.1352 |
| Sp-dCas9 | Day[1] | -1.42831 | 1.102192 | -1.3 | 0.2084 |
| Sp-dCas9 | Day[2] | 1.52915 | 1.102192 | 1.39 | 0.1792 |
| Sp-dCas9 | Day[3] | -0.10084 | 1.114132 | -0.09 | 0.9287 |
| Sp-dCas9 | gRNA X dCas | -0.53285 | 1.031707 | -0.52 | 0.6107 |
| Fn-dCas12 | Intercept | 39.56584 | 4.282594 | 9.24 | <.0001 |
| Fn-dCas12 | gRNA | 23.40558 | 1.766532 | 13.25 | <.0001 |
| Fn-dCas12 | gRNA X gRNA | 1.290358 | 4.34796 | 0.3 | 0.7694 |
| Fn-dCas12 | dCas | -8.08291 | 1.766542 | -4.58 | 0.0001 |
| Fn-dCas12 | dCas X dCas | -20.8642 | 4.347885 | -4.8 | <.0001 |
| Fn-dCas12 | Day[1] | -0.05223 | 2.147187 | -0.02 | 0.9808 |
| Fn-dCas12 | Day[2] | 0.050369 | 2.147187 | 0.02 | 0.9815 |
| Fn-dCas12 | Day[3] | 0.001861 | 2.170448 | 0 | 0.9993 |
| Fn-dCas12 | gRNA X dCas | -11.1782 | 2.009874 | -5.56 | <.0001 |
| Lb-dCas12a | Intercept | 22.30358 | 8.045164 | 2.77 | 0.0111 |
| Lb-dCas12a | gRNA | 21.44371 | 3.318558 | 6.46 | <.0001 |
| Lb-dCas12a | gRNA X gRNA | 1.31819 | 8.167959 | 0.16 | 0.8733 |
| Lb-dCas12a | dCas | 11.83392 | 3.318578 | 3.57 | 0.0017 |
| Lb-dCas12a | dCas X dCas | -2.15639 | 8.167819 | -0.26 | 0.7942 |
| Lb-dCas12a | Day[1] | 1.510352 | 4.033646 | 0.37 | 0.7117 |
| Lb-dCas12a | Day[2] | 1.752461 | 4.033646 | 0.43 | 0.6682 |
| Lb-dCas12a | Day[3] | -3.26281 | 4.077345 | -0.8 | 0.4321 |
| Lb-dCas12a | gRNA X dCas | 9.917087 | 3.775695 | 2.63 | 0.0154 |

**Table S6. Parameters for the reduced linear models of CRISPRi repression**

| CRISPRi system | Term | Estimate | Standard error | t ratio | Probability > \|t\| |
|---|---|---|---|---|---|
| Sp-dCas9 | Intercept | 5.819385 | 0.776301 | 7.5 | <.0001 |
| Sp-dCas9 | gRNA | 5.152231 | 0.906525 | 5.68 | <.0001 |
| Fn-dCas12a | Intercept | 40.29966 | 3.233971 | 12.46 | <.0001 |
| Fn-dCas12a | gRNA | 23.40558 | 1.660536 | 14.1 | <.0001 |
| Fn-dCas12a | dCas | -8.08291 | 1.660545 | -4.87 | <.0001 |
| Fn-dCas12a | dCas X dCas | -20.5745 | 3.960814 | -5.19 | <.0001 |
| Fn-dCas12a | gRNA X dCas | -11.1781 | 1.889024 | -5.92 | <.0001 |
| Lb-dCas12a | Intercept | 21.68892 | 2.659138 | 8.16 | <.0001 |
| Lb-dCas12a | gRNA | 21.44371 | 3.105204 | 6.91 | <.0001 |
| Lb-dCas12a | dCas | 11.83391 | 3.105223 | 3.81 | 0.0008 |
| Lb-dCas12a | gRNA X dCas | 9.867669 | 3.532478 | 2.79 | 0.0097 |

### Table S7. Sequences of gRNA designs

| Name | CRISPRi system[a] | Output gene[b,c] | Sequence[d] | PAM sequence(s) |
|---|---|---|---|---|
| -123N | Sp-dCas9 | *eyfp* | CTTTAGGACCTGTCAAAGTTGTTTTAGAGCTAGAAATAGCAAGTTAAAATAAGGCTAGTCCGTTATCAACTTGAAAAAGTGG | AGG |
| -89N | Sp-dCas9 | *eyfp* | AGCATTATAGACTATCCTTTGTTTTAGAGCTAGAAATAGCAAGTTAAAATAAGGCTAGTCCGTTATCAACTTGAAAAAGTGG | AGG |
| +34N | Sp-dCas9 | *eyfp* | GACCAGGATGGGCACCACCCGTTTTAGAGCTAGAAATAGCAAGTTAAAATAAGGCTAGTCCGTTATCAACTTGAAAAAGTGG | CGG |
| +334N | Sp-dCas9 | *eyfp* | CCTCGAACTTCACCTCGGCGGTTTTAGAGCTAGAAATAGCAAGTTAAAATAAGGCTAGTCCGTTATCAACTTGAAAAAGTGG | CGG |
| +664N | Sp-dCas9 | *eyfp* | CGGCGGTCACGAACTCCAGCGTTTTAGAGCTAGAAATAGCAAGTTAAAATAAGGCTAGTCCGTTATCAACTTGAAAAAGTGG | AGG |
| -144T | Sp-dCas9 | *eyfp* | CTAACTTTGACAGGTCCTAAGTTTTAGAGCTAGAAATAGCAAGTTAAAATAAGGCTAGTCCGTTATCAACTTGAAAAAGTGG | AGG |
| -128T | Sp-dCas9 | *eyfp* | TGCTAGCCTGAAGCTGTCACGTTTTAGAGCTAGAAATAGCAAGTTAAAATAAGGCTAGTCCGTTATCAACTTGAAAAAGTGG | CGG |
| +28T | Sp-dCas9 | *eyfp* | CGAGGAGCTGTTCACCGGGGGTTTTAGAGCTAGAAATAGCAAGTTAAAATAAGGCTAGTCCGTTATCAACTTGAAAAAGTGG | TGG |
| +325T | Sp-dCas9 | *eyfp* | CAACTACAAGACCCGCGCCGGTTTTAGAGCTAGAAATAGCAAGTTAAAATAAGGCTAGTCCGTTATCAACTTGAAAAAGTGG | AGG |
| +659T | Sp-dCas9 | *eyfp* | GCGCGATCACATGGTCCTGCGTTTTAGAGCTAGAAATAGCAAGTTAAAATAAGGCTAGTCCGTTATCAACTTGAAAAAGTGG | TGG |
| -119N | Fn-dCas12a | *sfgfp* | GTCTAAGAACTTTAAATAATTTCTACTGTTGTAGATGTATCATGTCAATTGGTAAC | TTTC |
| +112N | Fn-dCas12a | *sfgfp* | GTCTAAGAACTTTAAATAATTTCTACTGTTGTAGATTAGCATCACCTTCACCCTCT | TTTG |
| +373N | Fn-dCas12a | *sfgfp* | GTCTAAGAACTTTAAATAATTTCTACTGTTGTAGATACTCGATACGATTAACAAGG | TTTA |
| +710N | Fn-dCas12a | *sfgfp* | GTCTAAGAACTTTAAATAATTTCTACTGTTGTAGATTAGAGCTCATCCATGCCATG | TTTG |
| +131N | Fn-dCas12a | *sfgfp* | GTCTAAGAACTTTAAATAATTTCTACTGTTGTAGATAGGGTGAGTTTTCCGTTTGT | TTTA |
| -175T | Fn-dCas12a | *sfgfp* | GTCTAAGAACTTTAAATAATTTCTACTGTTGTAGATGGTCCGCTTCGACGTACGGT | TTTC |
| +20T | Fn-dCas12a | *sfgfp* | GTCTAAGAACTTTAAATAATTTCTACTGTTGTAGATACTGGAGTTGTCCCAATTCT | TTTC |
| +339T | Fn-dCas12a | *sfgfp* | GTCTAAGAACTTTAAATAATTTCTACTGTTGTAGATAAGGTGATACCCTTGTTAAT | TTTG |
| +666T | Fn-dCas12a | *sfgfp* | GTCTAAGAACTTTAAATAATTTCTACTGTTGTAGATTAACTGCTGCTGGGATTACA | TTTG |
| +139T | Fn-dCas12a | *sfgfp* | GTCTAAGAACTTTAAATAATTTCTACTGTTGTAGATCACTACTGGAAAACTACCTG | TTTG |

| | | | | |
|---|---|---|---|---|
| -119N | Lb-dCas12a | *sfgfp* | GTTTCAAAGATTAAATAATTTCTACTAAGTGTAGATGTATCATGTCAATTGGTAAC | TTTC |
| +112N | Lb-dCas12a | *sfgfp* | GTTTCAAAGATTAAATAATTTCTACTAAGTGTAGATTAGCATCACCTTCACCCTCT | TTTG |
| +373N | Lb-dCas12a | *sfgfp* | GTTTCAAAGATTAAATAATTTCTACTAAGTGTAGATACTCGATACGATTAACAAGG | TTTA |
| +710N | Lb-dCas12a | *sfgfp* | GTTTCAAAGATTAAATAATTTCTACTAAGTGTAGATTAGAGCTCATCCATGCCATG | TTTG |
| +131N | Lb-dCas12a | *sfgfp* | GTTTCAAAGATTAAATAATTTCTACTAAGTGTAGATAGGGTGAGTTTTCCGTTTGT | TTTA |
| -175T | Lb-dCas12a | *sfgfp* | GTTTCAAAGATTAAATAATTTCTACTAAGTGTAGATGGTCCGCTTCGACGTACGGT | TTTC |
| +20T | Lb-dCas12a | *sfgfp* | GTTTCAAAGATTAAATAATTTCTACTAAGTGTAGATACTGGAGTTGTCCCAATTCT | TTTC |
| +339T | Lb-dCas12a | *sfgfp* | GTTTCAAAGATTAAATAATTTCTACTAAGTGTAGATAAGGTGATACCCTTGTTAAT | TTTG |
| +666T | Lb-dCas12a | *sfgfp* | GTTTCAAAGATTAAATAATTTCTACTAAGTGTAGATTAACTGCTGCTGGGATTACA | TTTG |
| +139T | Lb-dCas12a | *sfgfp* | GTTTCAAAGATTAAATAATTTCTACTAAGTGTAGATCACTACTGGAAAACTACCTG | TTTG |
| +39N | Sp-dCas9 | *sfgfp* | CCATCTAATTCAACAAGAATGTTTTAGAGCTAGAAATAGCAAGTTAAAATAAGGCTAGTCCGTTATCAACTTGAAAAAGTGG | TGG |
| +267N | Sp-dCas9 | *sfgfp* | CGTTCCTGTACATAACCTTCGTTTTAGAGCTAGAAATAGCAAGTTAAAATAAGGCTAGTCCGTTATCAACTTGAAAAAGTGG | AGG |
| +20T, +339T | Lb-dCas12a | *sfgfp* | GTTTCAAAGATTAAATAATTTCTACTAAGTGTAGATACTGGAGTTGTCCCAATTCTGTTTCAAAGATTAAATAATTTCTACTAAGTGTAGATAAGGTGATACCCTTGTTAAT | TTTC, TTTG |
| +20T, +666T | Lb-dCas12a | *sfgfp* | GTTTCAAAGATTAAATAATTTCTACTAAGTGTAGATACTGGAGTTGTCCCAATTCTGTTTCAAAGATTAAATAATTTCTACTAAGTGTAGATTAACTGCTGCTGGGATTACA | TTTC, TTTG |
| +20T, +139T | Lb-dCas12a | *sfgfp* | GTTTCAAAGATTAAATAATTTCTACTAAGTGTAGATACTGGAGTTGTCCCAATTCTGTTTCAAAGATTAAATAATTTCTACTAAGTGTAGATCACTACTGGAAAACTACCTG | TTTC, TTTG |
| +339T, +20T | Lb-dCas12a | *sfgfp* | GTTTCAAAGATTAAATAATTTCTACTAAGTGTAGATAAGGTGATACCCTTGTTAATGTTTCAAAGATTAAATAATTTCTACTAAGTGTAGATACTGGAGTTGTCCCAATTCT | TTTG, TTTC |
| +339T, +666T | Lb-dCas12a | *sfgfp* | GTTTCAAAGATTAAATAATTTCTACTAAGTGTAGATAAGGTGATACCCTTGTTAATGTTTCAAAGATTAAATAATTTCTACTAAGTGTAGATTAACTGCTGCTGGGATTACA | TTTG, TTTG |
| +339T, +139T | Lb-dCas12a | *sfgfp* | GTTTCAAAGATTAAATAATTTCTACTAAGTGTAGATAAGGTGATACCCTTGTTAATGTTTCAAAGATTAAATAATTTCTACTAAGTGTAGATCACTACTGGAAAACTACCTG | TTTG, TTTG |
| +666T, +20T | Lb-dCas12a | *sfgfp* | GTTTCAAAGATTAAATAATTTCTACTAAGTGTAGATTAACTGCTGCTGGGATTACAGTTTCAAAGATTAAATAATTTCTACTAAGTGTAGATACTGGAGTTGTCCCAATTCT | TTTG, TTTC |

| | | | | |
|---|---|---|---|---|
| +666T, +339T | Lb-dCas12a | *sfgfp* | GTTTCAAAGATTAAATAATTTCTACTAAGTGTAGATTAACTGCTGCTGGGATTACAGTTTCAAAGATTAAATAATTTCTACTAAGTGTAGATAAGGTGATACCCTTGTTAAT | TTTG, TTTG |
| +666T, +139T | Lb-dCas12a | *sfgfp* | GTTTCAAAGATTAAATAATTTCTACTAAGTGTAGATTAACTGCTGCTGGGATTACAGTTTCAAAGATTAAATAATTTCTACTAAGTGTAGATCACTACTGGAAAACTACCTG | TTTG, TTTG |
| +139T, +20T | Lb-dCas12a | *sfgfp* | GTTTCAAAGATTAAATAATTTCTACTAAGTGTAGATCACTACTGGAAAACTACCTGGTTTCAAAGATTAAATAATTTCTACTAAGTGTAGATACTGGAGTTGTCCCAATTCT | TTTG, TTTC |
| +139T, +339T | Lb-dCas12a | *sfgfp* | GTTTCAAAGATTAAATAATTTCTACTAAGTGTAGATCACTACTGGAAAACTACCTGGTTTCAAAGATTAAATAATTTCTACTAAGTGTAGATAAGGTGATACCCTTGTTAAT | TTTG, TTTG |
| +139T, +666T | Lb-dCas12a | *sfgfp* | GTTTCAAAGATTAAATAATTTCTACTAAGTGTAGATCACTACTGGAAAACTACCTGGTTTCAAAGATTAAATAATTTCTACTAAGTGTAGATTAACTGCTGCTGGGATTACA | TTTG, TTTG |

[a] Identical spacer sequence was used for the Fn-dCas12a and Lb-dCas12a gRNA with the same name

[b] The DNA sequence for the *eyfp* target gene used to design gRNA is:
gataagtccctaactttgacaggtcctaaaggatagtctataatgctagcctgaagctgtcaccggatgtgctttccggtctgatgagtccgtgaggacgaaacagcctctacaaataattttgtttaatactagagaaagaggggaaatactagatggtgagcaagggcgaggagctgttcaccggggtggtgcccatcctggtcgagctggacggcgacgtaaacggccacaagttcagcgtgtccggcgagggcgagggcgatgccacctacggcaagctgaccctgaagttcatctgcaccacaggcaagctgcccgtgccctggcccaccctcgtgaccaccttcggctacggcctgcaatgcttcgcccgctaccccgaccacatgaagctgcacgacttcttcaagtccgccatgcccgaaggctacgtccaggagcgcaccatcttcttcaaggacgacggcaactacaagacccgcgccgaggtgaagttcgagggcgacaccctggtgaaccgcatcgagctgaagggcatcgacttcaaggaggacggcaacatcctggggcacaagctggagtacaactacaacagccacaacgtctatatcatggccgacaagcagaagaacggcatcaaggtgaacttcaagatccgccacaacatcgaggacggcagcgtgcagctcgccgaccactaccagcagaacacccccatcggcgacggccccgtgctgctgcccgacaaccactacctgagctaccagtccgccctgagcaaagaccccaacgagaagcgcgatcacatggtcctgctggagttcgtgaccgccgccgggatcactctcggcatggacgagctgtacaagtaa.

[c] The DNA sequence for the *sfgfp* target gene used to design gRNA is:
tttcggtccgcttcgacgtacggtggaatctgattcgttaccaattgacatgatacgaaacgtaccgtatcgttaaggtctgaagctgtcaccggatgtgctttccggtctgatgagtccgtgaggacgaaacagcctctacaaataattttgtttaactctcgaaggcccctgacacaatgagcaaaggagaagaacttttcactggagttgtcccaattcttgttgaattagatggtgatgttaatgggcacaaattttctgtccgtggagagggtgaaggtgatgctacaaacggaaaactcacccttaaatttatttgcactactggaaaactacctgttccgtggccaacacttgtcactactctgacctatggtgttcaatgcttttcccgttatccggatcacatgaaacgtcatgactttttcaagagtgccatgcctgaaggttatgtacaggaacgcactatatctttcaaagatgacgggacctacaagacgcgtgctgaagtcaagtttgaaggtgatacccttgttaatcgtatcgagttaaagggtattgattttaaagaagatggaaacattcttggacacaaactcgagtacaactttaactcacacaatgtatacatcacggcagacaaacaaaagaatggaatcaaagctaacttcaaaattcgccacaacgttgaagatggttccgttcaactagcagaccattatcaacaaaatactccaattggcgatggccctgtccttttaccagacaaccattacctgtcgacacaatctgtcctttcgaaagatcctaacgaaaagcgtgaccacatggtccttcttgagtttgtaactgctgctgggattacacatggcatggatgagctctacaaataa.

[d] gRNA DNA sequence is colored black for the variable spacer sequence and blue for the appropriate constant sequence of the CRISPRi system (sgRNA chimeric scaffold for Sp-dCas9 or direct repeat for the dCas12a variants).

**Table S8. Plasmids used in this work.**

| Plasmid name | Description | Reference |
|---|---|---|
| pAN1717 | Plasmid backbone (p15a stringent replicon, *kanR*, and sensors *lacI* and *tetR*) for the destination vectors for each CRISPRi system, and plasmid used as a positive control for flow cytometry for conversion of arbitrary units to relative promoter units (RPU). Contains a standard *eyfp* cassette for this conversion. | [2,3] |
| pAN-PTet-dCas9 | Source of the Sp-dCas9 gene ($P_{Tet}$, Sp-dCas9 CDS, $T_{rrnB}$ terminator) for the Sp-dCas9 destination vector; Addgene #62244 | [13] |
| pPPhlF-YFP | Source of $P_{PhlF}$ for construction of the *sfgfp* output plasmid | [8] |
| pAN871 | Source of the LacZα gene and coding sequence for the CRISPRi destination vectors | [8] |
| pCM29 | Source of *sfgfp* coding sequence for the *sfgfp* output plasmid | [19] |
| PTac-YFP | Source of *eyfp* gene (from $P_{Tac}$ to L3S2P21 terminator) for characterization of $P_{Tac}$ on the CRISPRi backbone | This study |
| pSR2000 | Sp-dCas9 destination vector; pAN1717 backbone with *eyfp* gene replaced with Sp-dCas9 gene from pAN-PTet-dCas9, L3S2P11 terminator, and LacZα gene from pAN871, with BsaI recognition sites surrounding the LacZα gene for insertion of a gRNA gene (including PTac and DT11 terminator) | This study |
| pSR2001 | Sp-dCas9 destination vector, version 2. pAN1717 backbone with *eyfp* gene replaced with Sp-dCas9 gene from pAN-PTet-dCas9, L3S2P11 terminator, and a DNA fragment containing $P_{Tac}$, LacZα the chimeric sgRNA scaffold, and DT11 terminator. BsaI recognition sites surround LacZα for insertion of gRNA spacer sequences | This study |
| pSR2002 | Plasmid synthesized by TWIST Biosciences containing the Fn-dCas12a coding sequence. Source of Fn-dCas12a coding sequence for the Fn-dCas12a CRISPRi destination vector | This study |
| pSR2003 | pSR2000 with Sp-dCas9 coding sequence replaced by the Fn-dCas12a coding sequence from pSR2002 | This study |
| pSR2004 | Fn-dCas12a destination vector. pSR2003 with LacZα replaced by a DNA fragment containing $P_{Tac}$, Fn-dCas12a direct repeat sequence, LacZα, spacer DNA, and L3S2P21 terminator with BbsI recognition sites surrounding the LacZα gene for insertion of gRNA spacer sequences | This study |
| pSR2005 | Plasmid synthesized by TWIST Biosciences containing the Lb-dCas12a coding sequence. Source of Lb-dCas12a coding sequence for the Lb-dCas12a CRISPRi destination vector | This study |
| pSR2006 | pAN-PTet-dCas9 with the Sp-dCas9 coding sequence replaced by the Lb-dCas12a coding sequence from pSR2005 | This study |
| pSR2007 | pAN1717 backbone with *eyfp* gene replaced with Lb-dCas12a gene from pSR2006, L3S1P13 terminator, and LacZα gene from pAN871 | This study |
| pSR2008 | Lb-dCas12a destination vector. pSR2007 with LacZα replaced by a DNA fragment containing $P_{Tac}$, Lb-dCas12a direct repeat sequence, LacZα, spacer DNA, and L3S2P21 terminator with BbsI recognition sites surrounding the LacZα gene for insertion of gRNA spacer sequences | This study |
| pP1_output | Output plasmid containing the *eyfp* target gene for characterization of Sp-dCas9. *eyfp* is expressed from the P1 promoter with the RiboJ insulator, B0064 RBS, and L3S2P21 terminator | [21] |
| pSR2009 | Output plasmid containing the *sfgfp* target gene for characterization of Fn-dCas12a and Lb-dCas12a. *sfgfp* is expressed from $P_{PhlF}$ with the RiboJ insulator, 2.5k RBS, and L3S2P21 terminator | This study |

| | | |
|---|---|---|
| pSR2010 | Plasmid to measure response curve of $P_{Tet}$ on the CRISPRi destination backbone. pSR2017 with the Sp-dCas9 coding sequence replaced by the *eyfp* gene (from the RiboJ insulator to shortly after the stop codon) from pAN1717 | This study |
| pSR2011 | Plasmid to measure response curve of $P_{Tac}$ on the CRISPRi destination backbone. pSR2000 with the LacZα gene replaced by the *eyfp* gene (from $P_{Tac}$ to L3S2P21 terminator) from PTac-YFP | This study |
| pSR2012 | pSR2000 with LacZα replaced by $P_{Tac}$, gRNA -123N, and DT11 terminator | This study |
| pSR2013 | pSR2000 with LacZα replaced by $P_{Tac}$, gRNA -89N, and DT11 terminator | This study |
| pSR2014 | pSR2000 with LacZα replaced by $P_{Tac}$, gRNA +34N, and DT11 terminator | This study |
| pSR2015 | pSR2000 with LacZα replaced by $P_{Tac}$, gRNA +334N, and DT11 terminator | This study |
| pSR2016 | pSR2000 with LacZα replaced by $P_{Tac}$, gRNA +664N, and DT11 terminator | This study |
| pSR2017 | pSR2000 with LacZα replaced by $P_{Tac}$, gRNA -144T, and DT11 terminator | This study |
| pSR2018 | pSR2000 with LacZα replaced by $P_{Tac}$, gRNA -128T, and DT11 terminator | This study |
| pSR2019 | pSR2000 with LacZα replaced by $P_{Tac}$, gRNA +28T, and DT11 terminator | This study |
| pSR2020 | pSR2000 with LacZα replaced by $P_{Tac}$, gRNA +325T, and DT11 terminator | This study |
| pSR2021 | pSR2000 with LacZα replaced by $P_{Tac}$, gRNA +659T, and DT11 terminator | This study |
| pSR2022 | pSR2004 with LacZα replaced by gRNA -119N | This study |
| pSR2023 | pSR2004 with LacZα replaced by gRNA +112N | This study |
| pSR2024 | pSR2004 with LacZα replaced by gRNA +373N | This study |
| pSR2025 | pSR2004 with LacZα replaced by gRNA +710N | This study |
| pSR2026 | pSR2004 with LacZα replaced by gRNA +131N | This study |
| pSR2027 | pSR2004 with LacZα replaced by gRNA -175T | This study |
| pSR2028 | pSR2004 with LacZα replaced by gRNA +20T | This study |
| pSR2029 | pSR2004 with LacZα replaced by gRNA +339T | This study |
| pSR2030 | pSR2004 with LacZα replaced by gRNA +666T | This study |
| pSR2031 | pSR2004 with LacZα replaced by gRNA +139T | This study |
| pSR2032 | pSR2008 with LacZα replaced by gRNA -119N | This study |
| pSR2033 | pSR2008 with LacZα replaced by gRNA +112N | This study |
| pSR2034 | pSR2008 with LacZα replaced by gRNA +373N | This study |
| pSR2035 | pSR2008 with LacZα replaced by gRNA +710N | This study |
| pSR2036 | pSR2008 with LacZα replaced by gRNA +131N | This study |
| pSR2037 | pSR2008 with LacZα replaced by gRNA -175T | This study |
| pSR2038 | pSR2008 with LacZα replaced by gRNA +20T | This study |
| pSR2039 | pSR2008 with LacZα replaced by gRNA +339T | This study |
| pSR2040 | pSR2008 with LacZα replaced by gRNA +666T | This study |
| pSR2041 | pSR2008 with LacZα replaced by gRNA +139T | This study |
| pSR2042 | pSR2008 with LacZα replaced by gRNA array +20T, +339T | This study |

| | | |
|---|---|---|
| pHB048 | pSR2008 with LacZα replaced by gRNA array +20T, +666T | This study |
| pHB050 | pSR2008 with LacZα replaced by gRNA array +20T, +139T | This study |
| pSR2043 | pSR2008 with LacZα replaced by gRNA array +339T, +20T | This study |
| pSR2044 | pSR2008 with LacZα replaced by gRNA array +339T, +666T | This study |
| pHB054 | pSR2008 with LacZα replaced by gRNA array +339T, +139T | This study |
| pHB049 | pSR2008 with LacZα replaced by gRNA array +666T, +20T | This study |
| pHB053 | pSR2008 with LacZα replaced by gRNA array +666T, +339T | This study |
| pSR2045 | pSR2008 with LacZα replaced by gRNA array +666T, +139T | This study |
| pHB051 | pSR2008 with LacZα replaced by gRNA array +139T, +20T | This study |
| pHB055 | pSR2008 with LacZα replaced by gRNA array +139T, +339T | This study |
| pSR2046 | pSR2008 with LacZα replaced by gRNA array +139T, +666T | This study |
| pSR2047 | pSR2001 with LacZα replaced by gRNA +39N | This study |
| pSR2048 | pSR2001 with LacZα replaced by gRNA +267N | This study |

### Extended Methods

#### Phylogenetic tree analysis

Phylogenetic analysis was performed on 40 strains of *E. coli*, including the four used in this work, using multi-locus sequencing analysis (MLSA) of 7 housekeeping genes (*adk*, *fumC*, *gyrB*, *icd*, *mdh*, *purA*, and *recA*) used in previous work.[22] Strains of *E. coli* were chosen for analysis based on common use for study and genetic diversity (**SI Table 1**). Additionally, the type strain of *Escherichia fergusonii* (FDAARGOS_1499) was included to root the phylogenetic tree. The annotated coding sequences (CDS) of each strain were downloaded from NCBI using the datasets tool (v15.33.0)[23] using the RefSeq Accession number (**SI Table 1**). The DNA sequences of the 7 housekeeping genes from MG1655 were aligned to the annotated CDS file for each strain using BLAST+ (v2.12.0)[24] on a command line and results parsed using a custom Python script to determine the gene sequence for each housekeeping gene in each strain. The housekeeping gene sequences for all strains were concatenated together, then aligned using MAFFT (v7.487)[25] with the 'globalpair' option.

The multiple sequence alignment results were analyzed in R (v4.3.0)[26] using the ape (v5.0)[27] and phangorn (v2.11.1)[28] packages. The tree was constructed using the maximum likelihood (ML) method by choosing the best fitting ML model via the modelTest() and as.pml() functions. The best model was considered the one with the minimum Bayesian information criterion (BIC) value. This model was then optimized for topology using the optim.pml() function. The phylogenetic tree was then created using treeio (v1.24.3),[29,30] tidytree (v0.4.6),[30] and ggtree (v3.8.2)[30,31] with a custom R script. The ecological niche of each strain was determined based on consensus in the literature, and the phylogroup determined using ClermonTyping.[1]

#### DNA Assembly

PCRs were performed using Q5 DNA polymerase (NEB) according to the manufacturer's protocol. DNA oligos were annealed for the assembly of most gRNA designs into the destination vector or the creation of certain parts (e.g. terminators) for some plasmid assemblies. This was performed by first determining the melting and maximum hairpin temperatures for each oligo in a set using the OligoAnalyzer tool (IDT) under the following reaction conditions: 22.4 µM oligo, 6.7 mM $MG^{2+}$, 0 mM $Na^{+}$, and 0 mM dNTPs. To anneal the oligo pairs, 8.5 µL of a 50 µM stock of each oligo was combined with 2 µL of ExoI buffer (NEB) and incubated on a thermocycler under the following conditions: 98 °C for 2 minutes; decreasing by 2 °C with 1 minute at each step until 10 °C above the highest predicted melting temperature; decreasing by 1 °C with 1 minute at each step until reaching the highest predicted hairpin temperature; and decreasing by 2 °C with 1 minute at each step until 50 °C. The reaction mixture was then incubated at 37 °C with 20 U ExoI (NEB) for 1 hour. PCRs and annealed oligos were purified for downstream applications using the GenCatch PCR Purification Kit according to the manufacturer's protocol.

Plasmids were assembled via Type IIS DNA assembly using T4 DNA ligase (NEB) and a Type IIS enzyme (BsaI, BbsI, or SapI) as previously described.[21] Briefly, a reaction mixture was created using 20 fmol of a destination vector (or inverse PCR product of a plasmid backbone), 40 to 200 fmol of each DNA insert (or 0.5 µL of an annealed DNA oligo fragment), 0.5 µL of 10X T4 ligase buffer (NEB), 10 U Type IIS enzyme (NEB), 250 U T4 DNA ligase, and nuclease-free water up to a total reaction volume of 5 µL. To assemble the DNA fragments, the reaction mixture was incubated on a thermocycler for either an isothermal reaction (one DNA insert) for 4-8 hours at 37 °C for or a cycling reaction with alternating steps at 16 °C and 37 °C for 2-5 minutes each for a total of 30-36 cycles. The assembly reaction was then incubated for 30 minutes at 55 °C to digest unassembled DNA fragments and then 20 minutes at 65 °C to deactivate the enzymes. The DNA assembly reaction was transformed into a cloning strain of *E. coli* using chemical transformation and plated onto LB agar plates with the appropriate antibiotic and, if applicable, X-gal and IPTG for blue-white screening. Colonies were screened using colony PCR and gel electrophoresis. A suspected positive colony was then grown overnight at 37 °C and 250 rpm, a frozen glycerol stock made, and the remaining culture miniprepped using the QIAprep Spin Miniprep kit (Qiagen) according to the manufacturer's protocol. The purified plasmid was sent for DNA sequencing (Azenta or Plasmidsaurus) to verify the construct before characterization.

Plasmids used to characterize the single gRNA of each CRISPRi system were constructed via Type IIS DNA assembly with either a DNA fragment containing $P_{Tac}$, the gRNA sequence, and terminator (Sp-dCas9) or annealed DNA oligos containing the 20 bp spacer sequence (Fn-dCas12a and Lb-dCas12a). Both were flanked by the appropriate Type II recognition sites (BsaI for Sp-dCas9 and BbsI for the dCas12a systems) and 4 nt linker sequences for insertion into the destination vector. For the dual gRNA designs for Lb-dCas12a, the procedure for construction was the same except the gRNA arrays were composed of two DNA fragments made of annealed oligos. These fragments were designed to contain part of the Lb-dCas12a direct repeat at the 5′ of the first gRNA and the remaining part at the 3′ end of the second gRNA, with an orthogonal 4 nt linker sequence chosen within the direct repeat for seamless assembly.

### Chemical transformation of *E. coli*

Chemical transformation was used to introduce cloned plasmid DNA into chemically competent *E. coli* cells of either NEB 5-alpha or NEB 10-beta. Aliquots of 5 µL of cells were thawed on ice, to which 1-2 µL of a DNA assembly reaction or a minimum of 100 ng of purified DNA was added. The cell-DNA mixture was incubated on ice for 15 to 30 minutes, followed by a heat shock at 42 °C for 30 seconds. Subsequently, 100 µL of SOC recovery medium was added to each aliquot. The cells were then incubated in a shaking incubator at 37 °C and 250 rpm for 1 hour for recovery. The cells were spread on LB agar plates containing the appropriate antibiotics and incubated overnight at 37 °C to obtain single transformants. If needed, 4 µL of 1 M IPTG and 120 µL of 20 mg/mL X-gal (GoldBio) was spread on the plate surface for blue-white screening.

### Construction of promoter characterization plasmids for $P_{Tet}$ and $P_{Tac}$

To characterize the sensor response for $P_{Tet}$ and $P_{Tac}$ in the context of the CRISPRi plasmids, Sp-dCas9 and gRNA on the corresponding CRISPRi plasmid design (except for $P_{Tet}$ for Sp-dCas9) were replaced by an *eyfp* cassette (RiboJ, B0064, *efyp*, and L3S2P21) from the standard pAN1717.[3] The plasmid to characterize the aTc sensor containing $P_{Tet}$ was constructed using Type IIS DNA assembly with two DNA fragments. The plasmid backbone was PCR amplified around the Sp-dCas9 coding sequence on pSR2017 (containing gRNA –144T) and the *eyfp* cassette was amplified from pAN1717.[3] The plasmid to characterize the IPTG sensor was built in the same manner as the Sp-dCas9 destination vector, except the DNA fragment containing LacZα was replaced by the standard *eyfp* gene with $P_{Tac}$ from PTac-YFP.

#### Sensor characterization for inducible promoter inputs

The inducible promoters $P_{Tet}$ and $P_{Tac}$ were characterized in MG1655, EcN, CFT073, and UMN026 using flow cytometry with the same incubation and induction procedure as previously described (**Methods**), except each strain contained only the sensor characterization plasmid (pSR2010 or pSR2011) and a single inducer (aTc for $P_{Tet}$ or IPTG for $P_{Tac}$). Inducers were added to the media and titrated 2-fold for a large range of concentrations. Fluorescence was converted from arbitrary units to relative promoter units (RPU) using the same equation as used to calculate relative expression units (REU) (**Methods**).

#### Determination of sensor response functions

The response functions for the aTc and IPTG sensors were fit to the following Hill equation[32]:

$$y = y_{min} + (y_{max} - y_{min}) \frac{x^n}{K_d^n + x^n}$$

where $y$ is the promoter output in RPU, $x$ is the input inducer concentration, $y_{min}$ is the minimum promoter output in RPU, $y_{max}$ is the maximum promoter output in RPU, $K_d$ is the apparent dissociation constant, and $n$ is the Hill coefficient. The average outputs at each inducer concentration for each strain were $\log_{10}$-transformed and then fit using the least squares method to estimate $y_{min}$, $y_{max}$, $n$, and $K_d$. Fits were performed using the least_squares() function from SciPy (v1.12.0)[33] without bounds or initial values applied to fit the data.

#### Construction of CRISPRi plasmids

Destination vectors were constructed for each CRISPRi system for easy and efficient insertion of arbitrary gRNA sequences via Type IIS DNA assembly using BsaI for Sp-dCas9 and BbsI for Fn-dCas12a and Lb-dCas12a. These vectors have LacZα in place of the gRNA for blue-white screening after DNA assembly and transformation. The plasmid backbone chosen for all destination vectors was pAN1717,[2,3] which contains the p15A origin of replication and the sensor block containing the TetR and LacI regulators for $P_{Tet}$ and $P_{Tac}$, respectively. The RBSs controlling the expression of each dCas12a protein were designed using the RBS calculator from the Salis group[9–12] to be of a similar strength (translation initiation rate of about 3700) as that for Sp-dCas9 on pAN-PTet-dCas9[13] (Addgene #62244).

The Sp-dCas9 destination vector, pSR2000, was constructed using a single Type IIS DNA assembly with SapI. DNA fragments were PCR amplified for: Sp-dCas9 (including $P_{Tet}$ and the $T_{rrnB}$ terminator) from pAN-PTet-dCas9[13] and LacZα from pAN871[8], and the backbone. The L3S3P21 terminator[14] was annealed using two DNA oligos. The Fn-dCas12a destination vector, pSR2004, was constructed using two Type IIS DNA assembly steps. In the first DNA assembly using SapI, the Fn-dCas12a coding sequence was PCR amplified from a commercially synthesized plasmid (Twist Bioscience), and the backbone (including $P_{Tet}$ and $T_{rrnB}$) was amplified from a second version of the Sp-dCas9 destination vector using PCR (Fig. S32H). The subsequent DNA assembly used BsaI to insert a synthesized gene fragment (Twist Bioscience) containing $P_{Tac}$, the Fn-dCas12a direct repeat sequence, LacZα, a spacer DNA sequence, and the L3S2P21 terminator[14] in place of the initial LacZα fragment. The Lb-dCas12a destination vector, pSR2008, was assembled using three Type IIS assembly reactions due to a SapI recognition site in the Lb-dCas12a coding sequence. The first DNA assembly reaction used BbsI to replace the Sp-dCas9 coding sequence on pAN-PTet-dCas9[13] with the Lb-dCas12a coding sequence from a commercially synthesized plasmid (Twist Bioscience). The second DNA assembly used BbsI to place the L3S1P13 terminator between $P_{Tet}$ and the LacZa gene. The final DNA assembly for the Lb-dCas12a destination vector used the same strategy as that for the second assembly for Fn-dCas12a, except incorporating the Lb-dCas12a direct repeat sequence. For the dCas12a destination vectors, an extra spacer DNA sequence was placed between LacZa and the L3S2P21 terminator as the hairpin of a terminator has been shown to affect crRNA processing by Fn-Cas12a.[34]

#### Titration experiments for CRISPRi expression in *E. coli*

Samples were prepared for the CRISPRi repression assays using flow cytometry as previously described (**Methods**) with different concentrations of inducers added to the media. For each CRISPRi system, the level of expression of the dCas protein was titrated with the gRNA either uninduced (0 mM IPTG) or fully induced (1 mM IPTG). Titrations of aTc to vary dCas protein expression were performed as either 4-fold or 2-fold serial dilutions (48.4 µL or 145 µL of media with aTc, respectively, into 145 µL fresh media). Antibiotics and 1 mM IPTG were added to the media prior to dilutions as needed. The gRNAs chosen were –144T for Sp-dCas9 (all strains), +20T for Fn-dCas12a (all strains), and –119N (EcN, CFT073) or +20T (UMN026) for Lb-dCas12a. For the Sp-dCas9 system, additional experiments titrating gRNA expression were similarly performed via 2-fold serial dilutions of IPTG with Sp-dCas9 induced using 0.25 ng/mL aTc (same as in Figure 3). The experiments titrating aTc 4-fold were performed for all gRNA designs in each CRISPRi systems. Higher resolution 2-fold titrations of aTc and IPTG were performed as described above for the gRNA designs that provided the greatest repression of the corresponding target protein (+34N and -144T for Sp-dCas9; -119N and +20T for Fn-dCas12a and Lb-dCas12a).

#### CRISPRi modeling at quasi-steady state conditions

Here, we modified a previously published ODE model for Sp-dCas9 CRISPRi repression[35] that includes resource competition when multiplexing. We incorporated the $P_{Tet}$ and $P_{Tac}$ inducible promoters used for expression of Sp-dCas9 and the gRNA, respectively, into the model. Briefly, the model predicts the concentration of the product of a target gene based on the input production rates of Sp-dCas9 and the gRNA and using the rates of the components forming the CRISPRi complex and then binding to the target DNA. For this work, steady state conditions were assumed such that the change in production rates for all components (Sp-dCas9, gRNA, and the target gene product) were constant over time. The decay rates for Sp-dCas9 and the gRNA and the equilibrium constants for the CRISPRi complex and CRISPRi-DNA complex were assumed to be the same as those in the original model. The concentration of the target DNA site was increased 5-fold to simulate the increased copy number of a gene target on a low copy number plasmid (pSC101, which was used for our output plasmids) compared to the genome.

The inducible promoters were incorporated into the model by defining the Sp-dCas9 and gRNA production rates as the RPU outputs from the fitted Hill equation response function for the $P_{Tet}$ and $P_{Tac}$ inducible promoters, respectively, in MG1655 (**Table S3**). The concentration of the target gene's product was then simulated over a range of input aTc and IPTG inducer concentrations to create a surface plot. Additional curves were created at single concentrations of the inducers to simulate the experimental data obtained from the corresponding flow cytometry assays. All simulations were performed using a custom MATLAB script (R2023b).

The equations used for this model are defined below. The equations used for the output for the $P_{Tet}$ promoter (for Sp-dCas9) and $P_{Tac}$ promoter (for the gRNA) are respectively:

$$\alpha_d = \alpha_{min} + \frac{(\alpha_{max} - \alpha_{min}) \cdot x^n}{K^n + x^n}$$

$$u_i = b_{min} + \frac{(b_{max} - b_{min}) \cdot x^m}{K^m + x^m}$$

Here, $\alpha_d$ is the production rate of Sp-dCas9 (in RPU), $u_i$ is the production rate of the $i^{th}$ gRNA, and $x$ is the input inducer concentration. For Sp-dCas9 expression under the $P_{Tet}$ promoter, the fitted parameter values are $\alpha_{min}$ = 0.002843194, $\alpha_{max}$ = 7.445104004, $K$ = 1.950196571, and $n$ = 3.111927761, and $x$ corresponds to the aTc concentration in ng/mL. For gRNA expression under the $P_{Tac}$ promoter, the parameters are $b_{min}$ = 0.006978277, $b_{max}$ = 5.998225651, $K$ = 0.313434904, and $m$ = 1.658421251, with $x$ representing the IPTG concentration in mM. The conservation of mass for the Sp-dCas9 protein and gRNA are modelled using the equations

$$d_t = \frac{\alpha_d}{\delta}$$

$$\bar{g}_i = \frac{u_i}{\theta}$$

$d_t$ is the total concentration of Sp-dCas9 proteins (both free and in complex with a gRNA), $\delta$ is the decay rate of proteins (including Sp-dCas9), $\bar{g}_i$ is the concentration of the $i^{th}$ gRNA at steady state, and $\theta$ is the decay rate of the gRNA. Assuming quasi-steady state conditions and substituting the above conservation of mass equations, the following kinetic equation was derived to model the steady state output of target protein with varying inducer inputs:

$$d\left(1 + \sum_{i=1}^{n} \frac{g_i}{K_i}\right) + \sum_{i=1}^{n} \frac{d * g_i * D_{it}}{K_i Q_i + d * g_i} - d_t = 0,$$

where $d$ is the concentration of free Sp-dCas9 proteins, $K_i$ is the dissociation (equilibrium) constant for the $i^{th}$ Sp-dCas9-gRNA complex, $D_{it}$ is the total concentration of all target sites for the $i^{th}$ gRNA (both bound and unbound), and $Q_i$ is the dissociation constant for the $i^{th}$ Sp-dCas9-gRNA-DNA complex. Under steady state conditions, the output concentration for the protein of the target gene can be modeled using

$$\bar{Y}_i = \frac{\kappa_i D_{it}/\delta}{1 + \frac{d * \bar{g}_i}{K_i Q_i}}$$

where $\bar{Y}_i$ is the concentration of the $i^{th}$ target protein at steady state and $\kappa_i$ is the production rate of the $i^{th}$ target protein. The constants chosen for this work are the same as that in the original publication,[35] except $D_{it}$, which was 5 times greater than the original model to describe the difference in copy number in our system. The constants were defined as follows: $\theta$ = 100 hr$^{-1}$, $\delta$ = 1 hr$^{-1}$, $K_i$ = 0.01 nM, $Q_i$ = 0.5 nM, $\kappa_i$ = 1000 hr$^{-1}$, and $D_{it}$ = 10 nM.

For the purposes of this work, only a single target gene was considered, although these equations could be used to simulation the repression of multiple target genes at once. Repression for these simulations was calculated as the ratio of the target protein concentration at steady state ($\overline{Y}_i$) without inducers present (0 ng/mL aTc and 0 mM IPTG) and that with at least one inducer present, analogous to what was done for the flow cytometry experiments.

**Design of Experiments (DOE) to assess effects of CRISPRi component expression**

A custom DOE was created and analyzed using JMP Pro v17.0. The expression of the dCas protein and gRNA were designated as continuous variables with 5 levels each. The day that the experiment was performed was input as a blocking nominal variable. The measured response variable was the fold repression, defined previously for the flow cytometry assays (**Methods**). To allow coverage of the entire design space with a few replicates, a total of 30 measurements were taken for each CRISPRi system. A polynomial model to analyze the output was designed to include all primary factors and secondary interactions for the gRNA and dCas protein expression levels, with the goal of maximizing the repression.

The levels of expression for each CRISPRi component were determined using the output RPU values from the component's respective inducible promoter. To allow estimation of curvature within the design space, the levels were chosen using a Box-Wilson central composite inscribed (CCI) design, using the designated minimum and maximum points in the range of expression as the star points. Thus, in terms of the normalized distance from the central point in the design space, the levels were chosen as -1, -0.707, 0, 0.707, and 1 within the range of expression. RPU values were $\log_{10}$-transformed prior to creation of the DOE matrix in JMP Pro. No growth toxicity or bimodality in fluorescence was observed with changing expression levels of the gRNA from the Sp-dCas9 titration experiments. Therefore, the range of expression for the gRNA was chosen as the entire range of output for the $P_{Tac}$ promoter. In contrast, growth toxicity was observed at high expression of the dCas protein (≥ 2 ng/mL aTc; **Figures 4, 6 and 8**). Thus, the span of expression levels for the dCas protein was limited to a maximum of 0.830 RPU (1 ng/mL aTc) as output. The specific RPU expression values and corresponding aTc concentrations for $P_{Tet}$ are the following: 0.00781 ng/mL for 0.00284 RPU, 0.169 ng/mL for 0.00653 RPU, 0.380 ng/mL for 0.0486 RPU, 0.748 ng/mL for 0.362 RPU, and 1 ng/mL for 0.830 RPU. The RPU values and IPTG concentrations for $P_{Tac}$ are: 0.00100 mM for 0.00741 RPU, 0.00755 mM for 0.0194 RPU, 0.0.399 mM for 0.197 RPU, 0.206 mM for 2.00 RPU, and 1.00 mM for 5.23 RPU.

The DOE matrix was assayed using flow cytometry as previously described (**Methods**). MG1655 strains containing gRNA designs that showed the greatest repression for each CRISPRi system (–119N for Fn-dCas12a and Lb-dCas12a, –144T for Sp-dCas9) were chosen for the greatest range of values for modeling. The results were analyzed by running the model fit in JMP Pro for each CRISPRi system using the standard least squares method and an emphasis on effect screening. Variables that showed insignificant effects in the output (defined as *p*-value > 0.01) were removed, starting from the variable with the lowest significance, until the remaining variables were all statistically significant. The resulting significant variables are shown in **Figure S27,** and specific values for the full and reduced models are provided in **Tables S5** and **S6**, respectively.

#### Epistatic models for Lb-dCas12a dual gRNA designs

Linear fits were performed using the lm() function to determine lines of best fit and goodness of fits for the investigation of the epistatic effects for fold repression of the dual gRNA designs relative to their corresponding single gRNA. Both the multiplicative and additive models were examined. The additive model predicted the average and error of the fold repression for the dual gRNA design as the sum of the average and standard deviation, respectively, of each single gRNA component. Linear fits with a y-intercept of 0 (y = m*x) were then performed between the average predicted and measured repression values for all strains combined and each strain individually. In the multiplicative model, the predicted fold repression was calculated as the product of the repression of each gRNA comprising the array. Error in this prediction was calculated using error propagation of the means and standard deviations of the repression of each gRNA in the array as: $\sqrt{\left(\frac{x_1}{\sigma_1}\right)^2 * \left(\frac{x_2}{\sigma_2}\right)^2}$.

### Supplemental Notes

#### Supplemental Note 1. Tuning CRISPRi repression in MG1655

Titrations of the dCas and gRNA can be performed to determine induction levels needed for maximum repression. We performed these experiments for each CRISPRi system in *E. coli* MG1655. For this demonstration, the expression of the dCas protein was titrated with either the gRNA induced or uninduced and the resulting fluorescence measured using flow cytometry (**SI Extended Methods**). In practice, the gRNA expression can also be titrated or combinatorial optimization methods can be applied to determine induction for both gRNA and dCas simultaneously (e.g. using design of experiments, **Supplementary Note 3**).

Fluorescent measurements and fold repression from the gRNA designs for all CRISPRi systems in MG1655 showed similar trends between the systems, with nuanced differences between systems and some gRNA. As expected, gRNA having little to no repression from previous experiments demonstrated similar fluorescence and repression across all aTc concentrations for each CRISPRi system (**Figures S22-S24**, **S30**), except –128T with Sp-dCas9 exhibited slightly greater repression at higher aTc concentrations (**Figure S22**). With induction of the gRNA, all single gRNA that showed repression from the prior experiments exhibited a maximum fold repression at 0.5 ng/mL aTc and a notable decrease in repression with more aTc (2 ng/mL aTc) (**Figures S22-S24**). In contrast, most dual gRNA designs (7/12) for Lb-dCas12a showed maximum repression at 2 ng/mL aTc (**Figure S30**). Active gRNA showed repression without dCas induction, demonstrating leaky expression of dCas led to some repression even when using the very tightly regulated $P_{Tet}$. This highlights the importance of strong transcriptional regulation for both the dCas protein and gRNA to limit unwanted, leaky repression of a target gene, which can occur when only one component is regulated.[36–39] Slight repression without gRNA induction for active gRNA designs was observed at the highest concentrations of aTc (0.5 and 2 ng/mL) for Lb-dCas12a and Fn-dCas12a (**Figures S23** and **S24**), but not Sp-dCas9, coinciding with the increased repression seen from the dCas12a systems.

Higher resolution titration experiments of the CRISPRi components for select gRNA of each CRISPRi system were performed to more precisely elucidate the induction levels for their maximal repression. These experiments were performed for the two gRNA that yielded the greatest repression for each CRISPRi system (**SI Extended Methods**). Two-fold titrations of aTc to vary expression of dCas protein at maximum induction of the gRNA (1 mM IPTG) showed similar patterns in repression and fluorescence (**Figure S25**) as observed from the lower resolution experiments for these gRNA (**Figures S22-S24**), except with variable induction levels for maximum repression ranging from 0.25 – 2 ng/mL aTc. This appeared to depend on both the CRISPRi system and gRNA design. Both gRNA designs for Fn-dCas12a showed the greatest average repression at 0.25 ng/mL aTc but at either 0.5 or 1 ng/mL aTc for Lb-dCas12a and 0.5 or 2 ng/mL aTc for Sp-dCas9, depending on the gRNA. Unlike for dCas, increasing expression of the gRNA at an intermediate expression of Sp-dCas9 (0.25 ng/mL aTc) demonstrated monotonically increasing repression and decreasing fluorescence output (**Figure S25B**). These results reveal similar but nuanced patterns in CRISPRi repression in MG1655 by the three tested CRISPRi systems at different expression levels of each CRISPRi component.

**Supplemental Note 2. Modeling of CRISPRi target gene repression**

An ODE model was created to describe CRISPRi repression of a target gene using Sp-dCas9, which was based on a previously published model for Sp-dCas9 repression using multiple gRNA,[35] and modified here to incorporate the $P_{Tet}$ and $P_{Tac}$ induction of dCas and gRNA used in our genetic design (**SI Extended Methods**). Similar to the experiments measuring repression using flow cytometry, fold repression through modeling simulations was calculated as the ratio of the output target protein concentration without and with the inducers present. Simulations were performed over a series of aTc and IPTG concentrations to model the resulting target protein concentration and fold repression.

Model simulations generally followed trends observed in experimental measurements of Sp-dCas9 repression in MG1655 with changing concentrations of IPTG and aTc, except at high concentrations of aTc. The model demonstrated a sigmoidal increase in fold repression with increasing concentrations of each inducer (**Figure S26A**), corresponding to an inverse sigmoidal decrease in target protein concentration (**Figure S26B**). This general trend agrees with the experimental measurements for Sp-dCas9 repression with increasing aTc concentrations, except for aTc concentrations above 0.5 ng/mL for most gRNA designs (**Figures S22** and **S25A**). This corresponds to the Sp-dCas9 expression levels that were shown to cause significant growth toxicity and a decrease in growth rate (**Figure 4**). In contrast, the model matches the experimental observations with increasing IPTG and an intermediate concentration of aTc (0.25 ng/mL; **Figure S25B**). Interestingly, the model captured leaky repression of a target gene with high concentration of a single inducer like in the experimental data (**Figure S22**), but only for aTc (i.e., dCas protein expression) and not IPTG (i.e., gRNA expression), which was observed more often in the experimental data for some CRISPRi systems (**Figures S22-S24**). This difference may be due to the constant parameter values chosen for the model, such as the decay rate of the gRNA. Overall, these results demonstrate that a basic, deterministic model for CRISPRi repression may be used to simulate the general trends observed from experimental data at induction levels of a dCas protein that do not cause significant decreases in growth rate.

**Supplemental Note 3. Design of experiments analysis and modeling**

We applied design of experiments (DOE) to statistically determine the effects of the expression of the dCas protein and gRNA on repression of the target gene for each CRISPRi system. For the DOE experimental matrix, the expression levels for the dCas and gRNA were determined based on a Box-Wilson central composite inscribed (CCI) design, using a range of outputs from each inducible promoter (in RPU) in MG1655 as inputs for the design (**SI Extended Methods**). This DOE model allows estimation of curvature in the design space for each factor. The range of expression for the dCas protein was limited to below the maximum expression at 2 ng/mL aTc due to growth toxicity observed (**Figures 4, 6,** and **8**). Samples were prepared and repression measured using the same flow cytometry assay as previously described (**SI Extended Methods**). The results were fit to a polynomial model that included second-order (pairwise) interactions between factors with the objective of maximizing repression.

The reduced fit models suggest that gRNA and dCas expression affect the observed repression differently for each CRISPRi system, with the dCas12a variants showing more similar results. The reduced model for Sp-dCas9 showed that only gRNA expression had a significant effect (first-order main effect) on repression of the target gene under these DOE conditions (**Figure S27A**, **Table S6**). In contrast, results for Fn-dCas12a exhibited significant main effects for both gRNA and Fn-dCas12a expression on the measured repression of the target gene, in addition to second-order interactions between the induction of both and a second-order dependence on Fn-dCas12a (**Figure S27B**, **Table S6**). Lb-dCas12a also yielded significant main effects for gRNA and Lb-dCas12a expression and only one significant second-order interaction between the gRNA and Lb-dCas12a expression (**Figure S27C**, **Table S6**). These models result in significant curvature in the repression response for the design space for Fn-dCas12a and Lb-dCas12a CRISPRi but not for Sp-dCas9 CRISPRi in MG1655. This DOE optimization approach and resulting models can reveal unique factors affecting repression for each CRISPRi system for a given application.

**Supplemental Note 4. Design of gRNAs using custom Python scripts**

The set of Python scripts developed for the three CRISPRi systems assayed in this work (adapted from Wang, et al.[40]) can be used to design gRNA for any number of coding sequences, including genome-wide libraries for any organism with a sequenced and annotated genome. The input files for the set of scripts include a configuration file containing input parameters for individual gRNA design and (if applicable) genome-wide library design, a FASTA file of the target sequence(s), and a FASTA file of the host organism's annotated genome (including multiple chromosomes and plasmids). If a genome-wide library is being designed and gRNA should be designed for genes with multiple copies in the genome, a BLAST results file of all target genes aligned against all target genes can be generated. Design criteria for gRNA design include the CRISPRi system (Sp-dCas9, Fn-dCas12a, or Lb-dCas12a), GC content, limited off-target effects, specified DNA sequence(s) to avoid (e.g. enzyme recognition sequences). Additionally, for *E. coli* and the Sp-dCas9 system, the scripts can apply the the "bad seed" effect[4] for gRNA design. Additional general input parameters include the allowable proportion of the coding sequence where gRNA designs can target, maximum number of gRNA designs per gene, strand of DNA to target (with option to target both DNA strands), and number of non-targeting negative control (NC) gRNA to design (used for genome-wide library design). Constant upstream and downstream sequences can be appended to the designed spacer sequences to create oligo sequences.

Using these input parameters and files, the scripts output the designed gRNA library in both CSV and FASTA format and additional files describing details and statistics of the gRNA library. This includes the genes clustered based on homology from BLAST alignment (if provided), gene statistics (length of and number of gRNA targeting each gene), gRNA statistics (relative targeting position along the target sequence and GC content), and gRNA spacer sequences with the target PAM sequence. Histograms of the gRNA designs per gene and relative targeting position along the target coding sequence are created.

For a target gene, the script will find all available spacer sequences on the specified strand of DNA based on the canonical PAM sequence and with the entire spacer sequence contained within the target gene sequence. The script screens putative spacer sequences for appropriate GC content and any designated disallowed DNA sequences. Additionally for *E. coli* using Sp-dCas9, putative spacer sequences can be screened to remove those with any of the ten "bad seeds" identified in previous work.[4] Spacer sequences that pass these design criteria are then aligned to the host's genome using SeqMap[41] to predict off-target effects using a penalty scoring system. The locations of mismatches between the spacer sequence and potential off-target sites using canonical and non-canonical PAM sequences are used to predict off-target effects. Candidate spacer sequences are acceptable if they are within the input penalty threshold, and they are then chosen by default using the target location, starting with those closest to the start codon if the acceptable candidate sequences exceed the input number of gRNA designs per gene. If user-specified and more gRNA designs are needed, additional gRNA targeting the beginning (within the first 100 bp) of the unpreferred strand can be designed, as a prior study demonstrated effective repression targeting this region of the template strand using Sp-dCas9.[4]

The design parameters for each CRISPRi system vary slightly to accommodate the different PAM sequences, gRNA structure, and propensity for off-target effects of the CRISPR system. For Sp-dCas9, these parameters were kept the same as those in the original script set[40] with additional code added to incorporate more optional design criteria (unallowed DNA sequences and the "bad seed" effect[4]). The canonical NGG PAM was used to search for spacer sequences and off-target effects, and the non-canonical NAG PAM was only used to search for potential off-target effects. The regions in the gRNA spacer sequence for assigning penalty values were defined from 3′ to 5′ (i.e. starting adjacent to the PAM) as follows: seed region = nucleotides 1 – 7, middle region = nucleotides 8 – 12, distal region = nucleotides 13 – 20. These regions were chosen based on a prior study.[42] For Fn-dCas12a, the set of scripts were adjusted to use the canonical TTTV PAM sequence and non-canonical GTTV, CTTV, ATTV, and TCTV PAM sequences.[43,44] The regions of the gRNA were redefined for the dCas12a system based on previous studies[44,45] and are as follows (5′ to 3′): seed region = nucleotides 1 – 6, middle region = nucleotides 7 – 15, distal region = nucleotides 16 – 20. Penalty values for each region of the gRNA were increased to account for the higher apparent specificity of dCas12a systems compared to Sp-dCas9 (based on binding kinetics[45,46] and specificity[46–48]) and adjusted for the apparent different effects of mismatches on the strength of binding to DNA.[46,49] For Lb-dCas12a, the same adjustments were made as those for Fn-dCas12a, except with different non-canonical PAMs (CTTV, TTTT, TTCV, TCTV)[43] and slightly lower penalty values for mismatches in the gRNA spacer sequence due to higher apparent binding affinity to DNA compared to Fn-dCas12a.[49]

The Python scripts can create genome-wide CRISPRi libraries in any host organism with an annotated genome via the additional input parameters described above. If indicated and a BLAST results file provided, genes are first clustered based on homology determined from the BLAST results file. It is important to note that if gene clustering is not enabled and a set of genes share significant homology (e.g. tRNA genes), then gRNA may not be designed to target the homologous genes due to detection of off-target effects. gRNAs are designed for each single target gene as described above. For clusters of homologous genes, gRNAs are designed for a representative gene from the cluster. After designing all gRNA targeting the specified genes in the genome, the script designs non-targeting control (NC) gRNA. Random spacer sequences are generated and aligned against the genome using SeqMap.[41] If the gRNA does not have an alignment with the genome, it is screened for the specified GC content and disallowed DNA sequences. NC gRNA that pass these criteria are chosen up to the specified number of NC gRNA. If selected, repeated gRNA spacers in the library (which can occur for overlapping genes) are removed. Finally, if given, constant upstream and downstream sequences are appended to each designed spacer and the resulting oligo sequence library provided in CSV format for ordering. A flowchart for the design of gRNA using the set of scripts is given in **Figure S33**. The scripts are available at: [https://github.com/AndrewsLabSynBio/CRISPRi_gRNA_library_design/tree/main].

(15) Qi, L. S.; Larson, M. H.; Gilbert, L. A.; Doudna, J. A.; Weissman, J. S.; Arkin, A. P.; Lim, W. A. Repurposing CRISPR as an RNA-Guided Platform for Sequence-Specific Control of

Gene Expression. *Cell* **2013**, *152* (5), 1173–1183. https://doi.org/10.1016/j.cell.2013.02.022.
(16) Liao, C.; Slotkowski, R. A.; Achmedov, T.; Beisel, C. L. The Francisella Novicida Cas12a Is Sensitive to the Structure Downstream of the Terminal Repeat in CRISPR Arrays. *RNA Biology* **2019**, *16* (4), 404–412. https://doi.org/10.1080/15476286.2018.1526537.
(17) Mekler, V.; Minakhin, L.; Semenova, E.; Kuznedelov, K.; Severinov, K. Kinetics of the CRISPR-Cas9 Effector Complex Assembly and the Role of 3′-Terminal Segment of Guide RNA. *Nucleic Acids Res* **2016**, *44* (6), 2837–2845. https://doi.org/10.1093/nar/gkw138.
(18) Zetsche, B.; Gootenberg, J. S.; Abudayyeh, O. O.; Slaymaker, I. M.; Makarova, K. S.; Essletzbichler, P.; Volz, S. E.; Joung, J.; van der Oost, J.; Regev, A.; Koonin, E. V.; Zhang, F. Cpf1 Is a Single RNA-Guided Endonuclease of a Class 2 CRISPR-Cas System. *Cell* **2015**, *163* (3), 759–771. https://doi.org/10.1016/j.cell.2015.09.038.
(19) Pang, Y. Y.; Schwartz, J.; Thoendel, M.; Ackermann, L. W.; Horswill, A. R.; Nauseef, W. M. Agr-Dependent Interactions of Staphylococcus Aureus USA300 with Human Polymorphonuclear Neutrophils. *Journal of Innate Immunity* **2010**, *2* (6), 546–559. https://doi.org/10.1159/000319855.
(20) Cormack, B. P.; Valdivia, R. H.; Falkow, S. FACS-Optimized Mutants of the Green Fluorescent Protein (GFP). *Gene* **1996**, *173* (1), 33–38. https://doi.org/10.1016/0378-1119(95)00685-0.
(21) Rondthaler, S. N.; Sarker, B.; Howitz, N.; Shah, I.; Andrews, L. B. Toolbox of Characterized Genetic Parts for Staphylococcus Aureus. *ACS Synthetic Biology* **2024**, *13* (1), 103–118. https://doi.org/10.1021/acssynbio.3c00325.
(22) Moriel, D. G.; Tan, L.; Goh, K. G. K.; Phan, M.-D.; Ipe, D. S.; Lo, A. W.; Peters, K. M.; Ulett, G. C.; Beatson, S. A.; Schembri, M. A. A Novel Protective Vaccine Antigen from the Core Escherichia Coli Genome. *mSphere* **2016**, *1* (6), 10.1128/msphere.00326-16. https://doi.org/10.1128/msphere.00326-16.
(23) Sayers, E. W.; Bolton, E. E.; Brister, J. R.; Canese, K.; Chan, J.; Comeau, D. C.; Connor, R.; Funk, K.; Kelly, C.; Kim, S.; Madej, T.; Marchler-Bauer, A.; Lanczycki, C.; Lathrop, S.; Lu, Z.; Thibaud-Nissen, F.; Murphy, T.; Phan, L.; Skripchenko, Y.; Tse, T.; Wang, J.; Williams, R.; Trawick, B. W.; Pruitt, K. D.; Sherry, S. T. Database Resources of the National Center for Biotechnology Information. *Nucleic Acids Research* **2022**, *50* (D1), D20–D26. https://doi.org/10.1093/nar/gkab1112.
(24) Camacho, C.; Coulouris, G.; Avagyan, V.; Ma, N.; Papadopoulos, J.; Bealer, K.; Madden, T. L. BLAST+: Architecture and Applications. *BMC Bioinformatics* **2009**, *10* (1), 421. https://doi.org/10.1186/1471-2105-10-421.
(25) Katoh, K.; Standley, D. M. MAFFT Multiple Sequence Alignment Software Version 7: Improvements in Performance and Usability. *Molecular Biology and Evolution* **2013**, *30* (4), 772–780. https://doi.org/10.1093/molbev/mst010.
(26) R Core Team. R: A Language and Environment for Statistical Computing, 2023. https://www.R-project.org/.
(27) Paradis, E.; Schliep, K. Ape 5.0: An Environment for Modern Phylogenetics and Evolutionary Analyses in R. *Bioinformatics* **2019**, *35* (3), 526–528. https://doi.org/10.1093/bioinformatics/bty633.
(28) Schliep, K. P. Phangorn: Phylogenetic Analysis in R. *Bioinformatics* **2011**, *27* (4), 592–593. https://doi.org/10.1093/bioinformatics/btq706.
(29) Wang, L.-G.; Lam, T. T.-Y.; Xu, S.; Dai, Z.; Zhou, L.; Feng, T.; Guo, P.; Dunn, C. W.; Jones, B. R.; Bradley, T.; Zhu, H.; Guan, Y.; Jiang, Y.; Yu, G. Treeio: An R Package for Phylogenetic Tree Input and Output with Richly Annotated and Associated Data. *Molecular Biology and Evolution* **2020**, *37* (2), 599–603. https://doi.org/10.1093/molbev/msz240.
(30) Guangchuang, Y. *Data Integration, Manipulation and Visualization of Phylogenetic Trees*, 1st edition.; Chapman and Hall/CRC, 2022.

(31) Xu, S.; Li, L.; Luo, X.; Chen, M.; Tang, W.; Zhan, L.; Dai, Z.; Lam, T. T.; Guan, Y.; Yu, G. Ggtree: A Serialized Data Object for Visualization of a Phylogenetic Tree and Annotation Data. *iMeta* **2022**, *1* (4), e56. https://doi.org/10.1002/imt2.56.
(32) Ang, J.; Harris, E.; Hussey, B. J.; Kil, R.; McMillen, D. R. Tuning Response Curves for Synthetic Biology. *ACS Synthetic Biology* **2013**, *2* (10), 547–567. https://doi.org/10.1021/sb4000564.
(33) Virtanen, P.; Gommers, R.; Oliphant, T. E.; Haberland, M.; Reddy, T.; Cournapeau, D.; Burovski, E.; Peterson, P.; Weckesser, W.; Bright, J.; van der Walt, S. J.; Brett, M.; Wilson, J.; Millman, K. J.; Mayorov, N.; Nelson, A. R. J.; Jones, E.; Kern, R.; Larson, E.; Carey, C. J.; Polat, İ.; Feng, Y.; Moore, E. W.; VanderPlas, J.; Laxalde, D.; Perktold, J.; Cimrman, R.; Henriksen, I.; Quintero, E. A.; Harris, C. R.; Archibald, A. M.; Ribeiro, A. H.; Pedregosa, F.; van Mulbregt, P. SciPy 1.0: Fundamental Algorithms for Scientific Computing in Python. *Nature Methods* **2020**, *17* (3), 261–272. https://doi.org/10.1038/s41592-019-0686-2.
(34) Liao, C.; Slotkowski, R. A.; Achmedov, T.; Beisel, C. L. The Francisella Novicida Cas12a Is Sensitive to the Structure Downstream of the Terminal Repeat in CRISPR Arrays. *RNA Biol* **2018**, *16* (4), 404–412. https://doi.org/10.1080/15476286.2018.1526537.
(35) Chen, P.-Y.; Qian, Y.; Del Vecchio, D. A Model for Resource Competition in CRISPR-Mediated Gene Repression. In *2018 IEEE Conference on Decision and Control (CDC)*; 2018; pp 4333–4338. https://doi.org/10.1109/CDC.2018.8619016.
(36) Yao, L.; Cengic, I.; Anfelt, J.; Hudson, E. P. Multiple Gene Repression in Cyanobacteria Using CRISPRi. *ACS Synthetic Biology* **2016**, *5* (3), 207–212. https://doi.org/10.1021/acssynbio.5b00264.
(37) Gordon, G. C.; Korosh, T. C.; Cameron, J. C.; Markley, A. L.; Begemann, M. B.; Pfleger, B. F. CRISPR Interference as a Titratable, Trans-Acting Regulatory Tool for Metabolic Engineering in the Cyanobacterium Synechococcus Sp. Strain PCC 7002. *Metabolic Engineering* **2016**, *38*, 170–179. https://doi.org/10.1016/j.ymben.2016.07.007.
(38) Lee, H. H.; Ostrov, N.; Wong, B. G.; Gold, M. A.; Khalil, A. S.; Church, G. M. Functional Genomics of the Rapidly Replicating Bacterium Vibrio Natriegens by CRISPRi. *Nature Microbiology* **2019**, *4* (7), 1105–1113. https://doi.org/10.1038/s41564-019-0423-8.
(39) Fontana, J.; Dong, C.; Ham, J. Y.; Zalatan, J. G.; Carothers, J. M. Regulated Expression of sgRNAs Tunes CRISPRi in E. Coli. *Biotechnology Journal* **2018**, *13* (9), 1800069. https://doi.org/10.1002/biot.201800069.
(40) Wang, T.; Guan, C.; Guo, J.; Liu, B.; Wu, Y.; Xie, Z.; Zhang, C.; Xing, X.-H. Pooled CRISPR Interference Screening Enables Genome-Scale Functional Genomics Study in Bacteria with Superior Performance. *Nature Communications* **2018**, *9* (1), 2475. https://doi.org/10.1038/s41467-018-04899-x.
(41) Jiang, H.; Wong, W. H. SeqMap: Mapping Massive Amount of Oligonucleotides to the Genome. *Bioinformatics* **2008**, *24* (20), 2395–2396. https://doi.org/10.1093/bioinformatics/btn429.
(42) Hsu, P. D.; Scott, D. A.; Weinstein, J. A.; Ran, F. A.; Konermann, S.; Agarwala, V.; Li, Y.; Fine, E. J.; Wu, X.; Shalem, O.; Cradick, T. J.; Marraffini, L. A.; Bao, G.; Zhang, F. DNA Targeting Specificity of RNA-Guided Cas9 Nucleases. *Nature Biotechnology* **2013**, *31* (9), 827–832. https://doi.org/10.1038/nbt.2647.
(43) Miao, C.; Zhao, H.; Qian, L.; Lou, C. Systematically Investigating the Key Features of the DNase Deactivated Cpf1 for Tunable Transcription Regulation in Prokaryotic Cells. *Synthetic and Systems Biotechnology* **2019**, *4* (1), 1–9. https://doi.org/10.1016/j.synbio.2018.11.002.
(44) Specht, D. A.; Xu, Y.; Lambert, G. Massively Parallel CRISPRi Assays Reveal Concealed Thermodynamic Determinants of dCas12a Binding. *Proceedings of the National Academy of Sciences* **2020**, *117* (21), 11274–11282. https://doi.org/10.1073/pnas.1918685117.
